# Brain-wide neural activity underlying memory-guided movement

**DOI:** 10.1101/2023.03.01.530520

**Authors:** Susu Chen, Yi Liu, Ziyue Wang, Jennifer Colonell, Liu D. Liu, Han Hou, Nai-Wen Tien, Tim Wang, Timothy Harris, Shaul Druckmann, Nuo Li, Karel Svoboda

## Abstract

Behavior requires neural activity across the brain, but most experiments probe neurons in a single area at a time. Here we used multiple Neuropixels probes to record neural activity simultaneously in brain-wide circuits, in mice performing a memory-guided directional licking task. We targeted brain areas that form multi-regional loops with anterior lateral motor cortex (ALM), a key circuit node mediating the behavior. Neurons encoding sensory stimuli, choice, and actions were distributed across the brain. However, in addition to ALM, coding of choice was concentrated in subcortical areas receiving input from ALM, in an ALM-dependent manner. Choice signals were first detected in ALM and the midbrain, followed by the thalamus, and other brain areas. At the time of movement initiation, choice-selective activity collapsed across the brain, followed by new activity patterns driving specific actions. Our experiments provide the foundation for neural circuit models of decision-making and movement initiation.

## Introduction

Behaviors are mediated by neural dynamics that are distributed and coordinated across multiple brain areas. However, neural correlates of behavior have traditionally been studied in small subsets of neurons in one brain area at a time. This is akin to listening to a symphony played by one musician in isolation, which would provide only a pale and incomplete reflection of the complete piece of music. Moreover, different brain areas are typically investigated in the context of specific behaviors and with diverse recording methods, making it difficult to integrate data across studies into a more complete view of the neural dynamics underlying any one behavior.

Recently, new types of silicon array electrodes (J. J. Jun 2017; Steinmetz et al. 2021) have made recordings from multiple brain areas routine (Allen et al. 2019; Steinmetz et al. 2019; Siegle et al. 2021). Few studies have measured behavior-related activity with reference to multi-regional connectivity, and at most two brain areas at a time (Peters et al. 2021). Here we study neural activity across dozens of brain areas in a consistent behavioral task, guided by anatomical connectivity.

During decision-making, neural activity encodes sensory stimuli and later behavioral choices and actions. Choice-related activity, also sometimes referred to as ‘preparatory activity’ (Tanji and Evarts 1976; Z. V. Guo et al. 2014; Erlich, Bialek, and Brody 2011; Churchland et al. 2010), predicts upcoming actions, often seconds before they are initiated (Shibasaki and Hallett 2006). Choice-related activity has been observed in multiple brain areas, including cortical and subcortical brain areas in primates (Britten et al. 1996; S. Liu et al. 2013; Fried, Mukamel, and Kreiman 2011) and rodents (Z. V. Guo et al. 2014; Erlich, Bialek, and Brody 2011; G. Chen et al. 2021; Z. V. Guo et al. 2017; Chunyu A. Duan, Pan, et al. 2021; Gao et al. 2018). The sequence of neural events linking sensory stimuli and choice has not been mapped at brain-wide scales at the level of individual neurons and with temporal resolutions relevant to neural computation.

The mouse anterior lateral motor cortex (ALM) is a critical node for both the planning and execution of directional licking (Z. V. Guo et al. 2014; Economo et al. 2018; Inagaki, Chen, Ridder, et al. 2022; N. Li et al. 2015; Inagaki et al. 2018; Chunyu A. Duan, Pan, et al. 2021). During decision-making, individual neurons in ALM exhibit choice-related activity in anticipation of specific actions (Z. V. Guo et al. 2014). ALM population activity predicts specific licking direction, and sculpting ALM activity before the initiation modulates future actions in a predictable manner (N. Li et al. 2016; Inagaki et al. 2019; Daie, Svoboda, and Druckmann 2021; Finkelstein et al. 2021). Similar to other cortical areas, ALM activity is also modulated by animal movements, including movements that are not explicitly related to the behavioral task (Musall et al. 2019; Huber et al. 2012; Stringer et al. 2019; Niell and Stryker 2010; Salkoff et al. 2020).

ALM forms multi-regional loops with subcortical brain areas that play roles in action selection and execution (Inagaki, Chen, Daie, et al. 2022; Z. V. Guo et al. 2017; N. Li et al. 2015; Chunyu A. Duan, Pan, et al. 2021; Gao et al. 2018). Previous studies combining electrophysiological recordings and optogenetic manipulations have shown that choice-selective activity is maintained in neurons distributed across multiple brain areas. For example, persistent choice-selective activity cannot be sustained by ALM alone, but requires reverberant excitation with higher-order thalamus that is reciprocally connected with ALM (Z. V. Guo et al. 2017). Additionally, choice-related activity has been observed in the cerebellum (Gao et al. 2018) and basal ganglia (Y. Wang et al. 2021; Tang et al. 2021), specifically in regions that receive indirect input from ALM and relay inputs back to ALM through the thalamus. Midbrain reticular and pedunculopontine nuclei (MRN/PPN) route phasic signals through the motor thalamus to ALM, triggering a reorganization of cortical state from choice-related activity to a motor command, which in turn drives appropriate licking (Inagaki, Chen, Ridder, et al. 2022).

Determining how choice-related activity evolves across the brain during decision-making, and how choice signals relate to signals encoding animal movements, requires measurements of neural activity across the brain in a standardized behavioral task, and ideally across multiple connected brain areas simultaneously. Determining if choice-related activity can be maintained in subcortical structures independent of ALM input requires measurements of neural activity together with targeted inactivation. Answers to these questions are fundamental to neural circuit models of decision-making, action selection, and movement initiation. Here we address these questions by simultaneously recording from multiple brain areas while mice performed a delayed-response decision-making task. Recordings were guided by mesoscopic connectivity and focused on structures that are part of multi-regional loops involving ALM, including the thalamus, basal ganglia, midbrain, medulla, pons, and cerebellum. In a subset of experiments neural activity was inactivated in ALM.

Neural activity related to stimulus, choice and movement was distributed across the forebrain, midbrain and hindbrain. However, choice-related activity was concentrated in subcortical brain areas that are connected to ALM. Moreover, maintenance of choice-related activity in these areas required input from ALM. Choice-selective neural activity was detected first in ALM, followed by the midbrain, thalamus, and other structures, including the striatum and hindbrain. During the delay epoch, choice-related activity was strongly correlated on a trial-by-trial manner across ALM and its target areas. At the time of movement initiation, choice-selective activity collapsed across the brain, followed by new modes of movement-selective activity. Our studies provide the foundation for understanding decision-making, action selection, and movement initiation on a brain-wide scale.

## Results

### Anatomy-guided recordings of multi-regional neural activity

Mice (n = 28, Table 1) were trained in a memory-guided movement task (Z. V. Guo et al. 2014; Inagaki et al. 2018; Chunyu A. Duan, Pan, et al. 2021) (**Figure 1A**). During a sample epoch, mice distinguished the pitch of one of two possible auditory stimuli (3 kHz or 12 kHz; three 150 ms beeps). After a 1.2 seconds long delay (memory) epoch, an auditory ‘Go’ cue (6 kHz, 0.1 s) initiated the response epoch. Mice reported their choice with directional licking according to the auditory instruction. Correct responses (high tone → lick left; low tone → lick right) triggered a water reward, whereas incorrect responses caused a mild punishment (a timeout). During the delay epoch, mice maintained a memory of their choice and planned a movement. Mice performed on average 476 (Mean; range, 130-785) trials per session with 84% correct rate (range, 65-99%; Methods).

**Figure 1.**
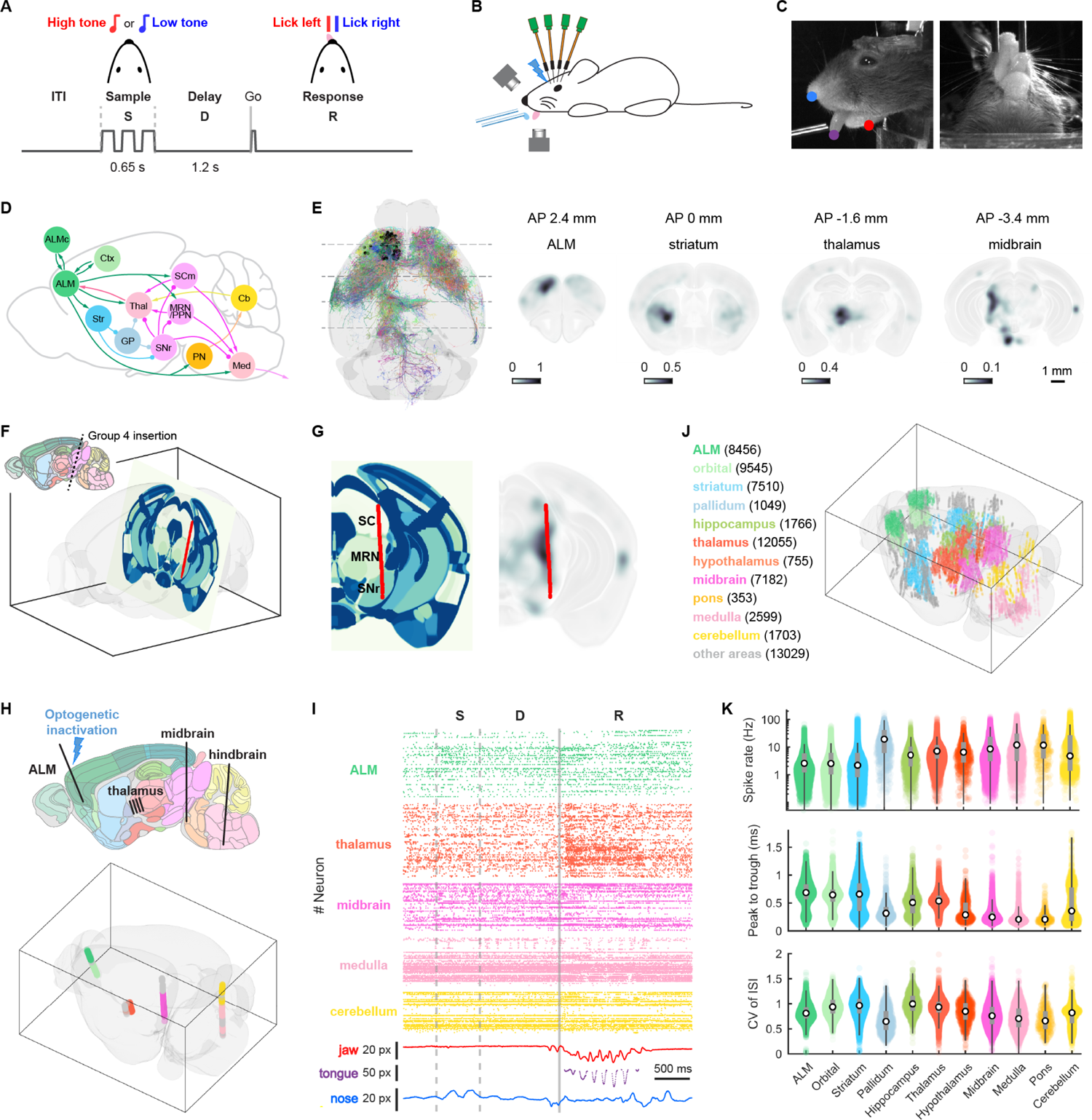
Brain-wide, anatomy-guided recordings of neural activity. A. Memory-guided directional licking task. Mice were trained to lick to one of two possible lickports instructed by one of two auditory stimuli (high or low tone), presented during the sample epoch (S). Mice respond after a ‘Go’ cue in the response epoch (R), after a delay epoch (D). Between trials is the inter-trial interval (ITI). B. Typical recording with four Neuropixels probes during behavior. Electrophysiological measurements were combined with movement tracking using high-speed videography and laser-based photoinhibition of ALM (lightning bolt). C. Video images, showing three tracked keypoints (nose, tongue, jaw) in the side view. D. Major brain areas connected to ALM in multi-regional networks, including the isocortex (ALMc - contralateral ALM, Ctx - cortex; green), Striatum (Str; cyan), Globus Pallidus/pallidum (GP; light blue), Thalamus (Thal, red), Midbrain (SCm - motor area of superior colliculus, MRN - midbrain reticular nucleus, SNr - substantia nigra pars reticulata; magenta), Pontine nucleus (PN; orange), Cerebellum (Cb; yellow), and Medulla (pink). Arrowheads, excitatory connections; circles, inhibitory connections. E. Multi-regional connectivity from single neuron reconstructions. Left, horizontal view of 212 reconstructed ALM neurons used to define ALM projection zones (ml-neuronbrowser.janelia.org). Black dots, soma locations. Neurons in the right hemisphere were mirrored to the left hemisphere. Gray dashed lines, coronal slices shown on the right. Right, cortical projection density on four example coronal sections centered on ALM, striatum, thalamus and midbrain, approximately corresponding to the targets of four insertion groups (Group 1-4, see text). AP, anterior-posterior location relative to bregma. Voxel intensity is the axonal protection length-density smoothed with a 3D Gaussian (sigma = 150 µm). Projection density has been normalized to the range of 0-1 across the brain. F. Example insertion covering multiple nuclei in midbrain (Group 4). The insertion lies in an oblique, near-coronal section. Red line indicates the planned electrode track. G. Oblique section displayed in the Allen Reference Atlas (ARA) and the corresponding ALM projection density map in CCF. Planned electrode track (red) spans ALM projection regions, including superior colliculus (SC), midbrain reticular nucleus (MRN). In addition, the insertion also targets the substantia nigra pars reticulata (SNr), which is downstream of the striatum. H. Example experiment with four Neuropixels probes. Single-shank probes target left frontal cortex, left midbrain, and right medulla; one four-shank probe targets left thalamus (corresponding to Group 1, 4, 5, 3, respectively; see text). Bottom, recording locations registered to the CCF. Neurons are color-coded by brain area based on the ARA color scheme (Wang and Ding et al, 2020). I. Population spike raster for one trial corresponding to the recording in H. Sample epoch is between the dashed gray lines, and the solid gray line depicts the ‘Go’ cue. Bottom traces show movement of fiducial points as in panel C, in pixels (px). J. Summary of the data set after stringent quality control, by brain area, aggregated over 660 penetrations, 173 behavioral sessions, and 28 mice. K. Summary of spike rate, spike width (peak to trough time), and coefficient of variation (CV) of the interspike interval (ISI).

To examine how task-related information is represented across multi-regional circuits, we performed recordings with up to five Neuropixels probes simultaneously (two probes, 2 sessions; three probes, 53 sessions; four probes, 98 sessions; five probes, 20 sessions) (**Figure 1B, H, Figure S1A, B**, Table 2). In a subset of randomly selected trials (typically 25%) one of the ALM hemispheres was transiently inactivated by optogenetic activation of GABAergic interneurons expressing ChR2 (i.e. VGAT-ChR2 mice, N = 25) (Z. V. Guo et al. 2014; Inagaki et al. 2018; Nuo Li et al. 2019; Zhao et al. 2011) late (final 0.5 s, N = 17) in the delay epoch (**Figure 1B, H**). Animal movements were recorded with multi-view, high-speed videography (300 Hz) (**Figure 1C**). Keypoints defining specific orofacial movements were tracked offline (Mathis et al. 2018)(**Figure 1C, I, Figure S1C**).

We focused our recordings on ALM, areas that receive ALM projections, and several areas downstream of these ALM projection zones, including pallidum and substantia nigra pars reticulata (SNr). These areas together form multi-regional loops (Inagaki, Chen, Daie, et al. 2022) (**Figure 1D, E, Figure S1D, E**). We defined ALM projection zones using 212 (see Methods) reconstructed single ALM neurons (http://ml-neuronbrowser.janelia.org), including 59 intratelencephalic (IT) neurons, 113 pyramidal tract (PT) neurons, and 40 corticothalamic (CT) neurons (Winnubst et al. 2019) (**Figure 1E**). The resulting projection map is in qualitative agreement with fluorescence-based measurements of mesoscale projections (Oh et al. 2014; Zingg et al. 2014) (**Figure S1D, E**), but the single cell reconstructions provide a more quantitative estimate of the innervation density (Methods). The strength of an ALM projection was defined as the axonal density (smoothed with a 3D Gaussian filter; sigma, 150 µm) in a CCF location (Allen Mouse Common Coordinate Framework, CCFv3) (Q. Wang et al. 2020) (**Figure 1E, Figure S1D**).

Using ALM projection zones and additional anatomical data (Oh et al. 2014) as guidance, Neuropixels probe insertions were targeted to the following groups of brain areas: Group 1, ALM and underlying cortical regions (e.g. orbital frontal cortex); Group 2, striatum; Group 3, higher-order thalamus; Group 4, midbrain; Group 5, medulla and the overlying cerebellum; Group 6, other regions (Table 2). When possible, we angled individual Neuropixels probes to sample neural activity from multiple relevant brain areas (**Figure 1H, I**). For example, Group 4 includes recordings from motor-related parts of the superior colliculus (SCm), MRN, and substantia nigra pars reticulata (SNr) (**Figure 1F, G**).

Spike sorting was performed using Kilosort2 (KS2) (Stringer et al. 2019) (Methods). After stringent quality control, we obtained approximately 66,000 single units (**Figure 1J, Figure S1F**; Methods) from KS2 sorting. The CCF locations of recorded neurons were identified using a combination of histological information and electrophysiological landmarks (L. D. Liu et al. 2020). Each recording session yielded simultaneous measurements from hundreds of neurons (median = 376) and often more than a dozen brain areas (**Figure 1H, I, Figure S1C, F**). Neurons exhibited a wide range of spike rates (**Figure 1K**), waveform features (such as spike width, defined as the time between peak and trough), and spiking variability (coefficient of variation of inter-spike-interval), with clear differences across brain areas.

### Neural coding of sensory stimulus, choice, action and outcome

Each behavioral trial was associated with a set of task variables, including sensory stimulus (high vs. low tone), choice (decision to lick left vs. right, before action), action (behavioral response after the ‘Go’ cue), and outcome (rewarded vs. unrewarded, after the action) (**Figure 2A**). We estimated the selectivity of single neurons by computing the spike rate difference across the values of the task variables during different epochs, using both correct and error trials (Methods). Individual neurons were selective for the sensory stimulus (**Figure 2B**), choice (measured during the delay epoch) (**Figure 2C**), action (measured during the response epoch) (**Figure 2D**), and/or outcome (measured after licking offset) (**Figure 2E, Figure S2E**). Many neurons encoded multiple task variables. For example, an ALM neuron could be selective for choice during the delay epoch and selective for outcome late in the response epoch (T. W. Chen et al. 2017).

**Figure 2.**
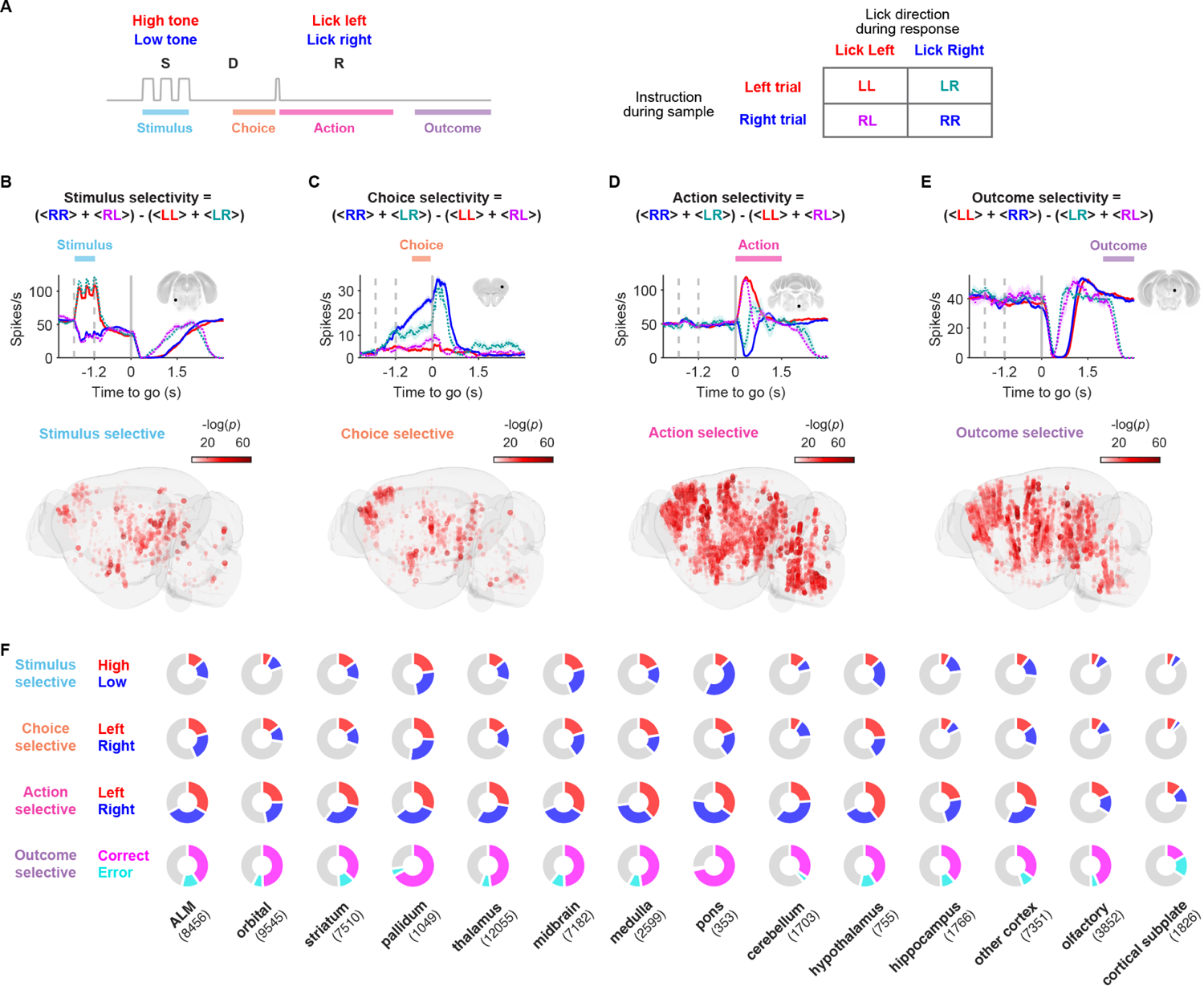
Widely distributed selective neurons. A. Time windows for calculating single neuron selectivity: 0.6 s overlapping with the sensory stimulus the sample epoch; last 0.6 s of the delay epoch to isolate choice selectivity; first 1.5 s post ‘Go’ cue for action selectivity; 2-3 s post ‘Go’ cue, when licking had terminated (Fig. S2E), for outcome selectivity. Trials were divided into four types based on stimulus (instruction, L or R) and choice (lick direction, L or R). Trials are labeled as (LL, RR, LR, RL). LL and RR are correct trials. LR and RL are error trials. B. Top, stimulus selectivity is defined as the mean spike rate difference, grouped based on the sensory stimulus (stimulus R - stimulus L). Middle, peri-stimulus time histogram (PSTH) of an example neuron (recorded in pons) selective for the auditory stimulus. Solid lines, correct trials; dashed lines, error trials. Color code as defined in A. Bottom, stimulus selective neurons; circle size and hue are proportional to -log(p). C. Same as B, but for choice selectivity, grouped based on upcoming licking direction (Choice R - Choice L). Example choice-selective neuron recorded in ALM. D. Same as B, but for action selectivity, grouped based on licking direction (Lick R - Lick L). Example response-selective neuron recorded in the medulla. E. Same as B, but for outcome selectivity, grouped based on correct response (Reward - Unrewarded) with an example neuron recorded in the midbrain. F. Proportions of selective neurons across major brain areas. Numbers in parentheses are neurons in each brain area.

We quantified the proportion of units selective for stimulus, choice, action, and outcome (rank sum test, P < 0.05; **Figure 2F, Figure S2A**). For each task variable, selective neurons were widely distributed, albeit with different incidences and strengths across brain areas (**Figure 2B-E**). ALM showed a higher proportion of choice-selective neurons during the delay epoch compared to its targets, including the striatum, thalamus, midbrain, and medulla (**Figure 2C,F, Figure S2A**). Interestingly, the pallidum also showed a high proportion of choice-selective activity (**Figure 2F, Figure S2A**), likely inherited from the striatum and/or thalamus. Brainstem regions, including the medulla and pons, contained the largest proportion of neurons encoding the action (**Figure 2D,F**). Neurons in the reticular formation of the medulla showed selectivity for ipsi-versive movements (**Figure S3B**), consistent with recordings from premotor neurons that control tongue movements into ipsilateral space (N. Li et al. 2015; Takatoh et al. 2021; Bennett and Ramsay 1941). Encoding of action and outcome was more global than encoding of sensory stimulus and choice (**Figure 2B-F, Figure S2C,D**). In regions outside of the ALM network, such as hippocampus and olfactory areas, we found little choice-related activity (**Figure 2F, Figure S2A**).

Averaging across selective neurons revealed the dynamics of representation of information across brain areas (**Figure S2B**). Auditory stimulus information peaked during the sample epoch after tone onset. Choice selectivity arose in the early sample epoch, ramped up throughout the delay epoch, and reached a maximum at the beginning of the response epoch. As expected, outcome selectivity was limited to the response epoch, after the animal had sampled one of the lick ports (and was measured after reward-related licking ceased).

### Neural coding of movement

In most laboratory tasks, decisions are reported by planning and executing movements (Selen, Shadlen, and Wolpert 2012; Carland, Thura, and Cisek 2019; Wilson 2002). Beyond these instructed movements, head-restrained mice can adjust their posture and make micro-movements as decisions are made. In addition, mice fidget and perform other movements that may be unrelated to the task (Huber et al. 2012; Musall et al. 2019; Stringer et al. 2019; Kahneman and Beatty 1966). These movements can differ across behavioral sessions and individual mice, and can be time-locked to task events and thus correlate with task-related activity. For example, subsets of neurons in the striatum, thalamus, midbrain, and medulla exhibited oscillatory spiking in phase with tongue protrusions as well as anticipatory rhythmic jaw movements (**Figure S3A**).

We quantified the neural encoding of movement by creating a statistical model that predicts single trial, time-varying neural activity (spike rate) of individual neurons from behavioral video. We use fraction of variance explained (R^2^) as a proxy for the strength of neural encoding of movement.

We used unsupervised convolutional autoencoders (CAE) to extract a low-dimensional (16-dimension embedding vectors) representation of the video (**Figure 3A**; Methods) (Batty et al. 2019). Alternatively, we tracked facial features (including the jaw, tongue, and nose) using ‘keypoint’ (part) tracking (Mathis et al. 2018) (**Figure 1C,I, Figure S1C**). We used both the embedding and tracked keypoints to predict spike rates using regularized linear regression (40 ms bin width for estimating spike rates; **Figure 3B-F**). Predictions based on embedding vectors outperformed those based on keypoint tracking (improvement in R^2^, mean = 23.3 ± 0.3% S.E.M., N = 23,044 neurons, two-sample t test, P < 0.001, **Figure S3C;** Methods), but at the expense of interpretability. Analyses provided similar results across both methods and multiple bin widths for estimation of spike rates (20, 40, 80, 160, 320 ms, **Figure S3D**).

**Figure 3.**
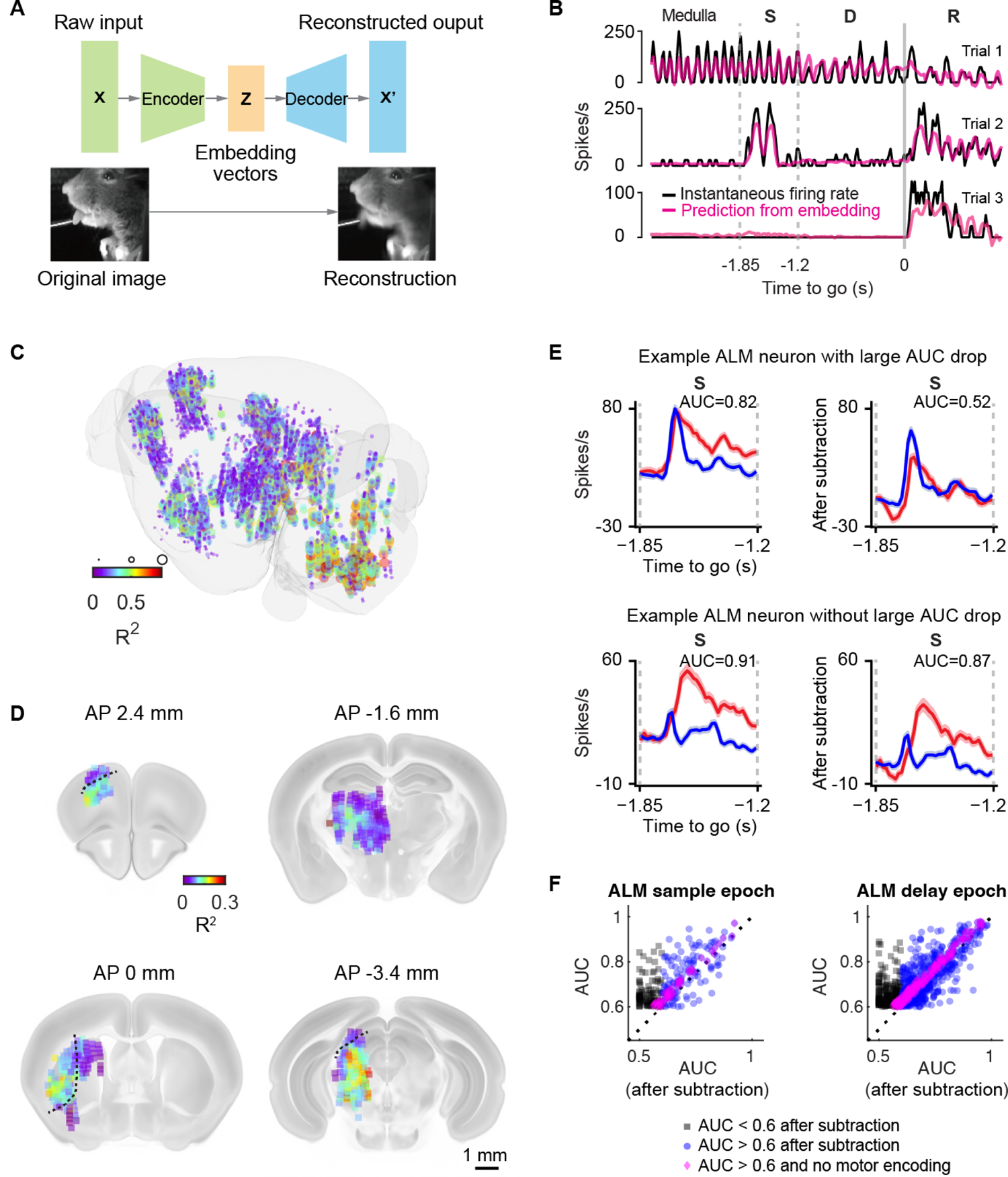
Neural coding of movement. A. Embedding-based analysis of behavioral videos. High-speed videos were compressed using a convolutional autoencoder, producing a low-dimensional representation of behavior (here a 16-dimensional embedding vector for each video frame). The embedding vectors were used to predict neural activity of individual neurons (B). B. Encoding of movements in an example neuron (medulla, three trials). Spike rate (black) and overlaid prediction of spike rate from embedding vectors (magenta). C. Spatial map of movement encoding. Each circle corresponds to a single neuron. Color and dot size represents explained fractional variance (R^2^). D. Map of movement encoding, projected onto coronal sections. The color of each voxel represents mean R^2^ in a 300 µm cube. The highlighted brain areas are (clockwise from upper left): ALM, thalamus, striatum, and midbrain. The dotted lines represent borders for regions of interest analysis (see main text). E. Effect of subtracting video-based prediction on choice coding (AUC) during the sample epoch. Left, before subtraction; right, after subtraction. Top, example ALM neurons with large AUC drop after subtraction, i.e. high movement encoding. Bottom, ALM neurons with small AUC drop. Lines, trial-averaged spike rates (red, lick-left; blue, lick-right). Shaded area, S.E.M. F. Effect of subtracting video-based prediction on decodability of choice during the sample epoch (left) and delay epoch (right). Each dot corresponds to a neuron. Only neurons with AUC > 0.6 before subtraction are shown.

The data revealed a gradient of movement encoding across the brain, with the highest encoding closest to the periphery (**Figure 3C, D, Figure S3C**). Among the brain areas that have direct projections from ALM, movement encoding was highest in the medulla, where multiple premotor and motor nuclei for orofacial movements reside (McElvain et al. 2018), in addition to brainstem relays for proprioceptive and reafference sensory input (mean R^2^: medulla = 0.216 ± 0.06 S.E.M., n = 36 insertions; midbrain = 0.120 ± 0.006, n = 81, P < 0.001, see Methods). The midbrain, which also contains premotor neurons and receives sensory input directly from the brainstem, showed the next strongest encoding of movements, followed by ALM/striatum/thalamus (**Figure S3C, D**).

Encoding of movement was structured within brain areas. For instance, in ALM, deeper cortical layers exhibited stronger encoding compared to superficial layers (R^2^ deep layers mean = 0.07 ± 0.001 S.E.M., N = 7084 neurons, R^2^ superficial layers mean = 0.04 ± 0.003 S.E.M., N = 487 neurons, Kolmogorov-Smirnov (KS) test, P < 1e-5, **Figure 3D**). This is consistent with the laminar organization of motor cortex: superficial motor cortex neurons receive primarily sensory cortex input, whereas deeper layer neurons project to motor centers and receive input from motor thalamus (Economo et al. 2018; Hooks et al. 2013; Kaneko et al. 2000).

The striatum showed strong movement encoding in the dorsal-lateral part (caudoputamen) but not in other sectors (Lee and Sabatini 2021) (mean R^2^: dorsal-lateral = 0.11 ± 0.003 S.E.M., N = 2987 neurons; medial = 0.08 ± 0.003 S.E.M., N = 1158 neurons, KS test, P < 1e-5, **Figure 3D**). In superior colliculus (SC), movement encoding was strong in the intermediate and deep (motor) layers which project to the medulla, but not in the superficial (sensory) layers (Rossi et al. 2016) (mean R^2^: deep mean = 0.15 ± 0.003 S.E.M., N = 1433 neurons; superficial layers = 0.04 ± 0.007 S.E.M., N = 73 neurons, KS test, P < 1e-5, **Figure 3D**).

We analyzed movement encoding in ALM. Among selective ALM neurons (quantified as area-under-the-curve, AUC > 0.6), a subset of neurons lost selectivity after subtracting neural activity predicted from movement, whereas other neurons maintained selectivity (**Figure 3E**). For many ALM neurons with robust coding for choice (AUC > 0.6) fractional R^2^ predicted from movement was not significantly different from zero (**Figure 3F).** For example, in the delay epoch, 801 neurons had AUC greater than 0.6, but of these 283 had negligible prediction (R^2^ < 0.02). For other ALM neurons, AUC dropped after movement-prediction subtraction, with 157 neurons dropping below the criterion value.

We note that correlation with movement does not imply that neural dynamics are not ‘cognitive’. After all, neural activity encoding ‘choice’ in ALM is expected to modulate movement and posture through ALM projections to motor centers, and in turn can be shaped by reafference input.

### Encoding of choice in ALM projection zones

Neurons encoding task-related variables were distributed across all of the recorded brain areas (**Figure 2**). However, the distribution of selective neurons is not uniform. For example, previous imaging experiments in the motor cortex have shown that neurons with choice-related activity during the delay epoch are concentrated in ALM, compared to neighboring parts of the motor cortex (T. W. Chen et al. 2017).

We evaluated the possibility that, similar to ALM, choice-related activity may be concentrated in specific sub-regions across subcortical brain areas. We assessed the spatial distribution of selectivity with respect to ALM projections (**Figure 4; Figure S4**). In each subcortical brain area we quantified the proportion of selective neurons in a brain area as a function of the local ALM projection strength (**Figure 4C**). In the thalamus, sub-regions that were densely innervated by ALM were enriched in neurons with choice selective activity (**Figure 4B, C**) (Z. V. Guo et al. 2017).

**Figure 4.**
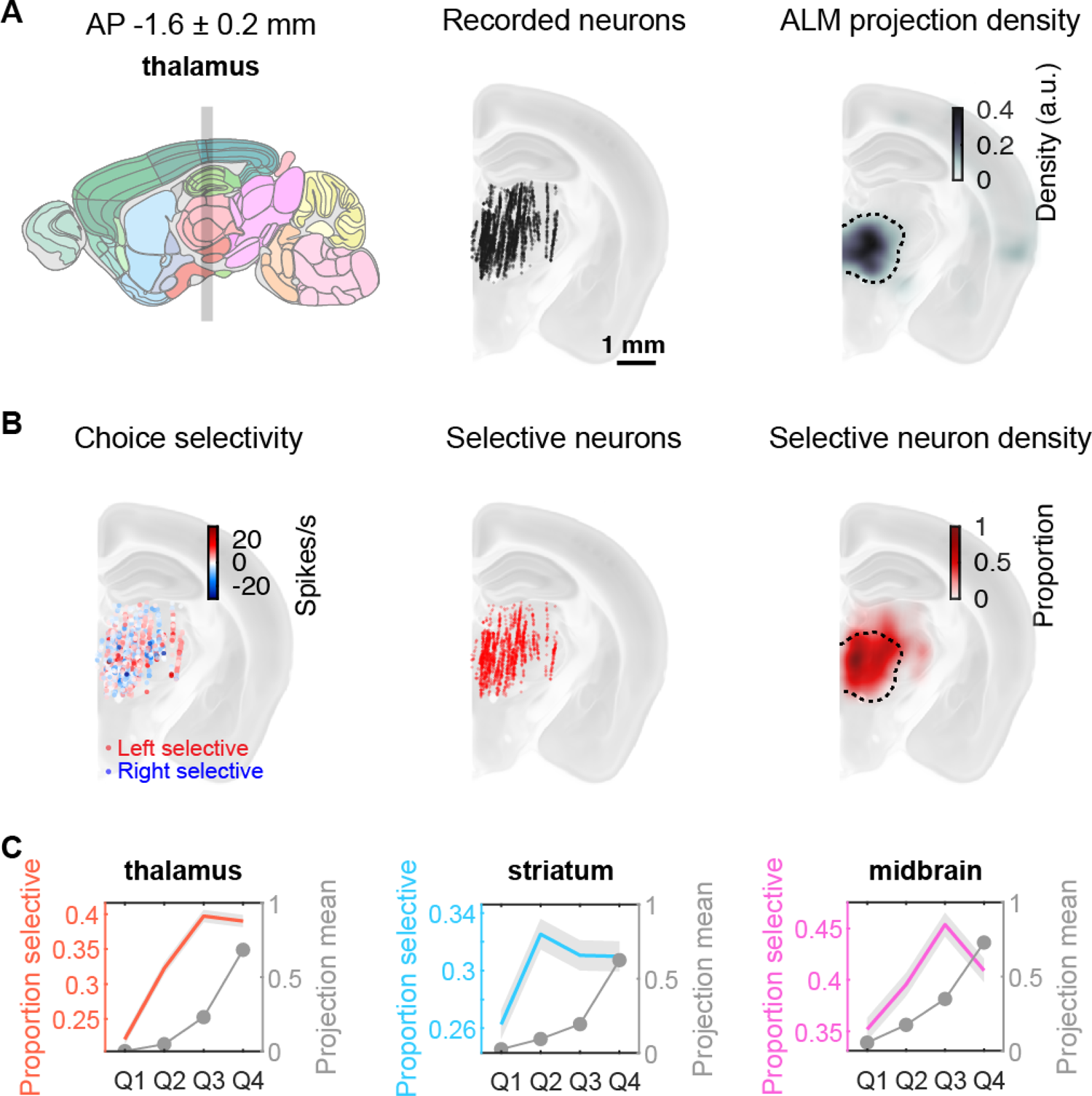
Encoding of choice in ALM projection zones. A. Left, sagittal view of the mouse brain. Gray shading, coronal section (AP −1.8 to −1.4) through the thalamus, used in the remaining panels in A and B. Middle, all recorded neurons in the thalamus. Right, ALM projection density in the thalamus. Dashed line, contour line 90% of peak density. B. Left, choice selectivity (spikes/s) in the thalamus. Red circles, lick left selective; blue circles, lick right selective. Middle, all significant choice selective neurons (rank sum test, P < 0.05). Right, proportion of choice selective neurons normalized by the density of recorded neurons (smoothed with a 3D Gaussian, sigma = 150 µm). Dashed line, same as in A. C. Proportion of choice selective neurons in thalamus, striatum and midbrain as a function of ALM projection density. In each subcortical area, neurons were divided into four quartiles (x-axis: Q1-Q4) based on overlap with ALM projection density. Gray shading, 95% confidence interval based on binomial distribution. See sFigure 4 for details.

Regions of the striatum that are innervated by ALM were enriched in neurons with choice-selective activity (**Figure S4A**). A similar situation holds for the midbrain. The lateral, deeper, motor-related part of the superior colliculus (SCm) and MRN that receives direct input from ALM exhibited strong choice-selective signals (**Figure S4B**). In contrast, the superficial, sensory-related part of SC (SCs), which does not receive input from ALM, contained little choice information (**Figure S4B**).

These measurements show that choice-selective activity is concentrated in a sparse subset of brain areas that are downstream of ALM, and form multi-regional loops with ALM via the thalamus.

### ALM drives mesoscale choice activity

We directly measured the influence of ALM on these downstream subcortical regions by comparing activity and selectivity in control trials, and in trials in which ALM was transiently (0.5 s) inactivated by photoinhibition late in the delay epoch (**Figure 5A**). Photoinhibition reduced ALM activity to 7.7 ± 0.3 % of control activity (Methods) (Nuo Li et al. 2019; Z. V. Guo et al. 2014).

**Figure 5.**
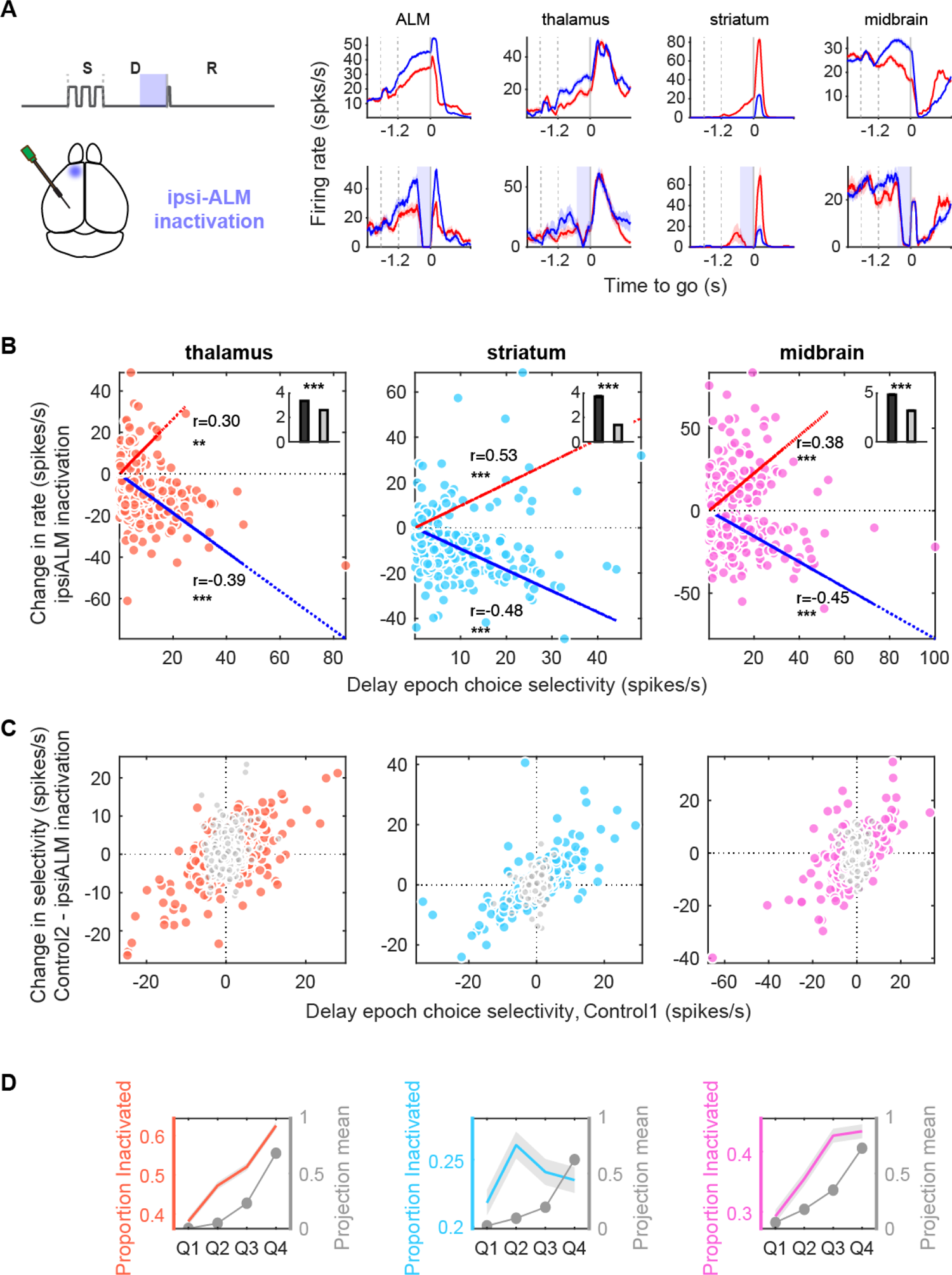
ALM drives mesoscale choice activity. A. Left, schematic of photoinhibition (blue shading) of ALM (S: sample epoch; D: delay epoch; R: response epoch). Photoinhibition was during the last 0.5 s of the delay epoch. Right, peri-stimulus time histograms for example neurons from ALM, thalamus (a mediodorsal nucleus), striatum (caudoputamen), and midbrain (superior colliculus) during control trials (upper row) and ipsi-ALM photoinhibition trials (bottom row). Blue, correct contra-trials; red, correct ipsi-trials. Vertical dashed lines separate behavioral epochs. B. Relationship between change in spike rates of individual neurons, 20-120 ms after ipsi-ALM photoinhibition onset, and delay-epoch choice selectivity in control trials for thalamus (left), striatum (middle), and midbrain (right). Dotted lines, linear regression for increasing group (red) and decreasing group (blue) (r, pearson correlation coefficient, *** P < 0.001; ** P < 0.01; bootstrapping rejects the null hypothesis that coefficient is zero). Inset, choice selectivity of significantly modulated (black bar, rank sum test, P < 0.05) and non-modulated (gray bar) neurons in corresponding regions. C. Selectivity change of individual neurons, 20-120 ms after ipsi-ALM photoinhibition, as a function of choice selectivity during control trials for thalamus (left), striatum (middle), and midbrain (right). In order to decorrelate selectivity change under perturbation against selectivity during control, we divided control trials into two groups: one group (Control 1) for evaluating choice selectivity under control conditions and another group (Control 2) for calculating choice selectivity change when compared to inactivation trials. Color dots, neurons with significant change in spike rate during inactivation (rank sum test, P < 0.05); gray dots, non-modulated neurons. D. Proportion of significantly inactivated neurons in each subcortical region as a function of ALM projection density (same as Figure 4).

The effect of silencing ALM on downstream regions was quantified as the change in spike rate (20 to 120 ms after photoinhibition onset, late delay inactivation, Methods), averaged over trials. Inactivation of ALM caused a net reduction in spike rate in a large proportion of neurons in the striatum, thalamus, midbrain and medulla (control vs. ipsilateral ALM photoinhibition: 21.1% in striatum, 53.7% in thalamus, 32.5% in midbrain, 11.3% in medulla, rank sum test, P < 0.05; median change in spike rate: striatum 5.5 Hz, thalamus 6.9 Hz, midbrain 7.5 Hz, and medulla 6.3 Hz) (Z. V. Guo et al. 2017), but a subset of neurons were excited (control vs. ipsilateral ALM photoinhibition: 4.5% in striatum, 4.0% in thalamus, 13.1% in midbrain, 11.3% in medulla, rank sum test, P < 0.05; median change in spike rate: striatum 5.5 Hz, thalamus 6.4 Hz, midbrain 9.9 Hz, and medulla 7.7 Hz) (**Figure 5B**). In all brain areas, the magnitude of a neuron’s spike rate change during ALM photoinhibition was correlated with its delay choice selectivity under control conditions (**Figure 5B**). Consistently, neurons whose spike rate was not changed by ALM photoinactivation had significantly less choice selectivity (**Figure 5B** insets, rank sum test, P < 0.001).

Furthermore, ALM silencing caused stronger changes in choice selectivity in more choice selective neurons in all downstream subcortical regions (**Figure 5C**). Neurons with highly reduced activity and selectivity during ALM photoinhibition therefore carried a disproportionate amount of choice information. Suppression of activity by ALM photoinhibition was most pronounced in ALM projection zones (**Figure 5D**).

These results indicate that ALM constitutes a major source of driving excitation to downstream circuits, and inhibits a subset of neurons through multi-synaptic pathways. For example, ALM innervates the thalamic reticular nucleus which inhibits thalamic relay cells in other nuclei (Winnubst et al. 2019; Z. V. Guo et al. 2017). GABAergic SC neurons locally suppress neighboring neurons, and inactivation of these neurons increases the activity of nearby excitatory neurons. Similarly, the two midbrain hemispheres also inhibit each other via long-range projection, and inactivating one hemisphere of midbrain therefore disinhibits the other side (Sprague 1966; C. A. Duan, Erlich, and Brody 2015; Essig, Hunt, and Felsen 2021) (**Figure S5A, B**).

Choice selectivity in subcortical regions therefore requires ALM input. The thalamus also receives input from subcortical areas, such as SC, SNr, and deep cerebellar nucleus (DCN), and these subcortical structures in turn receive direct or indirect input from ALM (Inagaki, Chen, Daie, et al. 2022). The precise roles of these long-range loops, alone or jointly, in generating and maintaining choice selectivity remain to be elucidated.

### Emergence of choice signal in ALM, thalamus, and midbrain

Our data show that choice selectivity is maintained by a multi-regional network that includes ALM and its connected brain areas (**Figure 4, 5**). We next examined the time course of the emergence of choice information across this multi-regional network.

In a subset of neurons, choice selectivity was detected early in the sample epoch (**Figure 6A**, example neuron 1), whereas intermingled other neurons were not modulated by choice (**Figure 6A**, example neuron 2). We quantified choice selectivity for single neurons by calculating the discriminability index d’ (Green and Swets 1966), across both correct and error trials. d’ was calculated using spike rates (200 ms bin) at different time points during the trial (Methods). The number of choice-selective neurons (d’ > 0.6) increased shortly after the start of the sample epoch (**Figure 6B, Figure S6A**), first in ALM and midbrain, followed by medulla, thalamus, and striatum (**Figure 6C**).

**Figure 6.**
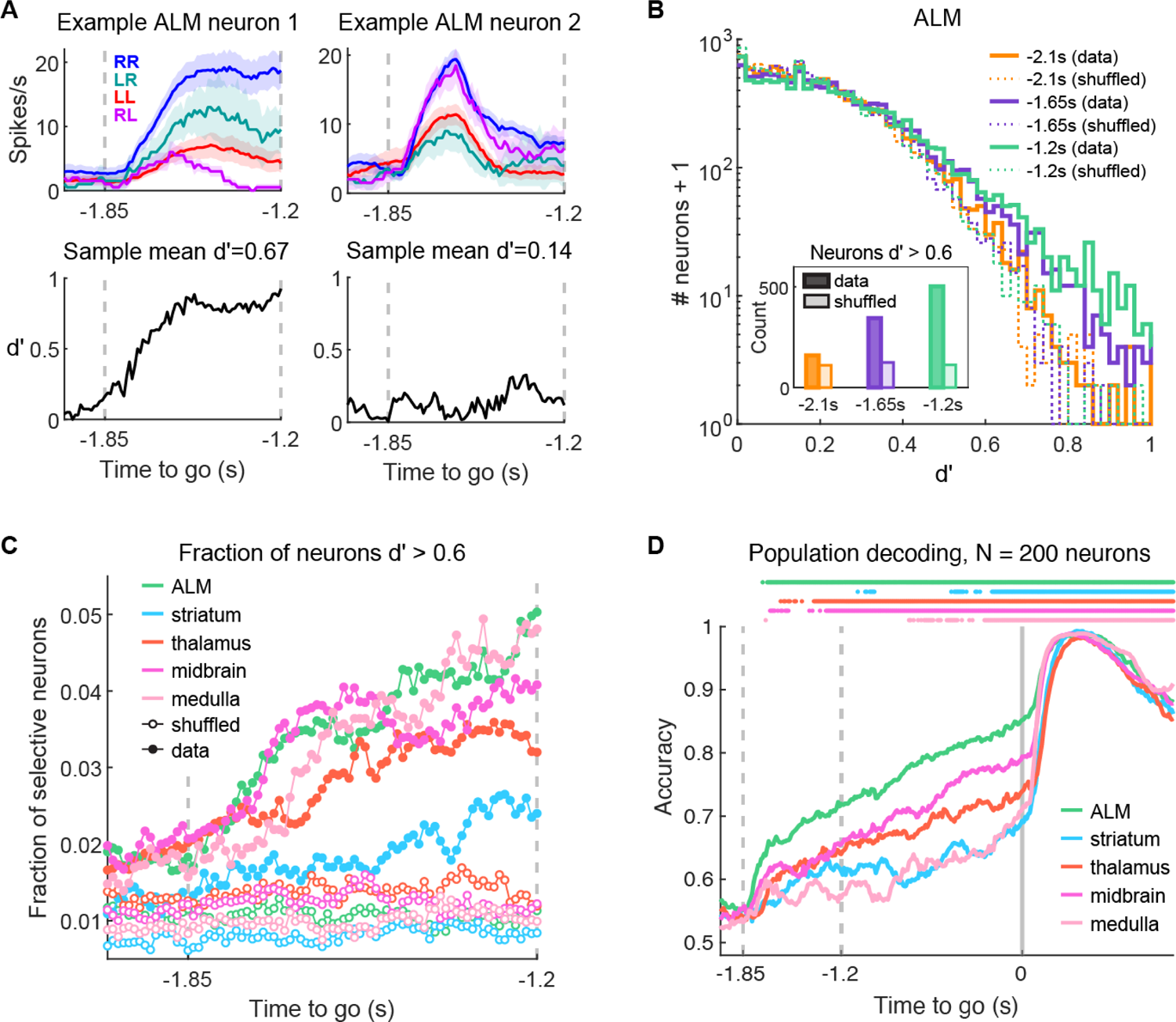
Emergence of choice signals across the brain. A. Peristimulus time histogram (PSTH) for two example neurons in ALM, focused on the sample epoch. Left, neuron with choice selectivity; right, neuron with selectivity for trial type but not choice. Top, mean PSTH in each trial group, labeled according to stimulus-choice (LL, RR, LR, RL; see Figure 2A). Shades are S.E.M. Bottom, same neuron with choice-selectivity computed as d’. B. Distribution of single-neuron choice selectivity (d’, horizontal axis) in a 10,000-neuron pseudo-population from hierarchical bootstrapping in ALM. Colors indicate different time points in pre-sample epoch (−2.1 s), mid sample epoch (−1.65 s), and the end of sample epoch (−1.2 s). Solid lines are distributions of data. Dashed lines are generated with the same method as solid lines, but from data with shuffled trial IDs. Inset, number of neurons with high choice selectivity (d’ > 0.6) at different time points for both data and data with shuffled trial IDs. C. The proportion of neurons with high choice selectivity (d’ > 0.6) in each brain area as a function of time. Each pseudo-population contains 10,000 neurons from hierarchical bootstrapping. Solid circles are calculated from data. Open circles are calculated in the same way but with shuffled trial IDs. Causal time bin, 200 ms, step size, 10 ms. D. Population choice decoding accuracy in each area as a function of time. Each time a pseudo-population of 200 neurons were subsampled from hierarchical bootstrapping in one area. The accuracy was the mean over 100 repetitions of hierarchical bootstrapping, and the standard error of mean of all regions is within 0.10. Bars at the top indicate times when the decoding accuracy of a brain area is larger than 0.5 in at least 95% of the repetitions. Time bin, 200 ms, step size, 10 ms.

Choice information might be encoded in weakly selective populations of neurons, before individual selective neurons can be detected. We thus used population decoding to compare the emergence of choice information across brain areas. To classify choice irrespective of other task variables, we assigned different weights to trials to address the issue of imbalanced number of correct and error trials, so the choice variable was uncorrelated with other task variables. Given that decoding accuracy increased with the number of recorded neurons, and that the number of neurons recorded differed across recording sessions and mice (**Figure S6B, D**), we performed decoding on pseudo-populations constructed using hierarchical bootstrapping (**Figure S6C**) (Z. V. Guo et al. 2014; Saravanan, Berman, and Sober 2020). We subsampled 200 neurons in each area from 1) individual mouse, 2) sessions within each mouse, and 3) neurons and trials within each session (Methods). We then used a linear decoder with regularization to classify choice from the population activity at each time point (bin step 10ms, bin size 200ms; logistic regression with nested 5-fold cross validation; the whole process was repeated 100 times; Methods). Population decoding showed that choice information developed in the sample epoch and gradually increased throughout the delay epoch (**Figure 6D**). The ramp-up of choice information occurred across ALM and its projection zones, but not in other areas, such as the hippocampus and olfactory cortex, and only weakly in other cortical areas (**Figure S6H**). To compare when choice information emerged across brain areas, we quantified the time points when each region’s neural population has a decoding accuracy above 0.6 (**Figure 6D**; conclusions did not change with a reasonable range of thresholds). ALM population showed the earliest and strongest choice information, followed by ALM projection zones in midbrain, thalamus, medulla, and striatum (**Figure 6D**).

We note that the early emergence of choice signals in ALM does not imply that decisions are computed in ALM and then transmitted to connected areas. Choice decoding improves with the number of sampled neurons: for example, a population of 300 thalamus neurons show similar early choice information as 100 ALM neurons (**Figure S6F, G**), and the size of the behaviorally relevant population in any brain area is unknown. In addition, previous experiments have shown that choice-related activity in ALM requires input from the thalamus, basal ganglia, and cerebellum (Z. V. Guo et al. 2017; Gao et al. 2018; Y. Wang et al. 2021; Tang et al. 2021). Together, these experiments suggest inherently multi-regional mechanisms underlying decision-making.

### Correlated encoding across brain areas

Our brain-wide recordings revealed parallel representations of stimulus, choice, and action across anatomically coupled brain areas. How is task-related activity coordinated across brain areas? We examined the dynamics of task-related activity across simultaneously recorded brain areas.

We analyzed activity of each region in an activity space, where individual dimensions correspond to the activity of individual neurons. Over time population activity traces out a trajectory in this activity space. Using both correct and error trials, we defined a direction in activity space that maximally discriminated upcoming choice (lick left vs. lick right trials) during the late delay epoch (‘choice’ coding direction, **CD_choice_**) (N. Li et al. 2016; G. Chen et al. 2021; Inagaki, Chen, Daie, et al. 2022) (**Figure 7A**). Population activity projected onto the **CD_choice_** diverges in a choice-specific manner during the delay epoch (projection onto **CD_choice_**, pCD_choice_), providing a short-term memory of the upcoming choice (**Figure 7A**).

**Figure 7.**
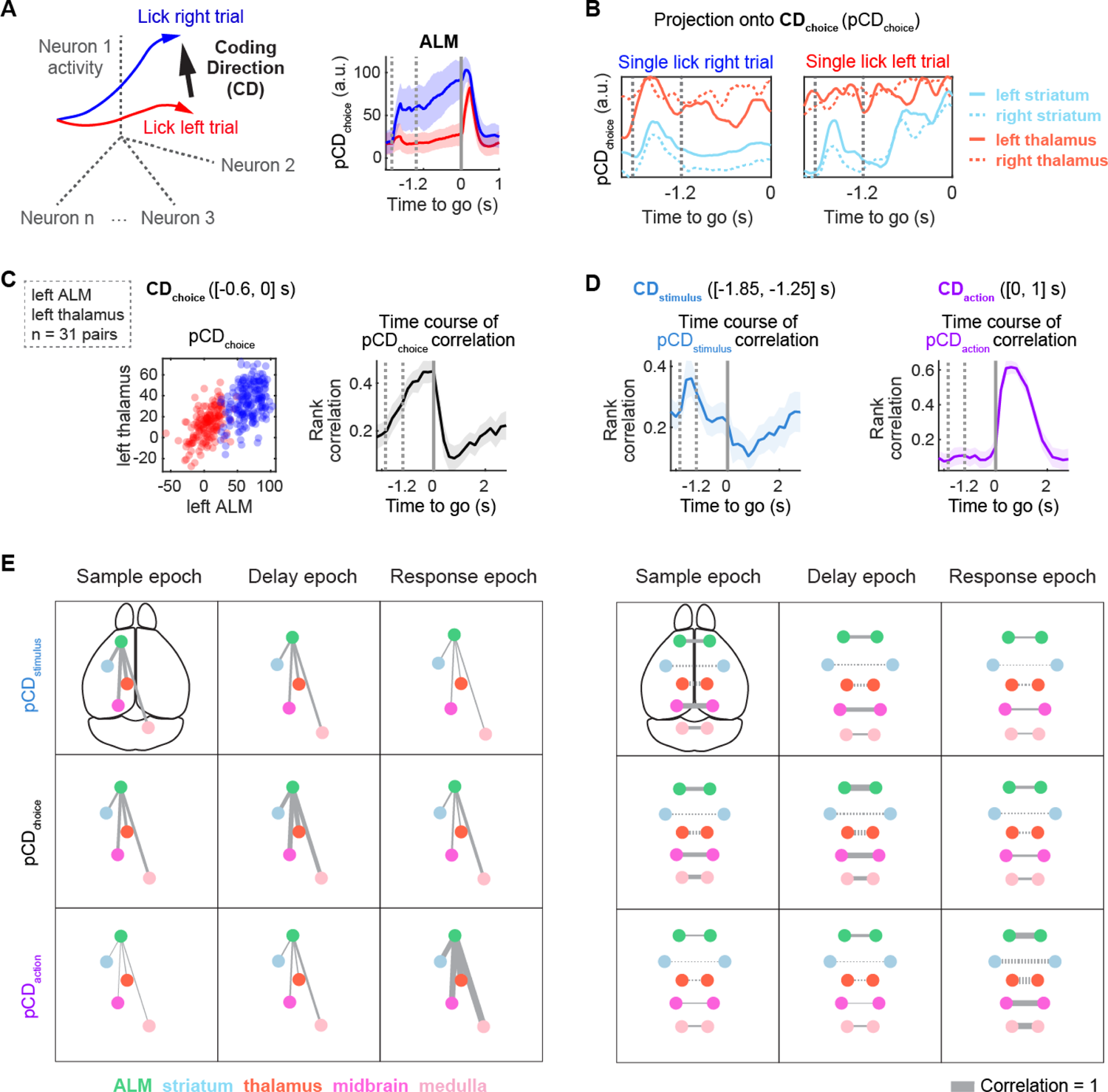
Correlated encoding across brain areas. A. Left, definition of coding direction (**CD**), a vector in activity space that best separates activity trajectories for different task conditions. For example, coding direction related to choice (**CD_choice_**) is the direction in activity space that best separates left (red) and right (blue) lick trials during the late delay epoch (0.6 s). Right, trial-averaged projection of ALM population activity onto **CD_choice_** (pCD_choice_) for one behavior session (shade, standard deviation). B. Simultaneously recorded activity in four brain areas on a lick right trial and a lick left trial, projected onto their respective choice CD (i.e. **CD_choice_**). Time bin, 200 ms. C. Left, correlations of pCD_choice_ in a pair of brain areas (left ALM, left thalamus) during late delay epoch ([-0.6, 0] s) (one example session; correct trials). Right, rank correlation calculated at different time points (200 ms time bin) and quantified across both trial types for pCD_choice_ of left ALM - left Thalamus pair (n = 31). Mean ± S.E.M. across sessions. D. Same data as in C, but now projected onto **CD_stimulus_** and **CD_action_** (directions in activity space separating stimulus- and action-related activity, respectively). Rank correlations were calculated in 200 ms time bins and are quantified across both trial types. Mean ± S.E.M. across sessions. E. Functional connectivity, measured using rank correlation, across five brain areas and different **CD**s. Colored circles represent brain areas. The widths of lines connecting brain areas are proportional to rank correlations. Left, ALM with ipsi- Striatum, ipsi-Thalamus, ipsi-Midbrain, and contra-Medulla. Right, five brain areas across hemispheres. Solid lines, direct connections; dashed lines, weak or absent direct connections (e.g. two hemispheres Striatum, and two hemispheres Thalamus).

We investigated how the representation of choice information is coordinated across brain areas, with **CD_choice_** computed for each brain area. In single trials, pCD_choice_ trajectories fluctuated in a correlated manner across brain areas (**Figure 7B**). Especially at the end of the delay epoch, pCD_choice_ were highly correlated across brain areas (**Figure 7C**). For example, when the left hemisphere ALM signaled lick left or lick right (error trials or correct trials), the left hemisphere thalamus usually signaled the same direction. Inter-regional correlations in pCD_choice_ developed gradually during the delay epoch, reaching a maximum in the late delay epoch (**Figure 7C, Figure S7A**). This correlated choice activity reflects functional coupling between brain areas during the delay epoch.

Notably, the inter-regional correlations collapsed rapidly after the ‘Go’ cue (**Figure 7C**), implying that functional coupling is not constant across behavioral epochs. To explore functional coupling across distinct behavioral epochs, we defined two additional coding directions. We computed i) a direction in activity space that maximally discriminated trial type as defined by the sensory stimulus during the sample epoch (high vs. low tone, i.e. ‘stimulus’ coding direction, **CD_stimulus_**) and ii) a direction in activity space that discriminated lick direction during the response epoch (lick right vs. lick left, i.e. ‘action’ coding direction, **CD_action_**) (Methods; **Figure 7D**). Activities between brain areas were correlated along both directions (stimulus, pCD_stimulus_; action, pCD_action_), but functional couplings along **CD_stimulus_** and **CD_action_** occurred during different epochs. Functional coupling along **CD_stimulus_** peaked soon after stimulus onset, but decayed during the delay epoch (Finkelstein et al. 2021). In contrast, functional coupling between **CD_action_** developed rapidly after the ‘Go’ cue and was strong during movement execution (**Figure 7D, E, Figure S7A, B**). These results suggest that brain areas are functionally coupled in distinct subspaces of population activity during different phases of behavior.

We quantified this time-dependent functional coupling between major pairs of brain areas as the strength of activity correlation along **CD_stimulus_, CD_choice_**, and **CD_action_** in different behavioral epochs (**Figure 7E, Figure S7B**). During the sample epoch, all the sampled brain area pairs preferentially exhibited functional coupling along **CD_stimulus_**, with the two sides of the midbrain showing the strongest coupling. During the delay epoch, ALM was strongly coupled to most connected brain areas, primarily along **CD_choice_**, but exhibited weaker coupling to ipsilateral medulla (**Figure S7**). During the response epoch, coupling was strongest between ALM and ipsilateral midbrain and contralateral medulla, specifically along **CD_action_**. Within each task epoch, functional coupling was observed across all the brain area pairs, even between brain areas without direct anatomical connections (**Figure 7E, Figure S7**, e.g. left and right striatum). Functional coupling rapidly switched between distinct subspaces of population activity across different phases of behavior and this switch occurred simultaneously for all the brain area pairs. These highly coordinated changes in functional coupling across the multi-regional network may reflect routing of distinct task-related information across brain areas during different phases of behavior (Druckmann and Chklovskii 2012; Kaufman et al. 2014; Semedo et al. 2019).

## Discussion

We measured neural activity across a connected pre-motor network during a memory-guided movement task in mice (Figure 1). Behavior-related neural activity was widely distributed (Figure 2, Figure 3), but choice-related activity was concentrated in connected multi-regional loops (Figure 4). Choice-related activity was first detected in ALM (Figure 6), and choice signals in subcortical regions require input from ALM (Figure 5). Choice signals are highly correlated across brain areas and collapse in a coordinated manner at the time of movement onset (Figure 7).

### Anatomy-guided measurements of multi-regional neural activity

The neural computations underlying behavior, such as decision-making, action selection, and goal-directed movement, rely on neural circuits spanning multiple brain areas. For example, classic work in non-human primates has shown that multiple connected cortical (Shadlen and Newsome 2001; Katz et al. 2016) and subcortical (E. J. Jun et al. 2021; Ding and Gold 2010) brain areas show similar ramping activity during decision-making based on visual motion signals. Even in such well-studied systems and behaviors, ascribing a specific function to a brain area has remained elusive, suggesting that neural computations may be an emergent function of multi-regional neural circuits (Latimer and Huk 2021). Deciphering the underlying neural mechanisms requires recordings from networks of connected neurons across multiple brain areas.

Such recordings are challenging because multi-regional neural circuits are complex and, apart from a small number of model systems, such as *C. elegans*, *Drosophila*, and mice, have been only sparsely mapped (Markov et al. 2014). Among mammals, our understanding of mesoscale connectivity is most advanced for mice, based on multiple, large-scale mapping projects (Zingg et al. 2014; Oh et al. 2014). The projection patterns across brain areas rarely obey the compartment boundaries shown in classic anatomical atlases (https://mouse.brain-map.org/static/atlas) (Paxinos and Franklin 2004). To align anatomical and functional measurements, and to combine data across individual mice, we registered measurements of neural activity and connectivity to a standardized brain coordinate system (Q. Wang et al. 2020; L. D. Liu et al. 2020) (common coordinate framework, CCF).

We targeted multiple Neuropixels probes to brain areas that form loops with anterior lateral motor cortex (ALM), guided by databases of multi-regional connectivity (Oh et al. 2014; Winnubst et al. 2019). ALM anchored our circuit analysis because a large body of work in multiple laboratories has shown ALM to be a critical circuit node in behaviors requiring the planning and execution of goal-directed licking (Z. V. Guo et al. 2014; Economo et al. 2018; Inagaki, Chen, Ridder, et al. 2022; N. Li et al. 2015; Inagaki et al. 2018; Chunyu A. Duan, Pan, et al. 2021). ALM was originally defined based on behavioral effects of inactivation experiments (N. Li et al. 2016; Z. V. Guo et al. 2014; Inagaki et al. 2018; Komiyama et al. 2010) and overlaps with anterior parts of primary and secondary motor cortex (https://mouse.brain-map.org/static/atlas). ALM is part of multiple closed loops with subcortical brain areas that are known to play roles in action selection and execution (Inagaki, Chen, Daie, et al. 2022; Z. V. Guo et al. 2017; N. Li et al. 2015; Chunyu A. Duan, Pan, et al. 2021; Gao et al. 2018). This multi-regional connectivity is complex, sparse, and specific (Zingg et al. 2014; Harris et al. 2019; Winnubst et al. 2019), with projection zones that can be as small as 200 micrometers (Lee, Wang, and Sabatini 2020; K. Guo et al. 2018). For example, ALM projects to parts of the ventral medial (VM) and mediodorsal (MD) regions of the thalamus, but not to many other parts of the thalamus (Z. V. Guo et al. 2017; K. Guo et al. 2018). ALM projects to lateral SCm, but not to medial and dorsal sensory SC (Figure 1). In this study, a combination of histology and electrophysiological landmarks was used to localize individual recorded units to better than 100 micrometers in the CCF (L. D. Liu et al. 2020), and thus relate neural signals and multi-regional connectivity.

We recorded with up to five Neuropixels probes, often from six or more connected brain areas (Figure 1 H). Here we focus on activity in ALM and its projection zones. These projection zones often partially occupy multiple nearby brain areas. In one widely used 3-D anatomical atlas, brain structures are part of hierarchies up to eight levels deep (Q. Wang et al. 2020). In our analysis we specify ALM using the full depth of the hierarchy (Cerebrum / cerebral cortex / cortical plate / isocortex / somatomotor area / secondary motor area / ALM). In analyzing ALM projection zones we average across multiple related brain areas, and thus specify only shallow levels of the hierarchy (brainstem / interbrain / thalamus; brainstem / midbrain; brainstem / hindbrain / medulla). Future analyses will consider sub-divisions within ALM projection zones that correspond to distinct thalamic, midbrain and hindbrain structures.

Encoding of sensory stimuli, choice, action and outcome were widely distributed across the brain (Huber et al. 2012; Hill et al. 2011; Yu et al. 2016; Steinmetz et al. 2019; Allen et al. 2019; Peters et al. 2021; Petreanu et al. 2012). In the cortex, choice-related activity is concentrated in ALM (T. W. Chen et al. 2017). Similarly, in subcortical brain areas choice-related activity was enriched in brain areas that receive input from ALM, and weaker in surrounding areas (Figure 4). Moreover, choice-selective activity in these subcortical brain areas depends on input from ALM (Figure 5). These measurements show that choice-related activity is generated and maintained in a sparse, multi-regional neural circuit including ALM, and specific parts of the thalamus (Z. V. Guo et al. 2017), basal ganglia (Y. Wang et al. 2021), midbrain (Chunyu A. Duan, Pan, et al. 2021) and hindbrain (Gao et al. 2018).

Choice-related activity is produced in part by attractor dynamics (Finkelstein et al. 2021; Inagaki, Chen, Daie, et al. 2022). The attractors underlying our binary decision task are likely learned based on experience-dependent synaptic plasticity (Hopfield 1982). Does wide-spread choice-related activity imply widely distributed changes in synaptic weights? Recent theoretical work has shown that in strongly coupled neural circuits, synaptic changes in a small subset of the network can cause wide-spread changes in task-related neural dynamics (Kim et al. 2022). Experience-dependent changes in connectivity are a hallmark of cortical circuits (Holtmaat et al. 2006; Trachtenberg et al. 2002), including the motor cortex (Peters, Liu, and Komiyama 2017), as well as corticostriatal synapses. Experience-dependent synaptic plasticity in a limited number of circuit nodes could therefore underlie changes in neural activity across the entire premotor circuit. Our experiments do not address the question if choice-related activity in all circuit nodes is equally important in driving future actions.

### Origins of choice-related activity

Analysis of latencies can reveal how computations evolve across multi-regional neural circuits. For example, numerous studies have measured neuronal latency with respect to presentation of a sensory stimulus across brain areas (Siegel, Buschman, and Miller 2015; Siegle et al. 2021; Steinmetz et al. 2019; Mormann et al. 2008; Schmolesky et al. 1998; Yu et al. 2016). One previous study has mapped the neural signals antecedent of movement initiation across the brain, revealing a wave of activity arising in the brainstem, propagating through the thalamus to the cortex (Inagaki, Chen, Ridder, et al. 2022; Schwab et al. 2020). Measurement of the short latency differences across connected brain areas, a few milliseconds, was enabled by the brief time between the ‘Go’ cue and movement initiation, as well as the phasic nature of the neural signal.

Few studies have compared the latency of choice information across brain areas under standardized task conditions, with inconsistent conclusions. Previous studies report that choice activity arises nearly simultaneously in frontal and parietal cortices (Hernandez et al. 2010; Siegel, Buschman, and Miller 2015). One of these studies finds choice activity in the frontal cortex before sensory cortex (Siegel, Buschman, and Miller 2015), whereas other studies find similar latencies (Hernandez et al. 2010). One study could not differentiate the timing of choice activity between frontal cortex and midbrain (Steinmetz et al. 2019), whereas another study finds earlier choice activity in superior colliculus than frontal cortex (Chunyu A. Duan, Pagan, et al. 2021).

For choice-related activity, the measurement of latency is challenging since it develops gradually over hundreds of milliseconds (Hernandez et al. 2010; Siegel, Buschman, and Miller 2015; Hanks et al. 2015), whereas time-constants of neural communication across brain areas are on the order of 10 milliseconds (Z. V. Guo et al. 2017; Inagaki, Chen, Ridder, et al. 2022). Aided by our large data set, we relied on decoding of choice based on neural activity of individual neurons and neural populations. Choice-related information appeared first and most strongly in ALM, closely followed by the midbrain and thalamus, with a similar time-course (Figure 6).

We note that our decoding analysis is limited. The decoding strength depends on the number of neurons included in the analysis, and as the number increases, differences in latencies across ALM, midbrain and thalamus shrink (Figure S6F). Moreover, previous studies have shown that input from the thalamus (Z. V. Guo et al. 2017), SNr (Y. Wang et al. 2021; Z. V. Guo et al. 2017), and DCN (Gao et al. 2018) are critical for ALM to develop and maintain choice-related activity. The most parsimonious explanation of our current study and previous data is that choice signals are computed in a multi-regional pre-motor loop, with ALM serving as a key node, in addition to connected basal ganglia, thalamus. The precise roles of midbrain structures, including the superior colliculus and midbrain reticular nucleus, remain to be investigated.

### Correlations in activity modes across the brain

Choice-related activity is strongly correlated across brain areas. Remarkably, coupling along choice-related subspaces collapses rapidly after the ‘Go’ cue across the brain. At the same time, activity in different action subspaces emerges that is also strongly correlated across brain areas. The correlated activity within distinct activity subspaces likely reflects sharing of specific task-related information across brain areas (Economo et al. 2018; Kaufman et al. 2014; Semedo et al. 2019). For example, correlated activity along the choice coding direction may reflect maintenance of choice information during the delay epoch. The correlated activity undergoes a coordinated, brain-wide switch between distinct subspaces across all brain areas. A previous study showed that the ‘Go’ cue triggers a phasic signal in the midbrain, which ascends through the thalamus into ALM (Inagaki, Chen, Ridder, et al. 2022). In addition to their projections to the thalamus, these ascending midbrain neurons project to the SNr, the subthalamic nucleus, and the striatum, providing a possible mechanism for coordinated switching across the brain.

Our approach to measuring the relationship between population activity across pairs of brain areas focused on the single-trial correlation of patterns of neural activity that predict behavior. These patterns were defined through the projection of neural activity on decoder vectors, i.e., projections on **CD_choice_**, which we refer to as pCD_choice_ (Figure 7). More generally, neural population activity is naturally expressed as high dimensional time series. Numerous ways have been suggested to quantify the relationship between two sets of high-dimensional time series (Kang and Druckmann 2020; Semedo et al. 2020). Our focus on correlation of projections on decoders was motivated by previous work, which has shown that such decoders applied to ALM, where they predict future behavior accurately, even following large scale perturbation (N. Li et al. 2016) and for continuity with a series of studies that have analyzed population activity in a similar fashion (Inagaki, Chen, Daie, et al. 2022). Future efforts will explore the tradeoff between robustness across individual animals, the amount of data necessary for stable inference, and the expressivity of different methods, in service of measuring the influence of one brain area dynamics on another.

### Coding of movement

Cortical activity is influenced by movement, including movements that are not instructed. In the sensory cortex the movement of eyes (Herrington et al. 2009; Martinez-Conde, Macknik, and Hubel 2000), digits (Papakostopoulos, Cooper, and Crow 1975), and whiskers (Crochet and Petersen 2006; Yu et al. 2016) as well as locomotion (Niell and Stryker 2010; Keller, Bonhoeffer, and Hübener 2012) can drive neural activity and modulate sensory coding. The advent of high-speed videography has revealed that ongoing movements are represented in patterns of neural activity across the forebrain and other brain areas (Musall et al. 2019; Stringer et al. 2019; Huber et al. 2012; Salkoff et al. 2020; Hill et al. 2011).

Prima facie, correlations of movement and neural activity are expected to be wide-spread based on neural connectivity. For example, ALM pyramidal tract neuron activity is modulated in multiple phases of behavior (N. Li et al. 2015; Economo et al. 2018). These neurons project to premotor circuits in the superior colliculus and the medulla and it would be surprising if this modulation was not reflected in the activity of motor neurons and muscle activation. In fact, neurons in the intermediate nucleus of the reticular formation in the medulla, premotor to the hypoglossal nucleus (N. Li et al. 2015; McElvain et al. 2018), show preparatory activity that is dependent on ALM (data not shown). Conversely, ALM receives channels of sensory input from the midbrain (via thalamus) (Inagaki, Chen, Ridder, et al. 2022) and somatosensory cortex, providing pathways for modulation of ALM activity by reafference and proprioceptive input. Finally, broadly projecting neuromodulatory systems, including serotonergic, cholinergic and noradrenergic neurons, project to motor systems and the forebrain and might cause correlations between forebrain neural activity and movements.

We measured encoding of movement by quantifying how well neural activity could be predicted from behavioral video (Figure 3). Consistent with previous measurements, movements, including uninstructed movements, were encoded by neurons distributed across the brain. Encoding was strongest in the hindbrain, followed by the midbrain and then the forebrain. In ALM, a considerable proportion of neurons showed little or no encoding of movement.

Cognitive neuroscience sometimes views decision-making and motor planning as separate from movement control. However, decisions may well involve motor structures and be reflected in an ongoing manner in posture and movement. For example, cognitive signals descend to brainstem nuclei that control facial muscles, including pupil size (Kahneman & Beatty 1966, Joshi & Gold 2020); in perceptual decision-making tasks, decision variables are reflected in the reflex gains in skeletal muscles (Selen et al. 2012).

In our data, in a subset of mice and sessions, movements are observed during motor planning and correlate with choice-related activity. It is unknown whether choice-related activity causes idiosyncratic movements, movement contributes to preparatory activity, or preparatory activity and movement are modulated by a common input. Movements could even be part of an embodied loop of motor planning, and may have to be considered when dissecting the neural circuits underlying planning. To answer these questions, future experiments will have to selectively block descending signals to motor centers, or ascending signals from sensory areas that encode reafference or proprioception.

## Data availability

Aligned electrophysiological and behavioral data, including metadata, has been deposited on dandiarchive.org in the NWB2 format. Reconstructed ALM neurons are available at https://ml-neuronbrowser.janelia.org/ (Table 3 for DOIs). Other data that support the findings of this study are available upon request.

## Acknowledgement

We thank H.K. Inagaki, J.H. Siegle and M.N. Economo for comments on the manuscript; B. Karsh for software development; J. Arnold for the design of the multi-probe insertion system. W.L. Sun for help with Neuropixels probes; T. Nguyen for Datajoint pipeline and data dissemination; T. Ferreira for help with the MouseLight dataset; B. Forster for whole-brain clearing; T. Pluntke and M. Inagaki for animal training. This work was funded by Simons Collaboration on the Global Brain (S.D., N.L., and K.S.), Howard Hughes Medical Institute (K.S., and T.H.), including the Janelia Visiting Scientist Program (S.C., N.L., and K.S.), and Wellcome Trust (S.C.).

## Author contributions

Conceptualization: K.S., N.L., S.D, S.C. Behavioral training: S.C., N.T. Electrophysiology: S.C. Data analysis and visualization: S.C., Y.L., Z.W., K.S., S.D., N.L. Spike sorting and quality control pipeline: J.C., S.C., N.L., L.D.L., K.S. Histology: S.C., T.W. Electrode localization: S.C., L.D.L., H.H., K.S., N.L. Writing: S.C., Y.L., S.D., N.L., K.S. Supervision: K.S., N.L., S.D., T.H., S.C. Funding acquisition: K.S., N.L., S.D., S.C., T.H.

## Supplementary figure captions

**Figure S1.**
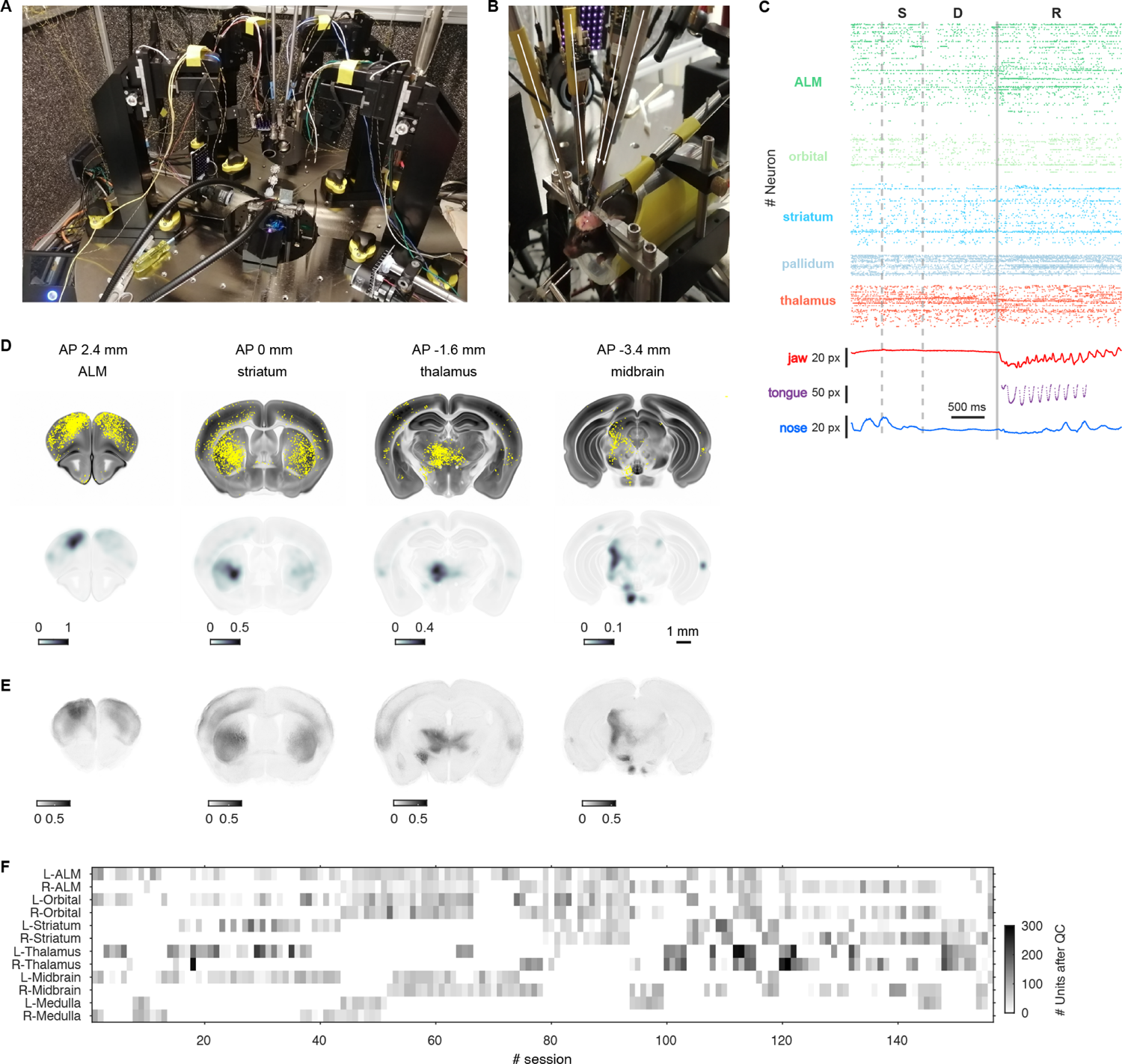
Brain-wide, anatomy-guided recordings of neural activity. A. The recording apparatus with four towers holding neuropixels probes. B. Zoomed-in view showing head-restrained mouse and a recording with four neuropixels probes. C. Example experiment with three Neuropixels probes, corresponding to insertion Group 1, 2, 3, respectively. Population raster plot showing one behavioral trial. Electrophysiological measurements were combined with movement tracking (bottom). D. Top row, four coronal sections (thickness, 20 µm) centered on ALM, striatum, thalamus and midbrain, approximately corresponding to the target of four insertion groups (see Figure 1), showing axonal segments from ALM neurons (yellow). Bottom, row, cortical projection density. Voxel intensity is the axonal protection length-density smoothed with a 3D Gaussian (sigma = 150 µm). Projection density has been normalized to the range of 0-1 across the brain. E. Anterograde tracing experiment. ALM was infected with AAV expressing GFP under the synapsin promoter. Gray levels indicate fluorescence measured in 100 µm thick sections. F. Summary of multi-regional recording data set. Horizontal axis, recording sessions, grouped by hierarchical clustering to demonstrate similar combinations of areas. Vertical axis, major brain compartments recorded in each session. Color intensity denotes the number of recorded units after quality control (QC).

**Figure S2.**
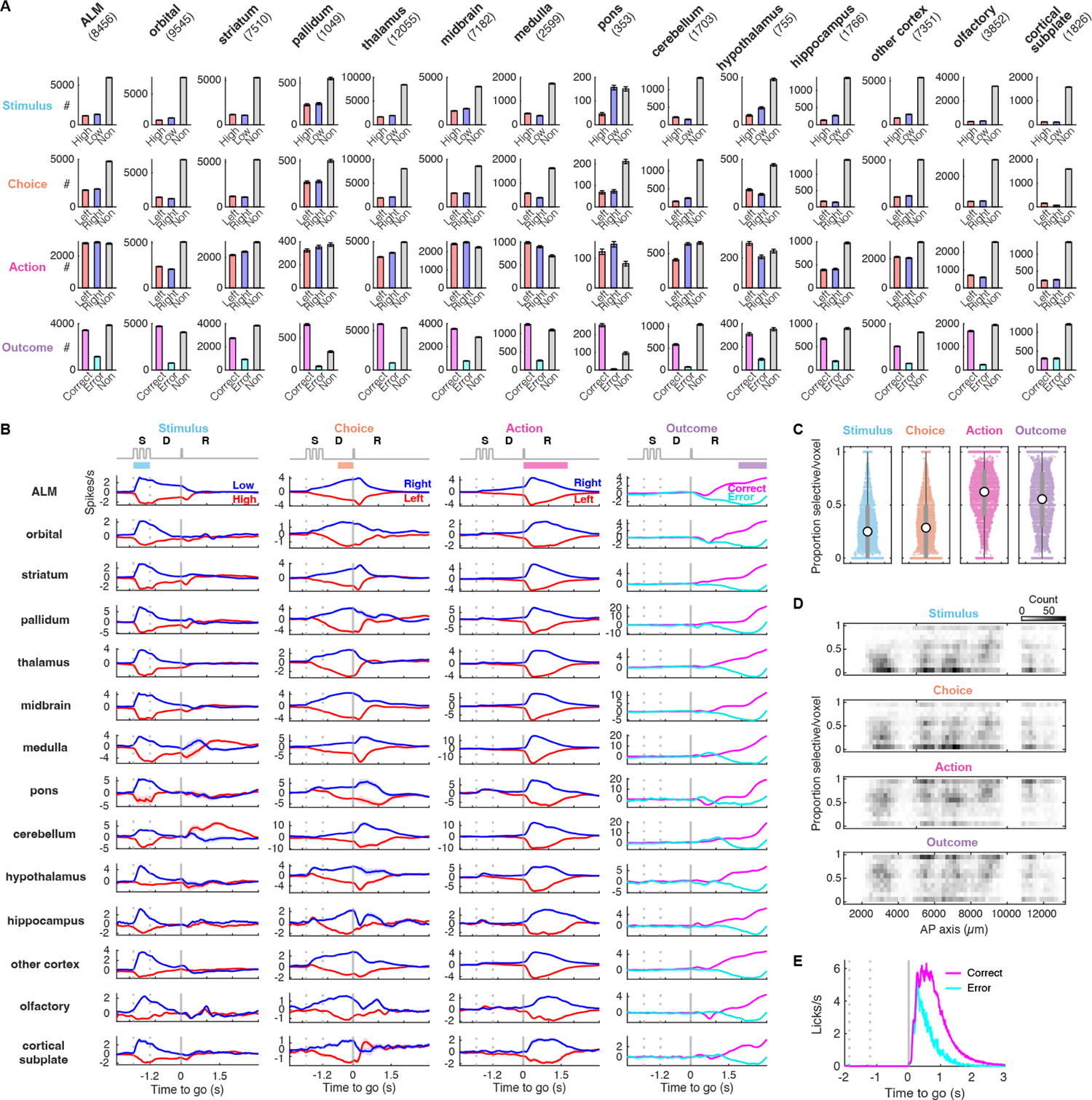
Selectivity across brain areas. A. Number of stimulus, choice, action and outcome selective (color codes same as in Figure 2) neurons in each brain area. Gray bars represent non-selective neurons. Error bars are S.E.M., determined by bootstrapping. B. Average population selectivity in spike rate (Mean ± S.E.M. across selective neurons) for stimulus, choice, action and outcome across major brain areas. Averaging window, 200 ms. C. The proportion of selective neurons across recorded voxels (volume, (200 µm)^3^, CCF) for stimulus, choice, action and outcome. D. Heatmaps of proportion of selective neurons in recorded voxels along the anterior-posterior (AP) axis for stimulus, choice, action and outcome. Note the high proportion of selective neurons distributed along a wide range of AP for action and outcome, but not for stimulus and choice, implying more prevalent and evenly distributed representations of action and outcome. E. Lick rate during correct and error trials (mean across sessions ± S.E.M.)

**Figure S3.**
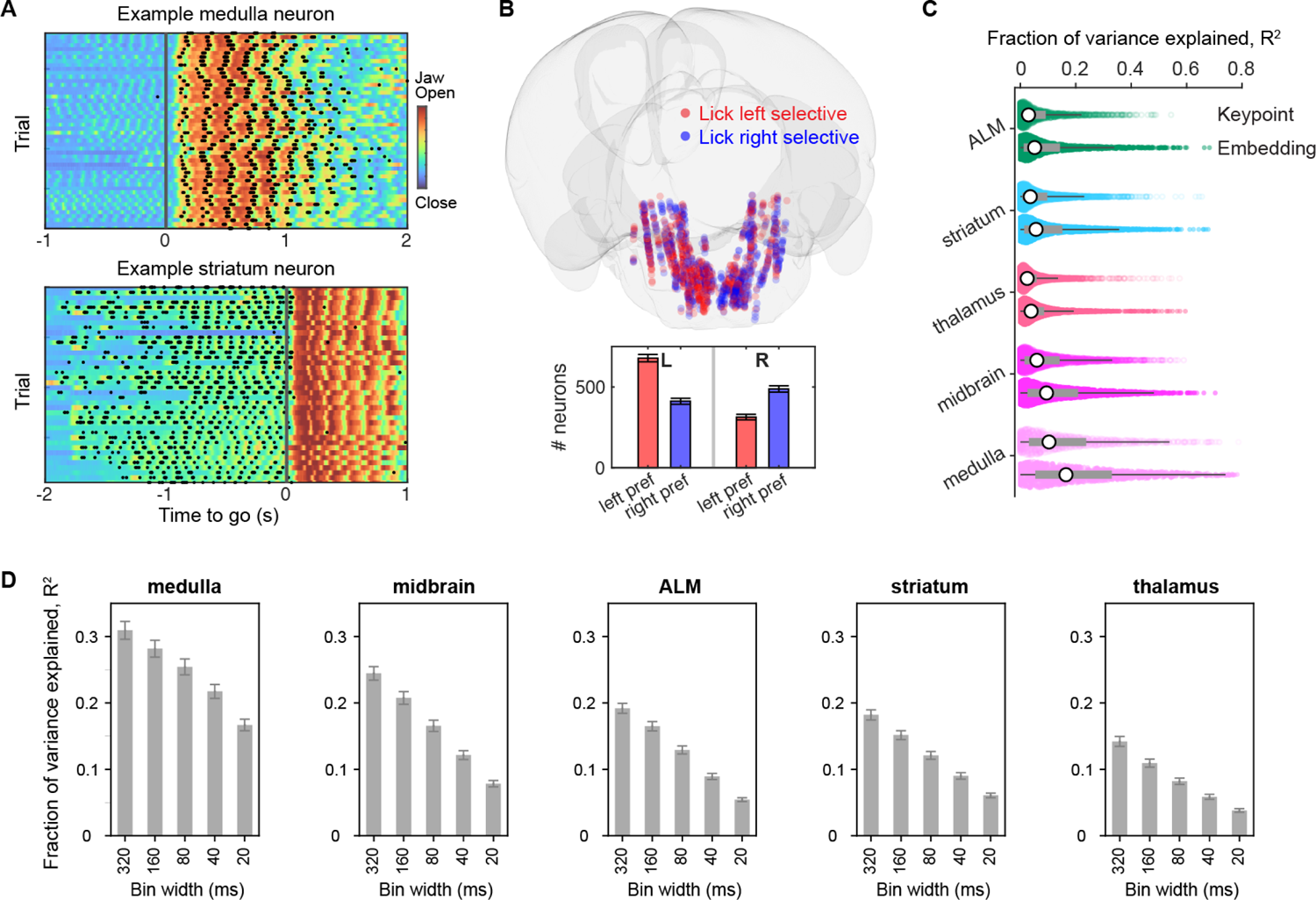
Coding of movement. A. Example neuron (spikes, black ticks) superposed on jaw position (color map) in a behavioral session with anticipatory movements during the delay epoch (time < 0). Top, medulla neuron showing oscillatory activity synchronized with jaw movement during the response epoch. Bottom, striatal neuron showing oscillatory activity during the delay epoch. B. Lateralized (ipsiversive) selectivity of medulla activity during the response epoch. Bottom, quantification for left hemisphere (L) and right hemisphere (R) respectively. Error bars are S.E.M., based on 1000 times bootstrapping. C. Fraction of variance explained (R^2^) for individual neurons across major brain areas for keypoint-based (top) and embedding-based (bottom) predictions. Encoding for movements was highest in the medulla, followed by the midbrain, striatum / ALM, and the thalamus. D. The rank ordering of movement encoding across brain areas is robust across different bin widths for averaging spike rates used for predictions.

**Figure S4.**
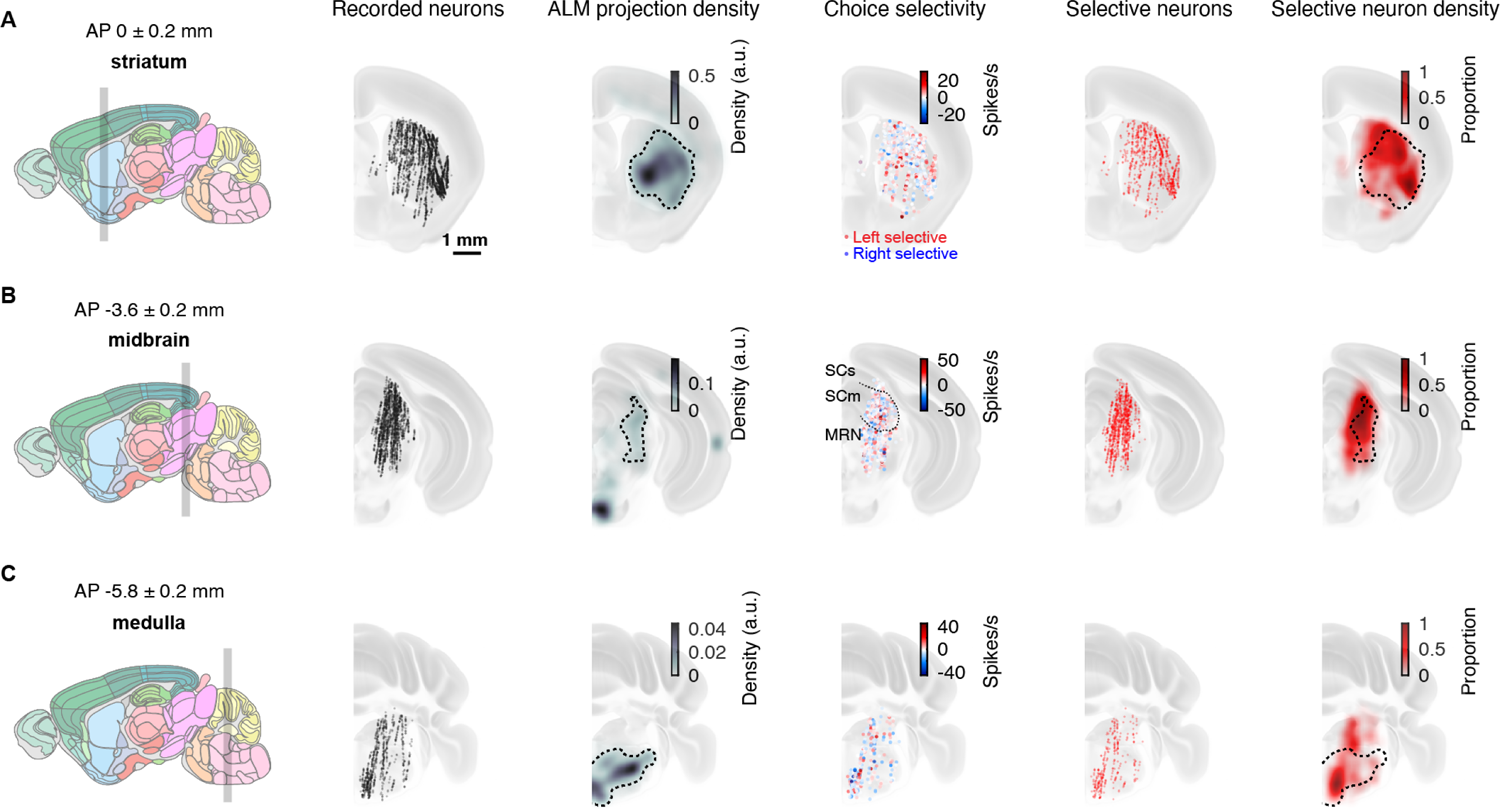
Spatial maps of choice selectivity. A. Left, sagittal view of the mouse brain. Gray shading, coronal section (AP 0.2 to −0.2 relative to bregma) through the striatum, corresponding to the remaining panels. Following panels, from left to right: recorded neurons in the striatum; ALM projection density (dashed line, contour indicating 90% of peak density); choice selectivity (individual neurons, spikes/s) (red/blue circles, left/right selective); choice selective neurons (rank sum test, P < 0.05); proportion of choice selective neurons, smoothed with a 3D Gaussian (sigma = 150 µm). B. Same as A for the midbrain. Gray shading, coronal section (AP −3.4 to −3.8 relative to bregma). The dotted line in the choice selectivity map represents borders for SCs, SCm, and MRN. C. Same as A for the hindbrain. Gray shading, coronal section (AP −5.6 to −6 relative to Bregma).

**Figure S5.**
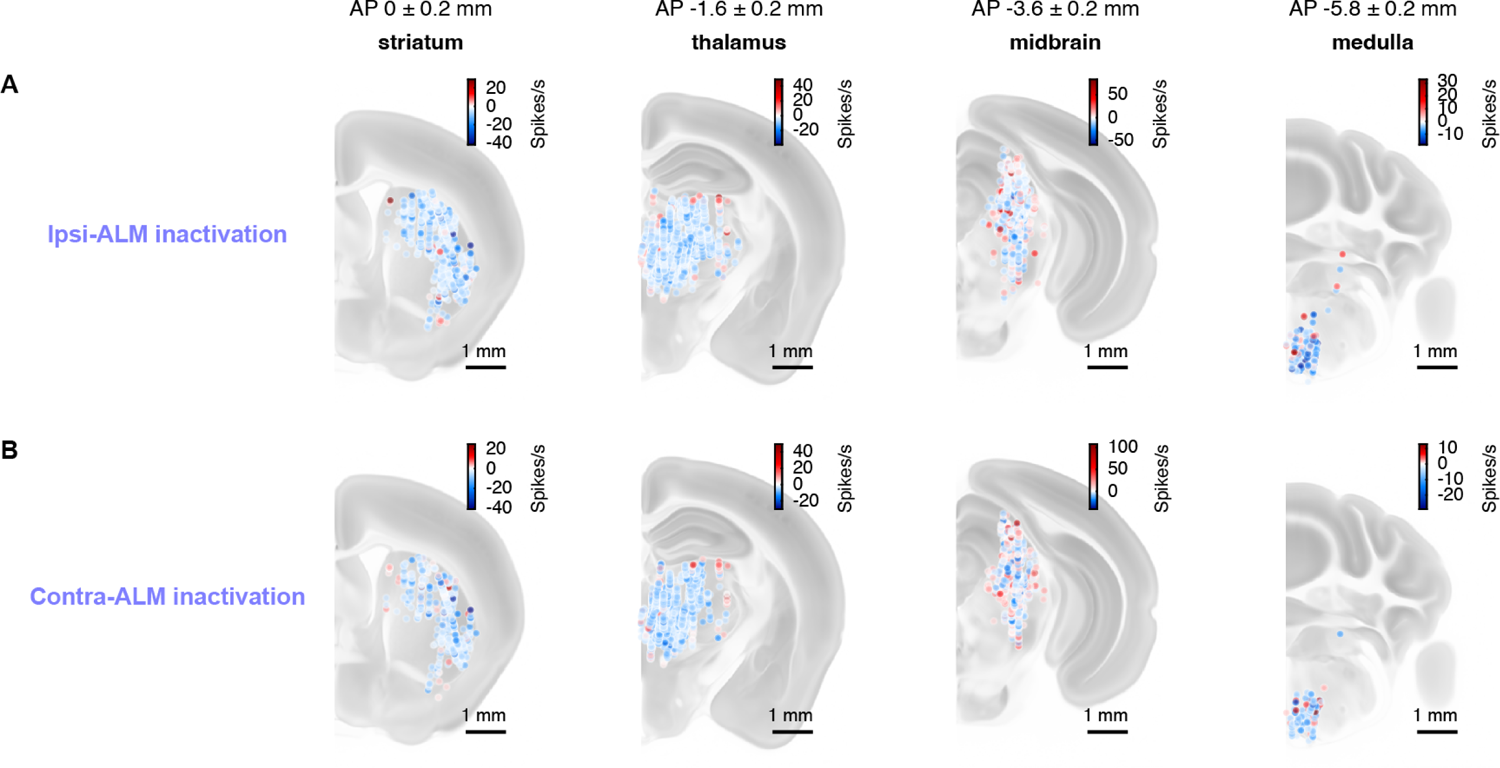
Spike rate change upon ALM inactivation. A. 2D coronal sections of spike rate change during ipsi-ALM photoinhibition. Blue dots, suppressed units. Red dots, activated units. B. 2D coronal sections of spike rate change during contra-ALM photoinhibition.

**Figure S6.**
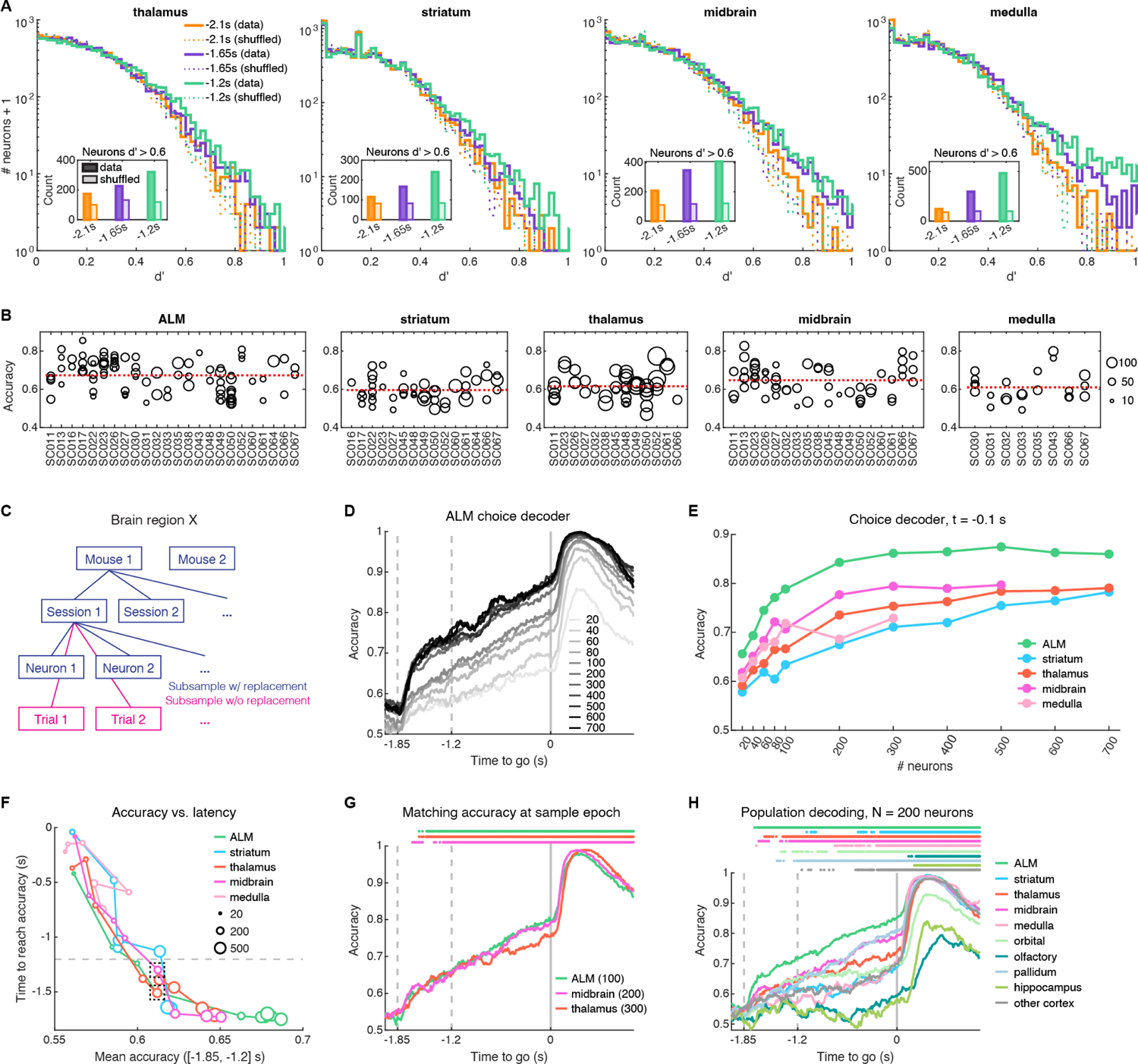
Decoding of choice across brain areas. A. Similar to Figure 6B, but here for subcortical areas. Distribution of single-neuron choice selectivity (i.e. d’, horizontal axis) in a 10,000-neuron pseudo-population from hierarchical bootstrapping. All neurons in subcortical areas (striatum, thalamus, midbrain, and medulla) were in ALM projection zones. B. Scatter plot showing single-session population choice decoding accuracy (vertical axis) along with mouse ID (horizontal axis labels) and number of neurons (size of the dots) in each recording. Each dot represents one recording with all neurons available in that brain area. The accuracy is the mean accuracy across −0.6 s to −0.1 s (i.e. late delay epoch). The red dotted line is the mean accuracy across all data points shown on each plot. This plot shows that naive averaging across sessions suffers from biases, such as unequal numbers of sessions per mouse, differing decoding accuracy across mice, various numbers of neurons per session, etc. C. A schematic of the process of hierarchical bootstrapping. D. Population choice decoding accuracy as a function of time in ALM, with various numbers of neurons (indicated by gray scale) subsampled by hierarchical bootstrapping. Standard error of the mean, not shown for clarity, is < 0.119. E. Population choice decoding accuracy at time −0.1 s (late delay epoch) as a function of number of neurons subsampled by hierarchical bootstrapping. Depending on the number of mice, number of sessions, and number of neurons recorded within each brain area, the upper limit of the number of neurons subsampled for the pseudo-population varies. Standard error of the mean, not shown for clarity, is < 0.10. F. As a larger number of neurons is subsampled in a pseudo-population by hierarchical bootstrapping, the mean population choice decoding accuracy over the sample epoch (horizontal axis) increases, and the time when choice can be decoded (vertical axis; see methods) shifts to earlier time-points. Size of dots indicates the number of neurons subsampled. The dashed box labeled three data points from ALM, thalamus, and midbrain that have similar mean accuracy across sample epoch and/or similar time to reach above-chance decodability, but different numbers of neurons. G. Population choice decoding accuracy as a function of time corresponding to the three data points in the dashed box in E. Bars at the top indicate times when the decoding accuracy of a brain area is higher than 0.5 with 95% confidence. Number of neurons for each area: ALM, 100 neurons; midrain, 200 neurons; thalamus, 300 neurons. H. Similar to Figure 6D, but including additional brain areas. ALM does not project to the orbital cortex, palladium, olfactory cortex, or hippocampus.

**Figure S7.**
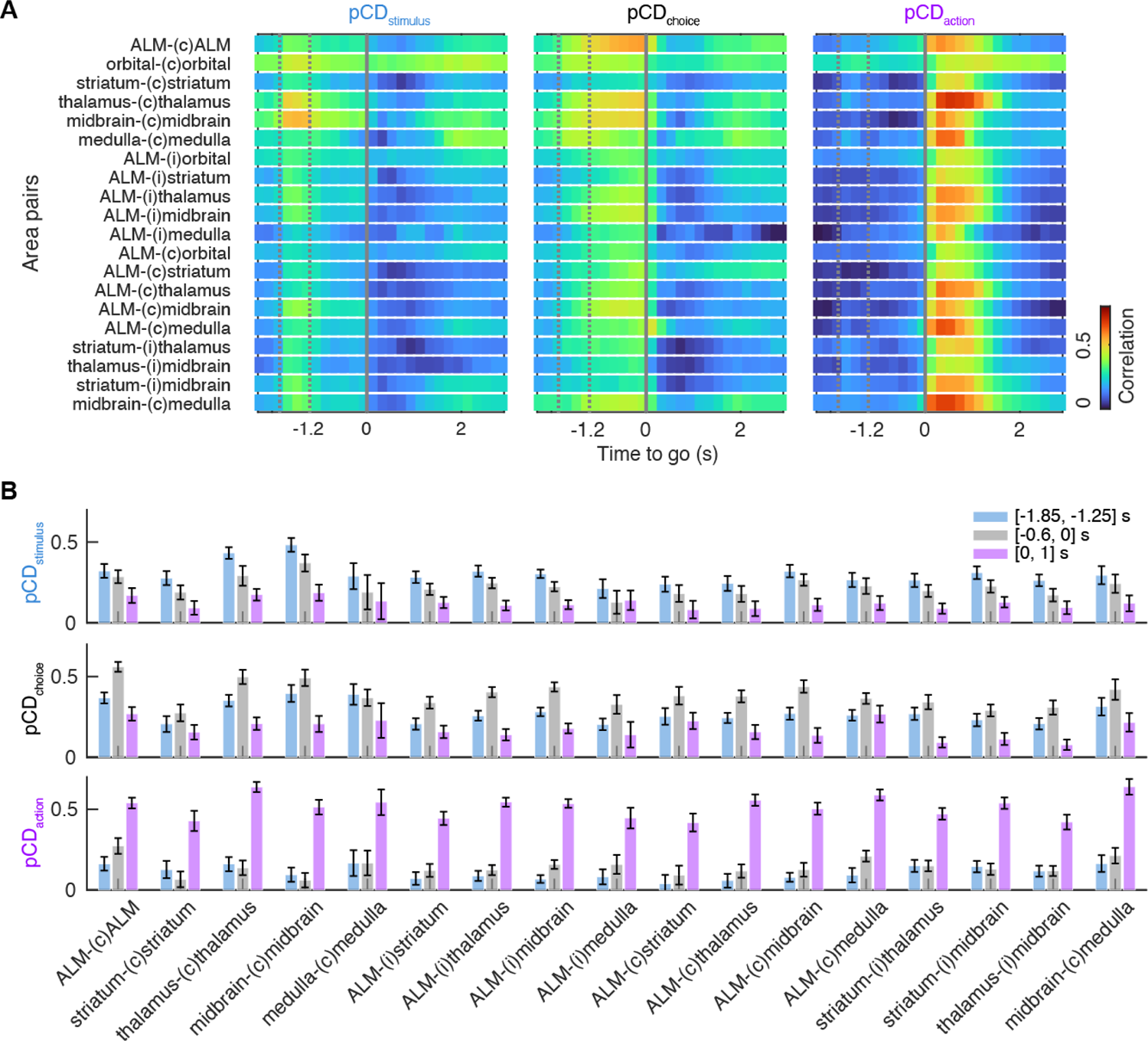
brain areas functional coupling. A. Time course of rank correlations of pCD_stimulus_, pCD_choice_ and pCD_action_ across major brain area pairs. (c): contralateral-; (i): ipsilateral-. B. Epoch-based quantification of pCD_stimulus_, pCD_choice_ and pCD_action_ correlation during sample epoch ([-1.85, −1.25] s), late delay epoch ([-0.6, 0] s) and response epoch ([0, 1] s) across major brain area pairs. Mean ± S.E.M. across sessions.

## Methods

### Animals and surgery

This study is based on data from 28 mice (Table 1), including twenty five VGAT-ChR2-EYFP (Jackson laboratory, JAX #014548), one C57BL/6J (JAX #000664), one Sst-IRES-Cre (JAX #013044) crossed with reporter mouse Ai32 (Rosa26-LSL-ChR2-EYFP, JAX #012569), and one Emx1-Cre (JAX #005628) crossed with R26-LNL-GtACR1-Fred-Kv2.1 reporter mouse (JAX #033089). See Table 2 for recordings made in each mouse.

All procedures were in accordance with protocols approved by the Janelia Research Campus Institutional Animal Care and Use Committee. Detailed information on water restriction has been published (Guo et al. 2014). Mice were housed in a 12:12 reverse light:dark cycle and performed behavioral tasks during the dark phase. Behavioral sessions lasted 1 to 2 h where mice received all their water (range, 0.3 to 1.5 mL). All surgical procedures were carried out aseptically. Mice were implanted with a titanium headpost and single housed. Recording well and craniotomy preparation for electrophysiology in head-restrained awake mice have been described in detail (<u>dx.doi.org/10.17504/protocols.io.9a8h2hw</u>). Buprenorphine (0.1 mg/kg, IP injection) was used for postoperative analgesia and Ketoprofen (5mg/kg, subcutaneous injection) was used at the time of surgery and postoperatively for two days. After head post implantation, all mice were allowed to recover for at least 3 days with free access to water before the start of water restriction. Craniotomies for recording were made after behavioral training.

### Behavior and video tracking

Mice performed an auditory delayed response task (Figure 1A) (Inagaki et al. 2018). The instruction stimuli during the sample epoch were pure tones played at one of two frequencies (3 kHz or 12 kHz). Each tone was played three times for 150 ms with 100 ms inter-tone intervals. The sample epoch was followed by a 1.2 s delay epoch. An auditory ‘Go’ cue (carrier frequency of 6 kHz with 360 Hz modulating frequency, 0.1 s duration) indicated the end of the delay epoch. Licking early during the sample/delay epoch triggered a replay of the epoch. During the response epoch (answer period: 1.5 s) mice reported the instruction by licking one of the two lick ports. Licking (consumption period: 1.5 s) the correct lick port triggered a small water reward (0.1 ∼ 0.2 ul). Licking the incorrect lick port triggered a timeout (1-3 s). After mice stopped licking for 1.5 s, the trial ended, followed by a 250 ms inter-trial-interval. Early lick trials and no response trials were excluded for analysis. Overall performance was computed as the fraction of correct control trials (i.e. no photostimulation), excluding any early lick trials. We selected experimental sessions for analysis based on following criteria: overall behavioral performance (> 65%), and at least 50 correct lick left and lick right trials each.

Two CMOS cameras (CM3-U3-13Y3M, FLIR) were used to track orofacial movements of the mouse under IR light illumination (940 nm LED). The cameras were equipped with 4-12 mm focal length lenses (12VM412ASIR, Tamron) and a pixel resolution of 71 µm. High-speed videos from a side view and a bottom view (Figure 1C) were acquired at 300 Hz using software FlyCapture (TELEDYNE FLIR). We trained DeepLabCut (Mathis et al. 2018) to track the movement of tongue, jaw and nose (Figure 1I).

### Multi-regional electrophysiological recordings

We designed a flexible manipulator system for multiple Neuropixels probe insertions (https://www.janelia.org/open-science/manipulator-system-for-multiple-neuropixels-probe-recordings). Three to four small craniotomies (diameter, 1 ∼ 1.5 mm) were made over target brain areas one day before the first recording session. Details of recordings made with multiple Neuropixels probes in head-restrained behaving animals have been published (dx.doi.org/10.17504/protocols.io.8tphwmn). Target regions, insertions angles, probe types, etc are summarized in Table 2. All coordinates are given with respect to Bregma (AP, anterior-posterior; ML, medial-lateral; DV, dorsal-ventral).

Extracellular spikes were recorded using Neuropixels 1.0 probes, 2.0 single-shank probes, and 2.0 multi-shank probes (J. J. Jun 2017; Steinmetz et al. 2021) (Table 2). Three to seven recordings were made from each craniotomy. Recording depth was inferred from manipulator readings (Sensapax uMp manipulators). After completion of probe insertion, brain tissue was allowed to settle for several minutes before recording. During recordings the craniotomies were immersed in saline.

Probes were configured and data were visualized and streamed to disk using the open-source software package SpikeGLX (https://billkarsh.github.io/SpikeGLX/). Data from up to five simultaneously recorded probes were acquired at 30 kHz. Sync waves (0.5 s duty cycle TTL pulses) were recorded across probes and on auxiliary channels for synchronization.

### Photoinhibition

Light from two 473 nm lasers (Coherent OBIS LS/LX) was used to illuminate the two hemispheres of ALM. The laser power was controlled by analog outputs controlled by a Bpod (Sanworks). Photoinactivation of ALM (centered on AP 2.5 mm; ML 1.5 mm) was performed through clear-skull cap implant (Guo et al. 2014) or via craniotomy by directing the laser over the skull (beam diameter, 1.8 mm at 4 sigma). Photoinhibition of the left-, right-ALM hemisphere, or both hemispheres was deployed on a subset of ∼25% randomly interleaved trials (N = 17 VGAT-ChR2-EYFP mice) (Z. V. Guo et al. 2014; Inagaki et al. 2018). To prevent the mice from distinguishing photostimulation trials from control trials using visual cues, a ‘masking flash’ (1 ms pulses at 10 Hz) was delivered using 470 nm LEDs in front of the mice throughout the entire trial. For silencing ALM, we stimulated cortical GABAergic neurons in VGAT-ChR2-EYFP mice. We used 40 Hz photostimulation with a sinusoidal temporal profile (5 mW average power per hemisphere) and a 100 ms linear ramping down during the laser offset to reduce rebound neuronal activity. This photo stimulus is expected to silence glutamatergic neurons in all of ALM (Nuo Li et al. 2019). We silenced ALM activity during the late delay epoch (last 0.5 s), including the 100 ms ramp-down. Thus, photoinhibition always ended before the ‘Go’ cue.

To analyze the effects of photoinhibition, units with at least 10 photostimulation trials were selected for analysis (Figure 5, Figure S5). Bilateral ALM photostimulation reduced behavior performance from 83.2% to 71.7% (N = 17 VGAT-ChR2-EYFP mice, n = 93 sessions). To exclude the effects of rebound population activity in the thalamus (Z. V. Guo et al. 2017) and other subcortical regions, we used a 20-120 ms time window after photo stimulation onset to measure the effect of ALM inactivation in downstream regions (Figure 5, Figure S5). For ALM itself, we used 500 ms (i.e. the entire inactivation period) to calculate spike rate reduction.

### Histology

To reconstruct recording locations, mice were perfused transcardially with PBS followed by 4% paraformaldehyde (PFA/0.1 M PBS). The brains were fixed overnight and transferred to SPiB solution for whole brain clearing. We used an aqueous-based clearing method ‘Uniclear’ for whole-brain delipidation and refractive index (RI) matching (dx.doi.org/10.17504/protocols.io.zndf5a6). Cleared brains were imaged with a light sheet microscope (Zeiss Z1) using a 5x objective (voxel size 1.2 µm x 1.2 µm x 6 µm). Tiled image stacks were registered and stitched in IMARIS. The whole brain image stacks were then downsampled 5x along x-y dimension respectively (yielding voxel size of 6 µm) in Image J and registered to the Allen Institute Common Coordinate Framework (CCF) of the mouse brain based on anatomical landmarks. Electrode tracks labeled with CM-DiI (Thermo-Fisher scientific) were used to determine recording locations. Nearby tracks were discernible from histology and individual tracks were carefully aligned using electrophysiological landmarks (L. D. Liu et al. 2020).

### Spike sorting and quality control

We developed a data pipeline for preprocessing, spike sorting and quality control (https://github.com/jenniferColonell/ecephys_spike_sorting). The extracellular recording traces were first band-pass filtered (butterworth acausal filter, 250 Hz - 9 kHz). We then sorted the dataset using both Kilosort2 (MATLAB version, <u>GitHub - jenniferColonell/ KS20_for_preprocessed_data: Release version of KS2 for preprocessed data</u>) (Stringer et al. 2019) and Kilosort2.5 (Python version, <u>GitHub - jenniferColonell/pykilosort: [WIP] Python port of Kilosort 2</u>) (Steinmetz et al. 2021). Both sorting methods gave similar results. In the analyses presented in the paper, we used the output from Kilosort2. Each output cluster was associated with a list of 15 quality metrics calculated to assess isolation and sorting quality (i.e. quality control) (https://github.com/AllenInstitute/ecephys_spike_sorting/tree/master/ecephys_spike_sorting/modules/quality_metrics) (Siegle et al. 2021).

The details of the spike sorting and quality control are described in an accompanying white paper (https://doi.org/10.25378/janelia.22154810.v1). In brief, we used 15 cluster quality metrics to train classifiers - one for each of the major five brain areas (cortex, striatum, thalamus, midbrain, and medulla). The classifiers were trained based on labels derived from manual curation. We curated Kilosort2 output (28 penetrations from five brain areas) using the open-source graphical user interface Phy (<u>GitHub - cortex-lab/ phy: phy: interactive visualization and manual spike sorting of large-scale ephys data</u>). Each cluster was labeled as a ‘good’ or ‘unlabeled’ unit. Annotators were blind to location, recording condition and trial type information. Based on the annotated labels of all clusters and associated quality metrics, we trained five region-specific classifiers using logistic regression. Applying the trained classifiers to the rest of the data set provided lists of units that were labeled as ‘good’ and were used for the analysis in this paper. We applied the cortex classifier to ALM and other cortex, hippocampus, olfactory and cortical subplate units; the striatum classifier to striatum and pallidum units; the thalamus classifier to thalamus and hypothalamus units; the midbrain classifier to midbrain and pons units; the medulla classifier to medulla and cerebellum units. This method yielded 8456 good units from ALM, 7510 from striatum, 12055 from thalamus, 7182 from midbrain and 2599 from medulla, etc. Overall, the data set consisted of 66,002 good units recorded across 173 behavioral sessions, from which 655 probe insertions were made. This corresponds to 24.4 % of clusters reported by Kilosort2.

## Data analysis

### ALM projection map

ALM projection zones were based on 212 reconstructed ALM neurons from the mouselight data set (Table 3) (Winnubst et al. 2019). ALM neurons were selected based on their soma location constrained by inactivation maps (soma CCF location within 100 µm) from previous loss of function experiments (Li et al. 2016). The relative strength of the projection to a target in the CCF was computed as fractional axonal density in the target (Figure S1D).

We used complete single neuron reconstructions to compute projection maps because the signal (axonal density) represents the actual fractional axonal length delivered to a particular location in the CCF. The projection map computed in this way corresponded qualitatively to projection maps based on fluorescence intensity after anterograde labeling with AAVs expressing fluorescent proteins (Figure S1 D, E) (Oh et al. 2014). However, the fluorescence intensity depends on axonal thickness, transport of the fluorescent protein, and local differences in scattering losses in the imaged tissue. In contrast, the projection map based on single neuron reconstructions is independent of these factors.

### Single neuron selectivity

Trial types differed with respect the auditory stimulus (‘stimulus’, high tone versus low tone), lick direction (‘choice’, lick left versus lick right before movement execution; ‘action’, lick left versus lick right) and reward (‘outcome’, correct/rewarded versus error/ unrewarded). We separately computed neuronal selectivity for each variable. Neurons were tested for significant stimulus (trial-type) selectivity during the sample epoch (0.65 s), choice selectivity during late delay epoch (last 0.6 s), action selectivity during response epoch (first 1.5 s after the ‘Go’ cue), and outcome selectivity post licking (2-3 s post ‘Go’ cue). In a subset of mice, licking persisted in occasional trials more than 2 s post ‘Go’ cue and therefore we cannot exclude that motor signals contribute to ‘outcome’ in some cases. To compute selectivity, four types of trials (1. stimulus left, choice left, <LL>; 2. stimulus left, choice right, <LR>; 3. stimulus right, choice left, <RL>; 4. stimulus right, choice right, <RR>) were pooled into two groups respectively (for stimulus, high vs. low; for choice and action, left vs. right; for outcome, rewarded vs. unrewarded). The spike counts of each type were first calculated over the time window of interest. Selectivity was then defined as the difference between the mean spike counts of the two groups. Neurons that significantly differentiated behavioral variables (Mann-Whitey U test, P < 0.05; Figure 2) during any one of the four trial epochs were scored as ‘selective’.

The four types of selectivity for each neuron *i* can be written as:

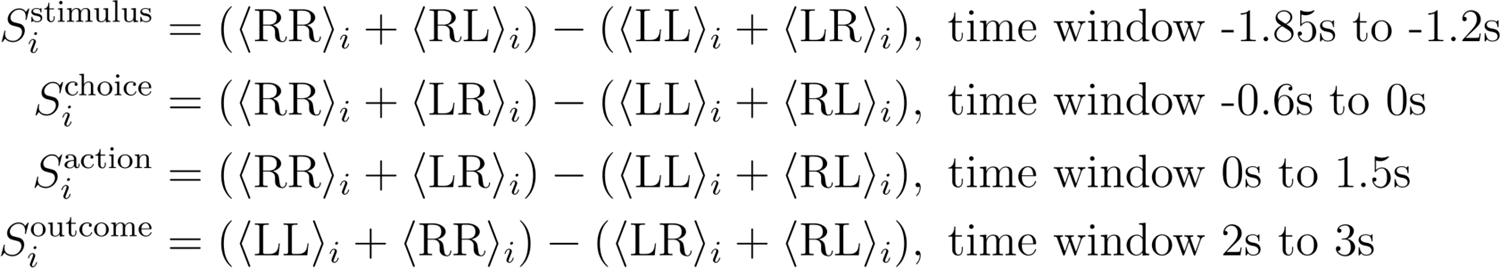

The equations above use the mean spike counts of each type of trial denoted by <stimulus choice> (stimulus choice = LL, RR, LR, RL), which is defined as:

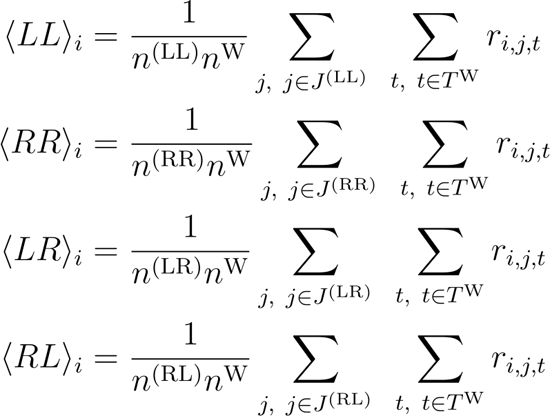

Where *r_i,j,t_* is the spike rate of neuron *i* in trial *j* at time *t*. *J*^(*stimulus choice*)^ represents the set of all trials with the specified stimulus and choice, and *_n_*(*stimulus choice*) is the total number of trials in *J*^(*stimulus choice*)^. *T^W^* represents the set of all time points within the time window (W) defined above when calculating a specific selectivity, and *n^W^* total number of time points in *T^W^*. is the

Selective neurons were classified into high tone preferring versus low tone preferring for stimulus, lick left preferring versus lick right preferring for choice and action, and correct preferring versus error preferring for outcome (Figure 2). To quantify the mean number of selective neurons in each brain area, the neuronal data set was sampled with replacement 1,000 times and the standard error of the mean was obtained by bootstrapping (Figure S2A, same for Figure S3B).

For the peri-stimulus time histograms (PSTHs), correct and incorrect trials were included and spike counts were binned by 1 ms and averaged over 200 ms using a box-car filter.

### Autoencoder of behavioral video

The architecture of the convolutional autoencoder was similar to BehaveNet (Batty et al. 2019). The encoder was composed of two residual blocks (He et al. 2016) and two fully connected layers. Each residual block consisted of four convolutional layers. The input image was resized into a 120×112 matrix. The output of the last convolutional layer was a 288-dimensional vector. This is followed by two fully connected layers consisting of rectified linear units that transform the output of the convolutional network into the output of the encoder, a 16-dimensional embedding vector. The decoder was a fully connected layer. We trained a separate autoencoder for each session. Sessions were screened for video artifacts and excluded if these were present.

### Predicting neural activity from behavioral video

We predicted neuron spike rates at time t using embedding vectors from the autoencoder concatenated across multiple time windows (t-34 ms, t-17 ms, t ms, t+17 ms, and t+34 ms) by linear regression with L2 penalty. Thus the dimensionality of the feature space was 80 (the 5 time steps times 16 embedding dimensions). An L2 penalty was used to regularize the model and inferred through cross-validation. Parameters of predictive models were the same across all time steps.

For Figure 3E-F, we used the Area Under the Curve (AUC) of the Receiver Operating Characteristic curve (ROC) to evaluate single neuron decodability of action before and after the subtraction of predictions from the embedding vectors, using only correct trials. The ROC is a non-parametric statistic calculated by sweeping a classification threshold from the smallest to the largest data value and measuring the increase of true and false positives. This sweep yields a curve of values. To condense this information to a single number the area under the curve is measured and is referred to as AUC. We developed a cross-test method when predicting animal behavior. We randomly split the dataset into 10 or 20 folds. With a specific classification threshold, we used one held-out fold for testing and the rest of the data for training and cross-validation. We looped over all held-out folds and thus generated predictions on the entire dataset on which the true and false positives (corresponding to one point on the ROC curve with the specific classification threshold) were evaluated. Neurons with low spike rates (under 2 Hz) were excluded from analysis. Changes in this threshold did not qualitatively modify results. Only neurons with an AUC > 0.6 based on spike rates were included in this analysis. Spike rates were binned in 40 ms time windows and 17ms time steps. In Figure 3F, neurons labeled as “no motor encoding” are those whose drop of explained variance ratio after the subtraction were lower than 0.05.

In Figure S3D, we re-trained models to predict spike rates from embedding vectors separately for each time bin width within which we binned spikes into spike rates. The explained variance ratio was evaluated accordingly.

### Definition of region subdivisions

For analyses across cortical depth and superior colliculus layers, we used layer annotations in the Allen Mouse Brain Common Coordinate Framework (CCFv3). Superficial layer ALM include units with annotations ‘layer 1’ and ‘layer 2/3’, and deep layer ALM include units with annotations ‘layer 5’ and ‘layer 6’. For analysis comparing dorsal and ventral striatum, we used annotation ‘Caudoputamen’ to identify dorsal striatum neurons. For subdivision in dorsal striatum, we defined dorsomedial striatum as units < 2.6 mm from the midline, and dorsolateral striatum as units > 2.6 mm from the midline (Figure 3D).

### Correlations between firing rate change during ALM inactivation and choice selectivity in ALM projection zones

To determine if the regression lines in Figure 5 are significantly different from zero, neurons were sampled with replacement 1,000 times for linear fitting. Zero lies outside of the 95% confidence interval, for the decreased and the increased group rejecting the null hypothesis that the coefficient is zero.

### Single-neuron choice selectivity (discriminability index d’)

We used the discriminability index d’ as a metric to evaluate single-neuron choice decodability (Figure 6A). For each neuron, we randomly subsampled 50 <LL> trials, 50 <RR> trials, 10 <LR> trials, and 10 <RL> trials (using the same <stimulus choice> convention as above) without replacement. The d’ at time t for neuron i was calculated using all the subsampled trials as:

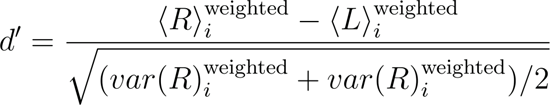

where the spike rate weighted mean 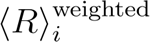 and weighted variance 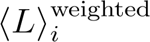 of choice R trials (for choice L trials they are defined similarly) are calculated as:

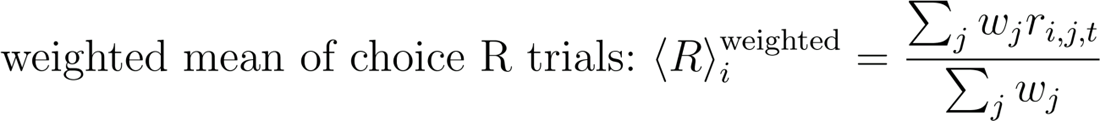

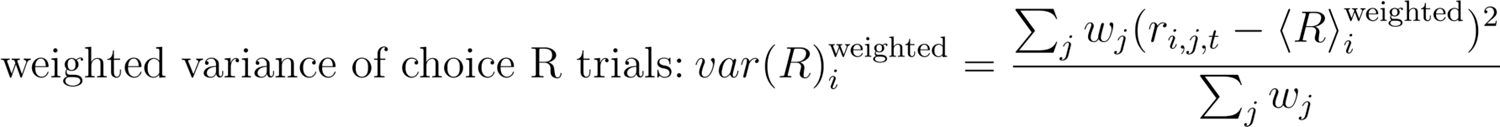

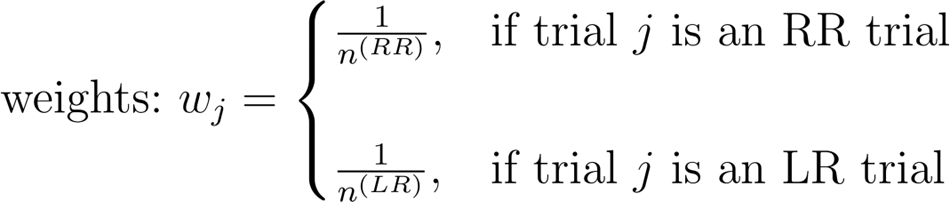

And the variables are defined in the same way as above mentioned. The spike rate of neuron *i* in trial *j* at time *t, r_i,j,t_*, was calculated with causal sliding time windows of 200 ms and a step size of 10 ms. The d’ evaluates the choice decodability while balancing the unequal numbers of correct and error trials within trials with the same choice, which is a unique advantage over other metrics such as AUC of the ROC. This enables d’ to differentiate choice decodability from stimulus decodability.

### Hierarchical bootstrapping

To minimize sampling bias, we used hierarchical bootstrapping to construct pseudo-populations (Figures 6B-D). Recordings with fewer than 10 neurons in a target brain area were rejected. For each brain area, we randomly sampled 5 mice with replacement from all mice with recordings from that region. For each sampled mouse, 2 sessions belonging to that mouse were sampled with replacement. Within each session, we randomly sampled 50 <LL> trials, 50 <RR> trials, 10 <LR> trials, and 10 <RL> trials (using the same <stimulus choice> convention as above; sessions with fewer trials of any type were discarded) without replacement, and we also sampled 10 (or 20) neurons with replacement. As a result, with each round of hierarchical bootstrapping, we constructed a pseudo-population of 100 (or 200) neurons. We repeated the hierarchical bootstrapping process 100 times. Varying the number of neurons subsampled within a reasonable range did not change the conclusions.

To calculate the distribution of single-neuron choice selectivity (Figures 6B-C), we subsampled 10 neurons from each session, resulting in a pseudo-population of 100 neurons in one round of hierarchical bootstrapping. Then we repeated the process 100 times and concatenated all the neurons together, constructing a large pseudo-population of 10,000 neurons. The distribution and fraction of selective neurons were calculated based on this large pseudo-population.

For population choice decoding (Figures 6D), we subsampled 20 neurons from each session, resulting in a pseudo-population of 200 neurons in one round of hierarchical bootstrapping. We then separately trained decoders on each round of hierarchical bootstrapping using the pseudo-population of 200 neurons and calculated the mean and standard error of the mean of the accuracy traces across 100 repetitions.

### Distribution of single-neuron choice selectivity

At each time point, we calculated the distribution of single-neuron choice selectivity (Figure 6B) or fraction of choice-selective neurons (Figure 6C) defined as the fraction of neurons with d’ > threshold (0.6). Varying the threshold in a reasonable range only changed the overall range of the fraction values and did not affect the conclusions.

As a comparison, we also performed a null distribution as a benchmark (labeled as “shuffled”). To calculate the shuffle distribution, the trial IDs of the spike rates were randomly shuffled, resulting in a mismatch between spike rates and other variables (trial types and choice). Compared to the null distribution, the real neuron populations showed a larger tail in highly-selective neurons (Figure 6B) and an increasing number of choice selective neurons across the sample epoch (Figure 6C).

### Population choice decoder

Population choice decoders were calculated using spike rates with causal sliding time windows of 200 ms and a step size of 10 ms. For each brain area, we trained a separate population decoder using logistic regression at each time point using the pseudo-population in each round of hierarchical bootstrapping (Figure 6D). We performed nested 5-fold cross validation to train and evaluate the decoder accuracy. In the outer loop, each time one fold (i.e. 20%) of the trials was used as test trials while keeping the ratio of the number of <LL>, <RR>, <LR>, and <RL> trials unchanged. In the inner loop, within the rest four folds (i.e. 80%) of the trials, we used 4-fold cross validation to select the best regularization parameter. The decoder was then trained on all the non-test trials using the best regularization, and evaluated on the test trials for its final performance in one iteration of the outer loop. The overall accuracy of this pseudo-population is the mean accuracy over all iterations of the outer loop.

In Figure S6F, the vertical axis is the time when choice can be decoded above chance, which is defined as the earliest time t that satisfies the condition that at least 9 out of 10 time bins following this time bin t have above-chance choice decodability.

### Coding direction (**CD**) analysis

To analyze the relationship between the population activity of pairs of brain areas, we restricted analysis to recording sessions with more than 10 neurons recorded simultaneously for each area. We projected population activity to distinct directions in activity space along which population activity was selective for stimulus, choice or action. For a population of *n* neurons, we defined a set of orthogonal directions in activity space (*n* x 1 vectors). In the n dimensional activity space these vectors maximally separated the response vectors in lick-left and lick-right trials at specific times of the task, and were termed the coding directions (**CD**). For each lick direction we computed the average spike counts, ^→^*x* _lick left_ and ^→^*x* _lick right_, *n* x 1 response vectors that described the population response at one time point. The direction of the difference in the mean response vectors at time point t is ^→^*w* _t_ = ^→^*x* _lick right_ - ^→^*x* _lick left_. Stimulus selectivity vectors were stable during the sample epoch. We averaged the ^→^*w* _t_ values in the first 0.6 s of the sample epoch to obtain stimulus **CD** (**CD_stimulus_**). Choice selectivity vectors were stable during the delay epoch. We therefore averaged the ^→^*w* _t_ values in the last 0.6 s of the delay epoch to obtain choice **CD** (**CD_choice_**). We defined action **CD** (**CD_action_**) by averaging ^→^*w* _t_ values in the first 1 s after the ‘Go’ cue. **CD_stimulus_** and **CD_action_** were rotated using the Gram-Schmidt process to be fully orthogonal to **CD_choice_**. To project population activity along a **CD** we used independent control trials from the trials used to compute the **CD** (leave-one-trial-out method). For each trial, we computed the spike counts for each neuron, **x** (*n* x 1 response vector), at each time point. Projections onto a **CD** (i.e. pCD_stimulus_, pCD_choice_ and pCD_action_) were obtained as **CD**^T^**x**.

To examine correlated stimulus, choice, and action encoding between pairs of brain areas, for each session we took the single trial trajectory values at each time bin (200 ms) and calculated Spearman’s rank correlation between trajectory values of region 1 and region 2. We calculated rank correlation using both trial types of the session (Figure 7, Figure S7).

### Statistics

The sample sizes are similar to sample sizes used in the field (more than 300 units per brain area). No statistical methods were used to determine sample size. Trial types were randomly determined by a computer program during behavior. During manual curation, annotators cannot tell the trial type, and therefore were blind to conditions. Error bars indicate mean ± S.E.M. unless otherwise stated.

