## Supplementary material for "Brain-wide neural activity underlying memory-guided movement": Tables

**Table 1 - Mice information**

| Animal | Strain | Sex | Animal source | Age at recording |
|---|---|---|---|---|
| SC011 | VGAT-ChR2-EYFP | M | Jackson Lab | 4 months |
| SC013 | SOM-IRES-Cre x Ai32 | M | Jackson Lab | 7 months |
| SC015 | VGAT-ChR2-EYFP | M | Jackson Lab | 4 months |
| SC016 | VGAT-ChR2-EYFP | M | Jackson Lab | 4 months |
| SC017 | VGAT-ChR2-EYFP | M | Jackson Lab | 4 months |
| SC022 | VGAT-ChR2-EYFP | M | Jackson Lab | 4 months |
| SC023 | VGAT-ChR2-EYFP | M | Jackson Lab | 3 months |
| SC026 | VGAT-ChR2-EYFP | M | Jackson Lab | 3 months |
| SC027 | VGAT-ChR2-EYFP | M | Jackson Lab | 3 months |
| SC030 | VGAT-ChR2-EYFP | M | Jackson Lab | 4 months |
| SC031 | VGAT-ChR2-EYFP | M | Jackson Lab | 4 months |
| SC032 | VGAT-ChR2-EYFP | M | Jackson Lab | 5 months |
| SC033 | VGAT-ChR2-EYFP | M | Jackson Lab | 4 months |
| SC035 | VGAT-ChR2-EYFP | M | Jackson Lab | 6 months |
| SC038 | VGAT-ChR2-EYFP | M | Jackson Lab | 5 months |
| SC043 | C57Bl/6Crl | M | Jackson Lab | 4 months |
| SC045 | Emx1-Cre x R26-LNL-GtACR1-Fred-Kv2.1 | M | Janelia | 5 months |
| SC048 | VGAT-ChR2-EYFP | M | Jackson Lab | 4 months |
| SC049 | VGAT-ChR2-EYFP | F | Jackson Lab | 5 months |
| SC050 | VGAT-ChR2-EYFP | F | Jackson Lab | 7 months |
| SC052 | VGAT-ChR2-EYFP | F | Jackson Lab | 4 months |
| SC053 | VGAT-ChR2-EYFP | M | Jackson Lab | 5 months |
| SC060 | VGAT-ChR2-EYFP | M | Jackson Lab | 4 months |
| SC061 | VGAT-ChR2-EYFP | M | Jackson Lab | 4 months |
| SC064 | VGAT-ChR2-EYFP | M | Jackson Lab | 5 months |
| SC065 | VGAT-ChR2-EYFP | M | Jackson Lab | 5 months |
| SC066 | VGAT-ChR2-EYFP | M | Jackson Lab | 4 months |
| SC067 | VGAT-ChR2-EYFP | M | Jackson Lab | 4 months |

**Table 2 - Insertions information**

| Animal | Target area | Hemi-sphere | Session date | Insertion no. | Probe type | AP location (µm) | ML location (µm) | depth (µm) | theta (deg) | phi (deg) | beta (deg) | Imec probe index |
|---|---|---|---|---|---|---|---|---|---|---|---|---|
| SC011 | ALM | left | 2019-02-19 | 1 | 3B 1.0 | 2500 | -1500 | -2900 | 15 | 135 | -135 | 0 |
| SC011 | Thalamus | left | 2019-02-19 | 2 | 3B 1.0 | -1700 | -1300 | -4400 | 15 | 180 | 90 | 1 |
| SC011 | Midbrain | left | 2019-02-19 | 3 | 3B 1.0 | -3800 | -1700 | -4500 | 15 | 0 | 90 | 2 |
| SC011 | ALM | left | 2019-02-20 | 1 | 3B 1.0 | 2500 | -1500 | -2900 | 15 | 135 | -135 | 0 |
| SC011 | Thalamus | left | 2019-02-20 | 2 | 3B 1.0 | -1700 | -1300 | -4700 | 10 | 180 | 90 | 1 |
| SC011 | Midbrain | left | 2019-02-20 | 3 | 3B 1.0 | -3800 | -1700 | -4700 | 5 | 0 | 90 | 2 |
| SC011 | ALM | left | 2019-02-21 | 1 | 3B 1.0 | 2500 | -1500 | -2900 | 15 | 135 | -135 | 0 |
| SC011 | Thalamus | left | 2019-02-21 | 2 | 3B 1.0 | -1700 | -1300 | -4700 | 10 | 180 | 90 | 1 |
| SC011 | Midbrain | left | 2019-02-21 | 3 | 3B 1.0 | -3800 | -1700 | -4550 | 5 | 0 | 90 | 2 |
| SC011 | ALM | left | 2019-02-22 | 1 | 3B 1.0 | 2500 | -1500 | -2900 | 15 | 135 | -135 | 0 |
| SC011 | Thalamus | left | 2019-02-22 | 2 | 3B 1.0 | -1700 | -1300 | -4800 | 10 | 180 | 90 | 1 |
| SC011 | Midbrain | left | 2019-02-22 | 3 | 3B 1.0 | -3800 | -1700 | -4500 | 5 | 0 | 90 | 2 |
| SC011 | ALM | left | 2019-02-23 | 1 | 3B 1.0 | 2500 | -1500 | -2900 | 15 | 135 | -135 | 0 |
| SC011 | ALM | right | 2019-02-23 | 2 | 3B 1.0 | 2500 | 1500 | -2950 | 15 | 45 | -45 | 1 |
| SC011 | Midbrain | right | 2019-02-23 | 3 | 3B 1.0 | -3900 | 1000 | -4600 | 10 | 180 | 90 | 2 |
| SC011 | Thalamus | right | 2019-02-23 | 4 | 3B 1.0 | -1700 | 2000 | -4800 | 15 | 0 | 90 | 3 |
| SC011 | ALM | right | 2019-02-24 | 1 | 3B 1.0 | 2500 | 1500 | -2900 | 15 | 45 | -45 | 0 |
| SC011 | Midbrain | right | 2019-02-24 | 2 | 3B 1.0 | -3900 | 1000 | -4600 | 10 | 180 | 90 | 1 |
| SC011 | Thalamus | right | 2019-02-24 | 3 | 3B 1.0 | -1700 | 2000 | -4800 | 15 | 0 | 90 | 2 |
| SC011 | ALM | right | 2019-02-25 | 1 | 3B 1.0 | 2500 | 1500 | -2900 | 15 | 45 | -45 | 0 |

| | | | | | | | | | | | | |
|---|---|---|---|---|---|---|---|---|---|---|---|---|
| SC011 | Midbrain | right | 2019-02-25 | 2 | 3B 1.0 | -3900 | 1000 | -4700 | 10 | 180 | 90 | 1 |
| SC011 | Thalamus | right | 2019-02-25 | 3 | 3B 1.0 | -1700 | 2000 | -4800 | 15 | 0 | 90 | 2 |
| SC011 | ALM | right | 2019-02-26 | 1 | 3B 1.0 | 2500 | 1500 | -2900 | 15 | 45 | -45 | 0 |
| SC011 | Midbrain | right | 2019-02-26 | 2 | 3B 1.0 | -3900 | 1000 | -4700 | 10 | 180 | 90 | 1 |
| SC011 | Thalamus | right | 2019-02-26 | 3 | 3B 1.0 | -1700 | 2000 | -4800 | 15 | 0 | 90 | 2 |
| SC013 | ALM | left | 2019-05-13 | 1 | 3B 1.0 | 2500 | -1500 | -2900 | 15 | 135 | -135 | 0 |
| SC013 | ALM | right | 2019-05-13 | 2 | 3B 1.0 | 2500 | 1500 | -2900 | 15 | 45 | -45 | 1 |
| SC013 | Midbrain | left | 2019-05-13 | 3 | 3B 1.0 | -4000 | -1800 | -4700 | 10 | 180 | 120 | 2 |
| SC013 | Midbrain | right | 2019-05-13 | 4 | 3B 1.0 | -4000 | 1800 | -4700 | 10 | 0 | 60 | 3 |
| SC013 | ALM | left | 2019-05-14 | 1 | 3B 1.0 | 2500 | -1500 | -2900 | 15 | 135 | -135 | 0 |
| SC013 | ALM | right | 2019-05-14 | 2 | 3B 1.0 | 2500 | 1500 | -2900 | 15 | 45 | -45 | 1 |
| SC013 | Midbrain | left | 2019-05-14 | 3 | 3B 1.0 | -4000 | -1800 | -4600 | 10 | 180 | 120 | 2 |
| SC013 | Midbrain | right | 2019-05-14 | 4 | 3B 1.0 | -4000 | 1800 | -4600 | 10 | 0 | 60 | 3 |
| SC013 | ALM | left | 2019-05-15 | 1 | 3B 1.0 | 2500 | -1500 | -2900 | 15 | 135 | -135 | 0 |
| SC013 | ALM | right | 2019-05-15 | 2 | 3B 1.0 | 2500 | 1500 | -2900 | 15 | 45 | -45 | 1 |
| SC013 | Midbrain | left | 2019-05-15 | 3 | 3B 1.0 | -4000 | -1800 | -4700 | 10 | 180 | 120 | 2 |
| SC013 | Midbrain | right | 2019-05-15 | 4 | 3B 1.0 | -4000 | 1800 | -4700 | 10 | 0 | 60 | 3 |
| SC013 | ALM | left | 2019-05-16 | 1 | 3B 1.0 | 2500 | -1500 | -2900 | 15 | 135 | -135 | 0 |
| SC013 | ALM | right | 2019-05-16 | 2 | 3B 1.0 | 2500 | 1500 | -2900 | 15 | 45 | -45 | 1 |
| SC013 | Midbrain | left | 2019-05-16 | 3 | 3B 1.0 | -4000 | -1800 | -4750 | 10 | 180 | 120 | 2 |
| SC013 | Midbrain | right | 2019-05-16 | 4 | 3B 1.0 | -4000 | 1800 | -4750 | 10 | 0 | 60 | 3 |
| SC013 | ALM | left | 2019-05-17 | 1 | 3B 1.0 | 2500 | -1500 | -2950 | 15 | 135 | -135 | 0 |
| SC013 | ALM | right | 2019-05-17 | 2 | 3B 1.0 | 2500 | 1500 | -2950 | 15 | 45 | -45 | 1 |
| SC013 | Midbrain | left | 2019-05-17 | 3 | 3B 1.0 | -4000 | -1800 | -4800 | 10 | 180 | 120 | 2 |

| | | | | | | | | | | | | |
|---|---|---|---|---|---|---|---|---|---|---|---|---|
| SC013 | Midbrain | right | 2019-05-17 | 4 | 3B 1.0 | -4000 | 1800 | -4800 | 10 | 0 | 60 | 3 |
| SC015 | ALM | left | 2019-02-07 | 1 | 3B 1.0 | 2500 | -1500 | -2600 | 15 | 135 | -135 | 0 |
| SC015 | ALM | right | 2019-02-07 | 2 | 3B 1.0 | 2500 | 1500 | -2600 | 15 | 45 | -45 | 1 |
| SC015 | Striatum | left | 2019-02-07 | 3 | 3B 1.0 | 0 | -2500 | -4200 | 10 | 180 | 120 | 2 |
| SC015 | Striatum | right | 2019-02-07 | 4 | 3B 1.0 | 0 | 2500 | -4200 | 10 | 0 | 60 | 3 |
| SC015 | ALM | left | 2019-02-08 | 1 | 3B 1.0 | 2500 | -1500 | -2700 | 15 | 135 | -135 | 0 |
| SC015 | ALM | right | 2019-02-08 | 2 | 3B 1.0 | 2500 | 1500 | -2700 | 15 | 45 | -45 | 1 |
| SC015 | Striatum | left | 2019-02-08 | 3 | 3B 1.0 | 0 | -2500 | -4300 | 10 | 180 | 120 | 2 |
| SC015 | Striatum | right | 2019-02-08 | 4 | 3B 1.0 | 0 | 2500 | -4000 | 10 | 0 | 60 | 3 |
| SC015 | Striatum | left | 2019-02-09 | 1 | 3B 1.0 | 0 | -2500 | -4200 | 10 | 180 | 120 | 0 |
| SC015 | Striatum | right | 2019-02-09 | 2 | 3B 1.0 | 0 | 2500 | -4200 | 10 | 0 | 60 | 1 |
| SC015 | ALM | left | 2019-02-10 | 1 | 3B 1.0 | 2500 | -1500 | -2700 | 15 | 135 | -135 | 0 |
| SC015 | ALM | right | 2019-02-10 | 2 | 3B 1.0 | 2500 | 1500 | -2700 | 15 | 45 | -45 | 1 |
| SC015 | Striatum | left | 2019-02-10 | 3 | 3B 1.0 | 0 | -2500 | -4200 | 10 | 180 | 120 | 2 |
| SC015 | Striatum | right | 2019-02-10 | 4 | 3B 1.0 | 0 | 2500 | -4200 | 10 | 0 | 60 | 3 |
| SC016 | ALM | left | 2019-02-11 | 1 | 3B 1.0 | 2500 | -1500 | -2700 | 15 | 135 | -135 | 0 |
| SC016 | ALM | right | 2019-02-11 | 2 | 3B 1.0 | 2500 | 1500 | -2700 | 15 | 45 | -45 | 1 |
| SC016 | Striatum | left | 2019-02-11 | 3 | 3B 1.0 | 0 | -2500 | -4300 | 10 | 180 | 120 | 2 |
| SC016 | Striatum | right | 2019-02-11 | 4 | 3B 1.0 | 0 | 2500 | -4300 | 10 | 0 | 60 | 3 |
| SC016 | ALM | left | 2019-02-12 | 1 | 3B 1.0 | 2500 | -1500 | -2700 | 15 | 135 | -135 | 0 |
| SC016 | ALM | right | 2019-02-12 | 2 | 3B 1.0 | 2500 | 1500 | -2700 | 15 | 45 | -45 | 1 |
| SC016 | Striatum | left | 2019-02-12 | 3 | 3B 1.0 | 0 | -2500 | -4300 | 10 | 180 | 120 | 2 |
| SC016 | Striatum | right | 2019-02-12 | 4 | 3B 1.0 | 0 | 2500 | -4300 | 10 | 0 | 60 | 3 |
| SC016 | ALM | right | 2019-02-13 | 1 | 3B 1.0 | 2500 | 1500 | -2800 | 15 | 45 | -45 | 0 |

| | | | | | | | | | | | | |
|---|---|---|---|---|---|---|---|---|---|---|---|---|
| SC016 | Striatum | left | 2019-02-13 | 2 | 3B 1.0 | 0 | -2500 | -4400 | 10 | 180 | 120 | 1 |
| SC016 | Striatum | right | 2019-02-13 | 3 | 3B 1.0 | 0 | 2500 | -4400 | 10 | 0 | 60 | 2 |
| SC016 | ALM | left | 2019-02-14 | 1 | 3B 1.0 | 2500 | -1500 | -2900 | 15 | 135 | -135 | 0 |
| SC016 | ALM | right | 2019-02-14 | 2 | 3B 1.0 | 2500 | 1500 | -2900 | 15 | 45 | -45 | 1 |
| SC016 | Striatum | right | 2019-02-14 | 3 | 3B 1.0 | 0 | 2500 | -4400 | 10 | 0 | 60 | 2 |
| SC017 | ALM | left | 2019-02-13 | 1 | 3B 1.0 | 2500 | -1500 | -2800 | 15 | 135 | -135 | 0 |
| SC017 | ALM | right | 2019-02-13 | 2 | 3B 1.0 | 2500 | 1500 | -2800 | 15 | 45 | -45 | 1 |
| SC017 | Striatum | left | 2019-02-13 | 3 | 3B 1.0 | 0 | -2500 | -4300 | 10 | 180 | 120 | 2 |
| SC017 | Striatum | right | 2019-02-13 | 4 | 3B 1.0 | 0 | 2500 | -4300 | 10 | 0 | 60 | 3 |
| SC017 | ALM | left | 2019-02-14 | 1 | 3B 1.0 | 2500 | -1500 | -2850 | 15 | 135 | -135 | 0 |
| SC017 | ALM | right | 2019-02-14 | 2 | 3B 1.0 | 2500 | 1500 | -2850 | 15 | 45 | -45 | 1 |
| SC017 | Striatum | left | 2019-02-14 | 3 | 3B 1.0 | 0 | -2500 | -4400 | 15 | 180 | 120 | 2 |
| SC017 | Striatum | right | 2019-02-14 | 4 | 3B 1.0 | 0 | 2500 | -4400 | 15 | 0 | 60 | 3 |
| SC017 | ALM | left | 2019-02-15 | 1 | 3B 1.0 | 2500 | -1500 | -2900 | 15 | 135 | -135 | 0 |
| SC017 | ALM | right | 2019-02-15 | 2 | 3B 1.0 | 2500 | 1500 | -2900 | 15 | 45 | -45 | 1 |
| SC017 | Striatum | left | 2019-02-15 | 3 | 3B 1.0 | 0 | -2500 | -4400 | 15 | 180 | 120 | 2 |
| SC017 | Striatum | right | 2019-02-15 | 4 | 3B 1.0 | 0 | 2500 | -4400 | 15 | 0 | 60 | 3 |
| SC017 | ALM | left | 2019-02-17 | 1 | 3B 1.0 | 2500 | -1500 | -2900 | 15 | 135 | -135 | 0 |
| SC017 | ALM | right | 2019-02-17 | 2 | 3B 1.0 | 2500 | 1500 | -2900 | 15 | 45 | -45 | 1 |
| SC017 | Striatum | left | 2019-02-17 | 3 | 3B 1.0 | 0 | -2500 | -4200 | 15 | 180 | 120 | 2 |
| SC017 | Striatum | right | 2019-02-17 | 4 | 3B 1.0 | 0 | 2500 | -4200 | 15 | 0 | 60 | 3 |
| SC022 | ALM | left | 2019-02-27 | 1 | 3B 1.0 | 2500 | -1500 | -2900 | 15 | 135 | -135 | 0 |
| SC022 | ALM | right | 2019-02-27 | 2 | 3B 1.0 | 2500 | 1500 | -2900 | 15 | 45 | -45 | 1 |
| SC022 | Striatum | left | 2019-02-27 | 3 | 3B 1.0 | 0 | -2800 | -4600 | 10 | 180 | 120 | 2 |

| | | | | | | | | | | | | |
|---|---|---|---|---|---|---|---|---|---|---|---|---|
| SC022 | Striatum | right | 2019-02-27 | 4 | 3B 1.0 | 0 | 2800 | -4600 | 10 | 0 | 60 | 3 |
| SC022 | ALM | left | 2019-02-28 | 1 | 3B 1.0 | 2500 | -1500 | -2900 | 15 | 135 | -135 | 0 |
| SC022 | ALM | right | 2019-02-28 | 2 | 3B 1.0 | 2500 | 1500 | -2900 | 15 | 45 | -45 | 1 |
| SC022 | Striatum | left | 2019-02-28 | 3 | 3B 1.0 | 0 | -2800 | -4600 | 10 | 180 | 120 | 2 |
| SC022 | Striatum | right | 2019-02-28 | 4 | 3B 1.0 | 0 | 2800 | -4600 | 10 | 0 | 60 | 3 |
| SC022 | ALM | left | 2019-03-01 | 1 | 3B 1.0 | 2500 | -1500 | -2900 | 15 | 135 | -135 | 0 |
| SC022 | ALM | right | 2019-03-01 | 2 | 3B 1.0 | 2500 | 1500 | -2900 | 15 | 45 | -45 | 1 |
| SC022 | Striatum | left | 2019-03-01 | 3 | 3B 1.0 | 0 | -2800 | -4650 | 10 | 180 | 120 | 2 |
| SC022 | Striatum | right | 2019-03-01 | 4 | 3B 1.0 | 0 | 2800 | -4650 | 10 | 0 | 60 | 3 |
| SC022 | ALM | left | 2019-03-02 | 1 | 3B 1.0 | 2500 | -1500 | -2900 | 15 | 135 | -135 | 0 |
| SC022 | ALM | right | 2019-03-02 | 2 | 3B 1.0 | 2500 | 1500 | -2900 | 15 | 45 | -45 | 1 |
| SC022 | Striatum | left | 2019-03-02 | 3 | 3B 1.0 | 0 | -2800 | -4600 | 10 | 180 | 120 | 2 |
| SC022 | Striatum | right | 2019-03-02 | 4 | 3B 1.0 | 0 | 2800 | -4600 | 10 | 0 | 60 | 3 |
| SC022 | ALM | left | 2019-03-03 | 1 | 3B 1.0 | 2500 | -1500 | -2900 | 15 | 135 | -135 | 0 |
| SC022 | ALM | right | 2019-03-03 | 2 | 3B 1.0 | 2500 | 1500 | -2900 | 15 | 45 | -45 | 1 |
| SC022 | Striatum | left | 2019-03-03 | 3 | 3B 1.0 | 0 | -2800 | -4500 | 10 | 180 | 120 | 2 |
| SC022 | Striatum | right | 2019-03-03 | 4 | 3B 1.0 | 0 | 2800 | -4500 | 10 | 0 | 60 | 3 |
| SC023 | ALM | left | 2019-05-30 | 1 | 3B 1.0 | 2500 | -1500 | -2900 | 15 | 135 | -135 | 0 |
| SC023 | ALM | right | 2019-05-30 | 2 | 3B 1.0 | 2500 | 1500 | -2900 | 15 | 45 | -45 | 1 |
| SC023 | Midbrain | left | 2019-05-30 | 3 | 3B 1.0 | -3800 | -1800 | -4600 | 10 | 180 | 120 | 2 |
| SC023 | Midbrain | right | 2019-05-30 | 4 | 3B 1.0 | -3800 | 1800 | -4600 | 10 | 0 | 60 | 3 |
| SC023 | ALM | left | 2019-05-31 | 1 | 3B 1.0 | 2500 | -1500 | -2900 | 15 | 135 | -135 | 0 |
| SC023 | ALM | right | 2019-05-31 | 2 | 3B 1.0 | 2500 | 1500 | -2900 | 15 | 45 | -45 | 1 |
| SC023 | Midbrain | left | 2019-05-31 | 3 | 3B 1.0 | -3800 | -1800 | -4650 | 10 | 180 | 120 | 2 |

| | | | | | | | | | | | | |
|---|---|---|---|---|---|---|---|---|---|---|---|---|
| SC023 | Midbrain | right | 2019-05-31 | 4 | 3B 1.0 | -3800 | 1800 | -4650 | 10 | 0 | 60 | 3 |
| SC023 | ALM | left | 2019-06-01 | 1 | 3B 1.0 | 2500 | -1500 | -2900 | 15 | 135 | -135 | 0 |
| SC023 | ALM | right | 2019-06-01 | 2 | 3B 1.0 | 2500 | 1500 | -2900 | 15 | 45 | -45 | 1 |
| SC023 | Midbrain | left | 2019-06-01 | 3 | 3B 1.0 | -3800 | -1800 | -4700 | 10 | 180 | 120 | 2 |
| SC023 | Midbrain | right | 2019-06-01 | 4 | 3B 1.0 | -3800 | 1800 | -4700 | 10 | 0 | 60 | 3 |
| SC023 | ALM | left | 2019-06-02 | 1 | 3B 1.0 | 2500 | -1500 | -2900 | 15 | 135 | -135 | 0 |
| SC023 | ALM | right | 2019-06-02 | 2 | 3B 1.0 | 2500 | 1500 | -2900 | 15 | 45 | -45 | 1 |
| SC023 | Midbrain | left | 2019-06-02 | 3 | 3B 1.0 | -3800 | -1800 | -4750 | 10 | 180 | 120 | 2 |
| SC023 | Midbrain | right | 2019-06-02 | 4 | 3B 1.0 | -3800 | 1800 | -4750 | 10 | 0 | 60 | 3 |
| SC023 | Striatum | left | 2019-06-03 | 1 | 3B 1.0 | 200 | -3000 | -4400 | 15 | 30 | -135 | 0 |
| SC023 | Striatum | right | 2019-06-03 | 2 | 3B 1.0 | 200 | 3000 | -4400 | 15 | 150 | -45 | 1 |
| SC023 | Thalamus | left | 2019-06-03 | 3 | 3B 1.0 | -2100 | -1500 | -4700 | 12 | 180 | 180 | 2 |
| SC023 | Thalamus | right | 2019-06-03 | 4 | 3B 1.0 | -2100 | 1500 | -4700 | 13.5 | 0 | 0 | 3 |
| SC023 | Striatum | left | 2019-06-04 | 1 | 3B 1.0 | 200 | -3000 | -4450 | 15 | 30 | -135 | 0 |
| SC023 | Striatum | right | 2019-06-04 | 2 | 3B 1.0 | 200 | 3000 | -4450 | 15 | 150 | -45 | 1 |
| SC023 | Thalamus | left | 2019-06-04 | 3 | 3B 1.0 | -2100 | -1500 | -4750 | 12 | 180 | 180 | 2 |
| SC023 | Thalamus | right | 2019-06-04 | 4 | 3B 1.0 | -2100 | 1500 | -4750 | 13.5 | 0 | 0 | 3 |
| SC026 | ALM | left | 2019-08-05 | 1 | 3B 1.0 | 2500 | -1500 | -2900 | 15 | 135 | -135 | 0 |
| SC026 | ALM | right | 2019-08-05 | 2 | 3B 1.0 | 2500 | 1500 | -2900 | 15 | 45 | -45 | 1 |
| SC026 | Thalamus | left | 2019-08-05 | 3 | 3B 1.0 | -1900 | -1100 | -4600 | 12.5 | 135 | 180 | 2 |
| SC026 | Midbrain | right | 2019-08-05 | 4 | 3B 1.0 | -3900 | 1750 | -4600 | 12.5 | 300 | 0 | 3 |
| SC026 | ALM | left | 2019-08-06 | 1 | 3B 1.0 | 2500 | -1500 | -2900 | 15 | 135 | -135 | 0 |
| SC026 | ALM | right | 2019-08-06 | 2 | 3B 1.0 | 2500 | 1500 | -2900 | 15 | 45 | -45 | 1 |
| SC026 | Thalamus | left | 2019-08-06 | 3 | 3B 1.0 | -1900 | -1100 | -4700 | 12.5 | 135 | 180 | 2 |

| | | | | | | | | | | | | |
|---|---|---|---|---|---|---|---|---|---|---|---|---|
| SC026 | Midbrain | right | 2019-08-06 | 4 | 3B 1.0 | -3900 | 1750 | -4700 | 12.5 | 300 | 0 | 3 |
| SC026 | ALM | left | 2019-08-07 | 1 | 3B 1.0 | 2500 | -1500 | -2900 | 15 | 135 | -135 | 0 |
| SC026 | ALM | right | 2019-08-07 | 2 | 3B 1.0 | 2500 | 1500 | -2900 | 15 | 45 | -45 | 1 |
| SC026 | Midbrain | right | 2019-08-07 | 3 | 3B 1.0 | -3900 | 1750 | -4750 | 12.5 | 300 | 0 | 2 |
| SC026 | ALM | left | 2019-08-08 | 1 | 3B 1.0 | 2500 | -1500 | -2900 | 15 | 135 | -135 | 0 |
| SC026 | ALM | right | 2019-08-08 | 2 | 3B 1.0 | 2500 | 1500 | -2900 | 15 | 45 | -45 | 1 |
| SC026 | Thalamus | left | 2019-08-08 | 3 | 3B 1.0 | -1900 | -1100 | -4800 | 12.5 | 135 | 180 | 2 |
| SC026 | Midbrain | right | 2019-08-08 | 4 | 3B 1.0 | -3900 | 1750 | -4800 | 12.5 | 300 | 0 | 3 |
| SC027 | ALM | left | 2019-07-29 | 1 | 3B 1.0 | 2500 | -1500 | -2900 | 15 | 135 | -135 | 0 |
| SC027 | Midbrain | left | 2019-07-29 | 2 | 3B 1.0 | -3900 | -1750 | -4600 | 12.5 | 180 | 120 | 1 |
| SC027 | Midbrain | right | 2019-07-29 | 3 | 3B 1.0 | -3900 | 1750 | -4600 | 12.5 | 0 | 60 | 2 |
| SC027 | ALM | left | 2019-07-30 | 1 | 3B 1.0 | 2500 | -1500 | -2900 | 15 | 135 | -135 | 0 |
| SC027 | ALM | right | 2019-07-30 | 2 | 3B 1.0 | 2500 | 1500 | -2900 | 15 | 45 | -45 | 1 |
| SC027 | Midbrain | left | 2019-07-30 | 3 | 3B 1.0 | -3900 | -1750 | -4650 | 12.5 | 180 | 120 | 2 |
| SC027 | Midbrain | right | 2019-07-30 | 4 | 3B 1.0 | -3900 | 1750 | -4650 | 12.5 | 0 | 60 | 3 |
| SC027 | ALM | left | 2019-07-31 | 1 | 3B 1.0 | 2500 | -1500 | -2900 | 15 | 135 | -135 | 0 |
| SC027 | ALM | right | 2019-07-31 | 2 | 3B 1.0 | 2500 | 1500 | -2900 | 15 | 45 | -45 | 1 |
| SC027 | Midbrain | left | 2019-07-31 | 3 | 3B 1.0 | -3900 | -1750 | -4650 | 12.5 | 180 | 120 | 2 |
| SC027 | Midbrain | right | 2019-07-31 | 4 | 3B 1.0 | -3900 | 1750 | -4650 | 12.5 | 0 | 60 | 3 |
| SC027 | ALM | left | 2019-08-01 | 1 | 3B 1.0 | 2500 | -1500 | -2900 | 15 | 135 | -135 | 0 |
| SC027 | Midbrain | left | 2019-08-01 | 2 | 3B 1.0 | -3900 | -1750 | -4750 | 12.5 | 180 | 120 | 1 |
| SC027 | Midbrain | right | 2019-08-01 | 3 | 3B 1.0 | -3900 | 1750 | -4750 | 12.5 | 0 | 60 | 2 |
| SC027 | Striatum | left | 2019-08-03 | 1 | 3B 1.0 | 0 | -3000 | -4400 | 15 | 30 | -135 | 0 |
| SC027 | Striatum | right | 2019-08-03 | 2 | 3B 1.0 | 0 | 3000 | -4400 | 15 | 150 | -45 | 1 |

| | | | | | | | | | | | | |
|---|---|---|---|---|---|---|---|---|---|---|---|---|
| SC027 | Thalamus | left | 2019-08-03 | 3 | 3B 1.0 | -2000 | -1000 | -4500 | 12.5 | 180 | 180 | 2 |
| SC027 | Thalamus | right | 2019-08-03 | 4 | 3B 1.0 | -2000 | 1000 | -4500 | 12.5 | 0 | 0 | 3 |
| SC027 | Striatum | left | 2019-08-04 | 1 | 3B 1.0 | 0 | -3000 | -4550 | 15 | 30 | -135 | 0 |
| SC027 | Striatum | right | 2019-08-04 | 2 | 3B 1.0 | 0 | 3000 | -4550 | 15 | 150 | -45 | 1 |
| SC027 | Thalamus | left | 2019-08-04 | 3 | 3B 1.0 | -2000 | -1000 | -4700 | 12.5 | 180 | 180 | 2 |
| SC027 | Thalamus | right | 2019-08-04 | 4 | 3B 1.0 | -2000 | 1000 | -4700 | 12.5 | 0 | 0 | 3 |
| SC030 | ALM | left | 2019-10-02 | 1 | 3B 1.0 | 2500 | -1500 | -2900 | 15 | 135 | -135 | 0 |
| SC030 | ALM | right | 2019-10-02 | 2 | 3B 1.0 | 2500 | 1500 | -2900 | 15 | 45 | -45 | 1 |
| SC030 | Medulla | left | 2019-10-02 | 3 | 3B 1.0 | -6650 | -2500 | -5300 | 20 | 180 | 120 | 2 |
| SC030 | Medulla | right | 2019-10-02 | 4 | 3B 1.0 | -6650 | 2500 | -5300 | 20 | 0 | 60 | 3 |
| SC030 | ALM | left | 2019-10-03 | 1 | 3B 1.0 | 2500 | -1500 | -2900 | 15 | 135 | -135 | 0 |
| SC030 | ALM | right | 2019-10-03 | 2 | 3B 1.0 | 2500 | 1500 | -2900 | 15 | 45 | -45 | 1 |
| SC030 | Medulla | left | 2019-10-03 | 3 | 3B 1.0 | -6650 | -2500 | -5000 | 20 | 180 | 120 | 2 |
| SC030 | Medulla | right | 2019-10-03 | 4 | 3B 1.0 | -6650 | 2500 | -5000 | 20 | 0 | 60 | 3 |
| SC030 | ALM | left | 2019-10-04 | 1 | 3B 1.0 | 2500 | -1500 | -2900 | 15 | 135 | -135 | 0 |
| SC030 | ALM | right | 2019-10-04 | 2 | 3B 1.0 | 2500 | 1500 | -2900 | 15 | 45 | -45 | 1 |
| SC030 | Medulla | left | 2019-10-04 | 3 | 3B 1.0 | -6650 | -2500 | -5000 | 20 | 180 | 120 | 2 |
| SC030 | Medulla | right | 2019-10-04 | 4 | 3B 1.0 | -6650 | 2500 | -5000 | 20 | 0 | 60 | 3 |
| SC030 | ALM | left | 2019-10-05 | 1 | 3B 1.0 | 2500 | -1500 | -2900 | 15 | 135 | -135 | 0 |
| SC030 | ALM | right | 2019-10-05 | 2 | 3B 1.0 | 2500 | 1500 | -2900 | 15 | 45 | -45 | 1 |
| SC030 | Medulla | left | 2019-10-05 | 3 | 3B 1.0 | -6650 | -2500 | -4900 | 15 | 180 | 120 | 2 |
| SC030 | Medulla | right | 2019-10-05 | 4 | 3B 1.0 | -6650 | 2500 | -4900 | 15 | 0 | 60 | 3 |
| SC031 | ALM | left | 2019-10-21 | 1 | 3B 1.0 | 2500 | -1500 | -2900 | 15 | 135 | -135 | 0 |
| SC031 | ALM | right | 2019-10-21 | 2 | 3B 1.0 | 2500 | 1500 | -2900 | 15 | 45 | -45 | 1 |

| | | | | | | | | | | | | |
|---|---|---|---|---|---|---|---|---|---|---|---|---|
| SC031 | Medulla | left | 2019-10-21 | 3 | 3B 1.0 | -6700 | -2000 | -4750 | 10 | 180 | 120 | 2 |
| SC031 | Medulla | right | 2019-10-21 | 4 | 3B 1.0 | -6700 | 2000 | -4750 | 10 | 0 | 60 | 3 |
| SC031 | ALM | left | 2019-10-22 | 1 | 3B 2.0 MS | 2500 | -1500 | -1200 | 15 | 135 | -180 | 0 |
| SC031 | Medulla | left | 2019-10-22 | 2 | 3B 2.0 MS | -6700 | -2000 | -4700 | 10 | 180 | -160 | 1 |
| SC031 | Medulla | right | 2019-10-22 | 3 | 3B 2.0 MS | -6700 | 2000 | -4700 | 10 | 0 | -20 | 2 |
| SC031 | ALM | right | 2019-10-23 | 1 | 3B 2.0 MS | 2500 | 1500 | -1200 | 15 | 45 | 0 | 0 |
| SC031 | Medulla | left | 2019-10-23 | 2 | 3B 2.0 MS | -6700 | -2000 | -4700 | 10 | 180 | -160 | 1 |
| SC031 | Medulla | right | 2019-10-23 | 3 | 3B 2.0 MS | -6700 | 2000 | -4700 | 10 | 0 | -20 | 2 |
| SC032 | ALM | left | 2019-12-17 | 1 | 3B 2.0 MS | 2500 | -1500 | -1150 | 12 | 135 | -135 | 0 |
| SC032 | Midbrain | left | 2019-12-17 | 2 | 3B 2.0 SS | -3800 | -1800 | -4750 | 8 | 240 | 180 | 1 |
| SC032 | Medulla | right | 2019-12-17 | 3 | 3B 2.0 MS | -6500 | 1200 | -4750 | 10 | 0 | 0 | 2 |
| SC032 | ALM | left | 2019-12-18 | 1 | 3B 2.0 MS | 2500 | -1500 | -1200 | 12 | 135 | -135 | 0 |
| SC032 | Midbrain | left | 2019-12-18 | 2 | 3B 2.0 SS | -3800 | -1800 | -4750 | 6 | 240 | 180 | 1 |
| SC032 | Medulla | right | 2019-12-18 | 3 | 3B 2.0 MS | -6500 | 1200 | -4700 | 10 | 0 | 0 | 2 |
| SC032 | ALM | right | 2019-12-19 | 1 | 3B 2.0 MS | 2500 | 1500 | -1100 | 12 | 45 | -45 | 0 |
| SC032 | Medulla | left | 2019-12-19 | 2 | 3B 2.0 MS | -6500 | -1200 | -4800 | 12 | 180 | 160 | 1 |
| SC032 | Midbrain | right | 2019-12-19 | 3 | 3B 2.0 SS | -3800 | 1800 | -4750 | 8 | 270 | 90 | 2 |
| SC032 | ALM | right | 2019-12-20 | 1 | 3B 2.0 MS | 2500 | 1500 | -1200 | 12 | 45 | -45 | 0 |
| SC032 | Medulla | left | 2019-12-20 | 2 | 3B 2.0 MS | -6500 | -1200 | -4600 | 15 | 180 | 170 | 1 |
| SC032 | Midbrain | right | 2019-12-20 | 3 | 3B 2.0 SS | -3800 | 1800 | -4850 | 8 | 270 | 90 | 2 |
| SC033 | ALM | left | 2019-11-12 | 1 | 3B 2.0 MS | 2500 | -1500 | -1200 | 15 | 135 | -180 | 0 |
| SC033 | Midbrain | left | 2019-11-12 | 2 | 3B 2.0 SS | -3800 | -1800 | -4800 | 8 | 225 | 90 | 1 |
| SC033 | Medulla | right | 2019-11-12 | 3 | 3B 2.0 MS | -6500 | 1200 | -4800 | 8 | 0 | 90 | 2 |
| SC033 | ALM | left | 2019-11-13 | 1 | 3B 2.0 MS | 2500 | -1500 | -1200 | 15 | 135 | -180 | 0 |

| | | | | | | | | | | | | |
|---|---|---|---|---|---|---|---|---|---|---|---|---|
| SC033 | Midbrain | left | 2019-11-13 | 2 | 3B 2.0 SS | -3800 | -1800 | -4750 | 8 | 225 | 90 | 1 |
| SC033 | Medulla | right | 2019-11-13 | 3 | 3B 2.0 MS | -6500 | 1200 | -4850 | 8 | 0 | 90 | 2 |
| SC033 | ALM | right | 2019-11-14 | 1 | 3B 2.0 MS | 2500 | 1500 | -1200 | 15 | 45 | -45 | 0 |
| SC033 | Medulla | left | 2019-11-14 | 2 | 3B 2.0 MS | -6500 | -1200 | -4800 | 8 | 180 | 90 | 1 |
| SC033 | Midbrain | right | 2019-11-14 | 3 | 3B 2.0 SS | -3800 | 1800 | -4800 | 8 | 315 | 90 | 2 |
| SC033 | ALM | right | 2019-11-15 | 1 | 3B 2.0 MS | 2500 | 1500 | -1200 | 12 | 45 | -45 | 0 |
| SC033 | Medulla | left | 2019-11-15 | 2 | 3B 2.0 MS | -6500 | -1200 | -4900 | 8 | 180 | 90 | 1 |
| SC033 | Midbrain | right | 2019-11-15 | 3 | 3B 2.0 SS | -3800 | 1800 | -4900 | 8 | 315 | 90 | 2 |
| SC035 | ALM | left | 2020-01-07 | 1 | 3B 2.0 MS | 2500 | -1500 | -1000 | 12 | 135 | -135 | 0 |
| SC035 | Midbrain | left | 2020-01-07 | 2 | 3B 2.0 SS | -3700 | -1800 | -4800 | 12 | 240 | 180 | 1 |
| SC035 | Medulla | right | 2020-01-07 | 3 | 3B 2.0 MS | -6600 | 2000 | -4800 | 12 | 0 | 0 | 2 |
| SC035 | ALM | left | 2020-01-08 | 1 | 3B 2.0 MS | 2500 | -1500 | -1100 | 12 | 135 | -135 | 0 |
| SC035 | Midbrain | left | 2020-01-08 | 2 | 3B 2.0 SS | -3700 | -1800 | -4750 | 14 | 240 | 180 | 1 |
| SC035 | Medulla | right | 2020-01-08 | 3 | 3B 2.0 MS | -6600 | 2000 | -4750 | 12 | 0 | 0 | 2 |
| SC035 | ALM | right | 2020-01-09 | 1 | 3B 2.0 MS | 2500 | 1500 | -1000 | 12 | 45 | -45 | 0 |
| SC035 | Medulla | left | 2020-01-09 | 2 | 3B 2.0 MS | -6600 | -1800 | -4750 | 12 | 180 | 160 | 1 |
| SC035 | Midbrain | right | 2020-01-09 | 3 | 3B 2.0 SS | -3700 | 1750 | -4850 | 12 | 300 | 0 | 2 |
| SC035 | ALM | right | 2020-01-10 | 1 | 3B 2.0 MS | 2500 | 1500 | -1200 | 12 | 45 | -45 | 0 |
| SC035 | Medulla | left | 2020-01-10 | 2 | 3B 2.0 MS | -6600 | -1800 | -4600 | 12 | 180 | 160 | 1 |
| SC035 | Midbrain | right | 2020-01-10 | 3 | 3B 2.0 SS | -3700 | 1750 | -4800 | 12 | 300 | 0 | 2 |
| SC038 | ALM | left | 2019-11-19 | 1 | 3B 2.0 MS | 2500 | -1500 | -1200 | 12 | 135 | -135 | 0 |
| SC038 | Midbrain | left | 2019-11-19 | 2 | 3B 2.0 MS | -3700 | -1500 | -2000 | 8 | 180 | 90 | 1 |
| SC038 | Midbrain | right | 2019-11-19 | 3 | 3B 2.0 MS | -3700 | 1500 | -2250 | 8 | 0 | 90 | 2 |
| SC038 | ALM | left | 2019-11-19 | 1 | 3B 2.0 MS | 2500 | -1500 | -1200 | 12 | 135 | -135 | 0 |

| | | | | | | | | | | | | |
|---|---|---|---|---|---|---|---|---|---|---|---|---|
| SC038 | Midbrain | left | 2019-11-19 | 2 | 3B 2.0 MS | -3700 | -1500 | -3000 | 8 | 180 | 90 | 1 |
| SC038 | Midbrain | right | 2019-11-19 | 3 | 3B 2.0 MS | -3700 | 1500 | -3000 | 8 | 0 | 90 | 2 |
| SC038 | ALM | left | 2019-11-20 | 1 | 3B 2.0 MS | 2500 | -1500 | -1000 | 12 | 135 | -135 | 0 |
| SC038 | Midbrain | left | 2019-11-20 | 2 | 3B 2.0 MS | -3700 | -1500 | -2800 | 8 | 180 | 90 | 1 |
| SC038 | Midbrain | right | 2019-11-20 | 3 | 3B 2.0 MS | -3700 | 1500 | -2800 | 8 | 0 | 90 | 2 |
| SC038 | ALM | left | 2019-11-20 | 1 | 3B 2.0 MS | 2500 | -1500 | -1000 | 12 | 135 | -135 | 0 |
| SC038 | Midbrain | left | 2019-11-20 | 2 | 3B 2.0 MS | -3700 | -1500 | -3600 | 8 | 180 | 90 | 1 |
| SC038 | Midbrain | right | 2019-11-20 | 3 | 3B 2.0 MS | -3700 | 1500 | -3600 | 8 | 0 | 90 | 2 |
| SC038 | ALM | right | 2019-11-21 | 1 | 3B 2.0 MS | 2500 | 1500 | -1100 | 12 | 45 | -45 | 0 |
| SC038 | Thalamus | left | 2019-11-21 | 2 | 3B 2.0 MS | -1750 | -1300 | -4000 | 8 | 180 | 90 | 1 |
| SC038 | Thalamus | right | 2019-11-21 | 3 | 3B 2.0 MS | -1750 | 1300 | -4000 | 8 | 0 | 90 | 2 |
| SC038 | ALM | right | 2019-11-21 | 1 | 3B 2.0 MS | 2500 | 1500 | -1100 | 12 | 45 | -45 | 0 |
| SC038 | Thalamus | left | 2019-11-21 | 2 | 3B 2.0 MS | -1750 | -1300 | -3280 | 8 | 180 | 90 | 1 |
| SC038 | Thalamus | right | 2019-11-21 | 3 | 3B 2.0 MS | -1750 | 1300 | -3280 | 8 | 0 | 90 | 2 |
| SC038 | ALM | right | 2019-11-22 | 1 | 3B 2.0 MS | 2500 | 1500 | -1100 | 12 | 45 | -45 | 0 |
| SC038 | Thalamus | left | 2019-11-22 | 2 | 3B 2.0 MS | -1750 | -1300 | -4500 | 8 | 180 | 90 | 1 |
| SC038 | Thalamus | right | 2019-11-22 | 3 | 3B 2.0 MS | -1750 | 1300 | -4500 | 8 | 0 | 90 | 2 |
| SC038 | ALM | right | 2019-11-22 | 1 | 3B 2.0 MS | 2500 | 1500 | -1100 | 12 | 45 | -45 | 0 |
| SC038 | Thalamus | left | 2019-11-22 | 2 | 3B 2.0 MS | -1750 | -1300 | -3780 | 8 | 180 | 90 | 1 |
| SC038 | Thalamus | right | 2019-11-22 | 3 | 3B 2.0 MS | -1750 | 1300 | -3780 | 8 | 0 | 90 | 2 |
| SC038 | ALM | left | 2019-11-23 | 1 | 3B 2.0 SS | 2500 | -1500 | -2750 | 12 | 135 | -135 | 0 |
| SC038 | Striatum | left | 2019-11-23 | 2 | 3B 2.0 MS | 0 | -2500 | -3200 | 8 | 180 | 90 | 1 |
| SC038 | Striatum | right | 2019-11-23 | 3 | 3B 2.0 MS | 0 | 2500 | -3200 | 8 | 0 | 90 | 2 |
| SC038 | ALM | left | 2019-11-23 | 1 | 3B 2.0 SS | 2500 | -1500 | -2750 | 12 | 135 | -135 | 0 |

| | | | | | | | | | | | | |
|---|---|---|---|---|---|---|---|---|---|---|---|---|
| SC038 | Striatum | left | 2019-11-23 | 2 | 3B 2.0 MS | 0 | -2500 | -2480 | 8 | 180 | 90 | 1 |
| SC038 | Striatum | right | 2019-11-23 | 3 | 3B 2.0 MS | 0 | 2500 | -2480 | 8 | 0 | 90 | 2 |
| SC043 | ALM | left | 2020-09-21 | 1 | 3B 1.0 | 2500 | -1500 | -3000 | 15 | 135 | -135 | 0 |
| SC043 | ALM | right | 2020-09-21 | 2 | 3B 1.0 | 2500 | 1500 | -3000 | 15 | 45 | -45 | 1 |
| SC043 | Medulla | left | 2020-09-21 | 3 | 3B 1.0 | -6900 | -2400 | -4800 | 12 | 180 | 90 | 2 |
| SC043 | Medulla | right | 2020-09-21 | 4 | 3B 1.0 | -6800 | 2400 | -4800 | 12 | 0 | 90 | 3 |
| SC043 | Medulla | left | 2020-09-21 | 5 | 3B 1.0 | -6800 | -1000 | -4800 | 12 | 180 | 90 | 4 |
| SC043 | ALM | right | 2020-09-22 | 1 | 3B 1.0 | 2500 | 1500 | -3000 | 15 | 45 | -45 | 0 |
| SC043 | Medulla | left | 2020-09-22 | 2 | 3B 1.0 | -6800 | -2400 | -4900 | 12 | 180 | 90 | 1 |
| SC043 | Medulla | right | 2020-09-22 | 3 | 3B 1.0 | -6800 | 2200 | -4900 | 12 | 0 | 90 | 2 |
| SC043 | Medulla | left | 2020-09-22 | 4 | 3B 1.0 | -6900 | -1000 | -4900 | 12 | 180 | 90 | 3 |
| SC043 | ALM | left | 2020-09-23 | 1 | 3B 1.0 | 2500 | -1500 | -3000 | 15 | 135 | -135 | 0 |
| SC043 | ALM | right | 2020-09-23 | 2 | 3B 1.0 | 2500 | 1500 | -3000 | 15 | 45 | -45 | 1 |
| SC043 | Medulla | left | 2020-09-23 | 3 | 3B 1.0 | -7000 | -2400 | -4700 | 12 | 180 | 90 | 2 |
| SC043 | Medulla | right | 2020-09-23 | 4 | 3B 1.0 | -7000 | 2400 | -4700 | 12 | 0 | 90 | 3 |
| SC043 | Medulla | left | 2020-09-23 | 5 | 3B 1.0 | -7000 | -1000 | -4700 | 12 | 180 | 90 | 4 |
| SC043 | ALM | left | 2020-09-24 | 1 | 3B 1.0 | 2500 | -1500 | -3000 | 15 | 135 | -135 | 0 |
| SC043 | ALM | right | 2020-09-24 | 2 | 3B 1.0 | 2500 | 1500 | -3000 | 15 | 45 | -45 | 1 |
| SC043 | Medulla | left | 2020-09-24 | 3 | 3B 1.0 | -7000 | -2400 | -4800 | 12 | 180 | 90 | 2 |
| SC043 | Medulla | right | 2020-09-24 | 4 | 3B 1.0 | -7000 | 2400 | -4800 | 12 | 0 | 90 | 3 |
| SC043 | Medulla | right | 2020-09-24 | 5 | 3B 1.0 | -7000 | 1000 | -4800 | 12 | 0 | 90 | 4 |
| SC043 | ALM | left | 2020-09-25 | 1 | 3B 1.0 | 2500 | -1500 | -3000 | 15 | 135 | -135 | 0 |
| SC043 | ALM | right | 2020-09-25 | 2 | 3B 1.0 | 2500 | 1500 | -3000 | 15 | 45 | -45 | 1 |
| SC043 | Midbrain | left | 2020-09-25 | 3 | 3B 1.0 | -4000 | -1800 | -4700 | 12 | 270 | 90 | 2 |

| | | | | | | | | | | | | |
|---|---|---|---|---|---|---|---|---|---|---|---|---|
| SC043 | Midbrain | right | 2020-09-25 | 4 | 3B 1.0 | -4000 | 1800 | -4700 | 12 | 270 | 90 | 3 |
| SC043 | ALM | right | 2020-09-28 | 1 | 3B 1.0 | 2500 | 1500 | -3000 | 15 | 45 | -45 | 0 |
| SC043 | Midbrain | left | 2020-09-28 | 2 | 3B 1.0 | -4100 | -1800 | -4800 | 12 | 270 | 90 | 1 |
| SC043 | Midbrain | right | 2020-09-28 | 3 | 3B 1.0 | -4100 | 1800 | -4800 | 12 | 270 | 90 | 2 |
| SC045 | ALM | left | 2020-12-09 | 1 | 3B 2.0 MS | 2500 | -1500 | -1200 | 12 | 120 | -150 | 0 |
| SC045 | Striatum | left | 2020-12-09 | 3 | 3B 2.0 MS | 0 | -3300 | -4000 | 0 | 180 | 100 | 2 |
| SC045 | Striatum | right | 2020-12-09 | 4 | 3B 2.0 MS | 0 | 3300 | -4000 | 0 | 0 | 80 | 3 |
| SC045 | ALM | left | 2020-12-10 | 1 | 3B 2.0 MS | 2500 | -1500 | -1200 | 12 | 120 | -150 | 0 |
| SC045 | ALM | right | 2020-12-10 | 2 | 3B 2.0 MS | 2500 | 1500 | -1200 | 12 | 60 | -20 | 1 |
| SC045 | Striatum | left | 2020-12-10 | 3 | 3B 2.0 MS | -500 | -3300 | -4000 | 0 | 180 | 100 | 2 |
| SC045 | Striatum | right | 2020-12-10 | 4 | 3B 2.0 MS | -500 | 3300 | -4000 | 0 | 0 | 80 | 3 |
| SC045 | ALM | left | 2020-12-11 | 1 | 3B 1.0 | 2500 | -1500 | -3000 | 12 | 135 | -135 | 0 |
| SC045 | ALM | right | 2020-12-11 | 2 | 3B 1.0 | 2500 | 1500 | -3000 | 12 | 45 | -45 | 1 |
| SC045 | Striatum | left | 2020-12-11 | 3 | 3B 1.0 | -500 | -3200 | -4200 | 20 | 270 | 90 | 2 |
| SC045 | Striatum | right | 2020-12-11 | 4 | 3B 1.0 | 0 | 3200 | -4200 | 20 | 270 | 90 | 3 |
| SC045 | Striatum | right | 2020-12-11 | 5 | 3B 1.0 | -500 | 3200 | -4200 | 20 | 270 | 90 | 4 |
| SC045 | Cortex | left | 2020-12-14 | 1 | 3B 1.0 | -1800 | -4400 | -3700 | 6 | 120 | -120 | 0 |
| SC045 | Cortex | right | 2020-12-14 | 2 | 3B 1.0 | -1800 | 4400 | -3700 | 6 | 30 | -30 | 1 |
| SC045 | Midbrain | left | 2020-12-14 | 3 | 3B 1.0 | -4000 | -1800 | -4800 | 6 | 270 | 180 | 2 |
| SC045 | Midbrain | right | 2020-12-14 | 4 | 3B 1.0 | -4000 | 1800 | -4800 | 6 | 280 | 0 | 3 |
| SC045 | BLA | left | 2020-12-16 | 1 | 3B 1.0 | -1800 | -4400 | -4800 | 12 | 180 | 180 | 0 |
| SC045 | BLA | right | 2020-12-16 | 2 | 3B 1.0 | -1800 | 4400 | -4800 | 12 | 0 | 0 | 1 |
| SC045 | Midbrain | right | 2020-12-16 | 3 | 3B 1.0 | -4000 | 1800 | -4900 | 8 | 260 | 90 | 2 |
| SC045 | Thalamus | left | 2020-12-17 | 1 | 3B 2.0 MS | -1950 | -1700 | -4700 | 5 | 180 | -100 | 0 |

| | | | | | | | | | | | | |
|---|---|---|---|---|---|---|---|---|---|---|---|---|
| SC045 | Thalamus | right | 2020-12-17 | 2 | 3B 2.0 MS | -2000 | 1800 | -4800 | 5 | 0 | 10 | 1 |
| SC045 | Midbrain | left | 2020-12-17 | 3 | 3B 1.0 | -4000 | -1800 | -4900 | 8 | 280 | 90 | 2 |
| SC045 | Thalamus | left | 2020-12-18 | 1 | 3B 2.0 MS | -2050 | -1700 | -4200 | 5 | 180 | -90 | 0 |
| SC045 | Thalamus | right | 2020-12-18 | 2 | 3B 2.0 MS | -2000 | 1600 | -4700 | 5 | 0 | 0 | 1 |
| SC045 | Midbrain | left | 2020-12-18 | 3 | 3B 1.0 | -4000 | -1800 | -4800 | 8 | 280 | 90 | 2 |
| SC048 | ALM | left | 2020-12-24 | 1 | 3B 2.0 MS | 2500 | -1800 | -1100 | 10 | 135 | -135 | 0 |
| SC048 | Midbrain | right | 2020-12-24 | 2 | 3B 2.0 MS | -3100 | 1500 | -4800 | 1 | 0 | 90 | 1 |
| SC048 | Thalamus | left | 2020-12-24 | 3 | 3B 2.0 MS | -1900 | -1500 | -4600 | 5 | 180 | -90 | 2 |
| SC048 | ALM | left | 2020-12-25 | 1 | 3B 2.0 MS | 2500 | -1500 | -1050 | 10 | 135 | -135 | 0 |
| SC048 | Midbrain | right | 2020-12-25 | 2 | 3B 2.0 MS | -2800 | 1500 | -5200 | 6 | 0 | 90 | 1 |
| SC048 | Thalamus | left | 2020-12-25 | 3 | 3B 2.0 MS | -2000 | -1400 | -4400 | 10 | 180 | 90 | 2 |
| SC048 | ALM | left | 2020-12-26 | 1 | 3B 1.0 | 2500 | -1600 | -3200 | 10 | 135 | -135 | 0 |
| SC048 | Midbrain | right | 2020-12-26 | 2 | 3B 1.0 | -3000 | 1900 | -5250 | 2.5 | 0 | 30 | 1 |
| SC048 | Thalamus | left | 2020-12-26 | 3 | 3B 1.0 | -2000 | -1900 | -4850 | 6 | 180 | 120 | 2 |
| SC048 | ALM | left | 2020-12-29 | 1 | 3B 1.0 | 2500 | -1500 | -3100 | 10 | 135 | -135 | 0 |
| SC048 | ALM | right | 2020-12-29 | 2 | 3B 1.0 | 2500 | 1500 | -3100 | 10 | 45 | -45 | 1 |
| SC048 | Midbrain | left | 2020-12-29 | 3 | 3B 1.0 | -3000 | -1900 | -5000 | 5 | 180 | 120 | 2 |
| SC048 | Thalamus | right | 2020-12-29 | 4 | 3B 1.0 | -2000 | 1900 | -4600 | 8 | 0 | 30 | 3 |
| SC048 | ALM | left | 2020-12-30 | 1 | 3B 1.0 | 2500 | -2100 | -3000 | 13 | 135 | -135 | 0 |
| SC048 | ALM | right | 2020-12-30 | 2 | 3B 1.0 | 2500 | 2000 | -3000 | 13 | 45 | -45 | 1 |
| SC048 | Midbrain | left | 2020-12-30 | 3 | 3B 1.0 | -2950 | -1900 | -5000 | 5 | 225 | 120 | 2 |
| SC048 | Thalamus | right | 2020-12-30 | 4 | 3B 1.0 | -2100 | 1700 | -4900 | 8 | 315 | 30 | 3 |
| SC048 | Thalamus | left | 2020-12-30 | 5 | 3B 1.0 | -2000 | -1500 | -5000 | 8 | 225 | 120 | 4 |
| SC048 | Striatum | left | 2020-12-31 | 1 | 3B 1.0 | -300 | -3600 | -4400 | 8 | 135 | -135 | 0 |

| | | | | | | | | | | | | |
|---|---|---|---|---|---|---|---|---|---|---|---|---|
| SC048 | Striatum | right | 2020-12-31 | 2 | 3B 1.0 | -300 | 3600 | -4400 | 13 | 45 | -45 | 1 |
| SC048 | Midbrain | left | 2020-12-31 | 3 | 3B 1.0 | -3000 | -1200 | -5500 | 8 | 225 | 0 | 2 |
| SC048 | Striatum | left | 2021-01-01 | 1 | 3B 1.0 | -500 | -3500 | -4400 | 10 | 135 | -135 | 0 |
| SC048 | Striatum | right | 2021-01-01 | 2 | 3B 1.0 | -500 | 3600 | -4400 | 10 | 45 | -45 | 1 |
| SC048 | Midbrain | left | 2021-01-01 | 3 | 3B 1.0 | -3000 | -1500 | -5000 | 5 | 225 | 0 | 2 |
| SC048 | Cortex | left | 2021-01-03 | 1 | 3B 2.0 MS | -2500 | -4500 | -3600 | 6 | 135 | 180 | 0 |
| SC048 | Cortex | right | 2021-01-03 | 2 | 3B 2.0 MS | -2500 | 4500 | -3600 | 6 | 45 | 0 | 1 |
| SC048 | Midbrain | left | 2021-01-03 | 3 | 3B 1.0 | -4200 | -1900 | -4700 | 6 | 260 | 180 | 2 |
| SC048 | Midbrain | right | 2021-01-03 | 4 | 3B 1.0 | -4200 | 1800 | -4700 | 6 | 280 | 0 | 3 |
| SC048 | BLA | left | 2021-01-04 | 1 | 3B 2.0 MS | -2500 | -4500 | -4500 | 15 | 135 | 180 | 0 |
| SC048 | BLA | right | 2021-01-04 | 2 | 3B 2.0 MS | -2500 | 4500 | -4500 | 15 | 45 | 0 | 1 |
| SC048 | Midbrain | left | 2021-01-04 | 3 | 3B 1.0 | -4200 | -1800 | -5000 | 15 | 260 | 180 | 2 |
| SC048 | Midbrain | right | 2021-01-04 | 4 | 3B 1.0 | -4200 | 1900 | -5000 | 15 | 280 | 0 | 3 |
| SC049 | Cortex | left | 2021-01-06 | 1 | 3B 1.0 | -1000 | -4200 | -4200 | 6 | 225 | 120 | 0 |
| SC049 | Cortex | right | 2021-01-06 | 2 | 3B 1.0 | 1400 | 4400 | -4000 | 6 | 45 | -45 | 1 |
| SC049 | Midbrain | right | 2021-01-06 | 3 | 3B 1.0 | -4100 | 1800 | -5100 | 12 | 270 | 0 | 2 |
| SC049 | Striatum | left | 2021-01-06 | 4 | 3B 1.0 | -200 | -3500 | -4200 | 6 | 225 | 120 | 3 |
| SC049 | Cortex | left | 2021-01-07 | 1 | 3B 1.0 | -1000 | -4250 | -4300 | 6 | 225 | 120 | 0 |
| SC049 | Cortex | right | 2021-01-07 | 2 | 3B 1.0 | -1200 | 4500 | -4300 | 6 | 45 | -45 | 1 |
| SC049 | Midbrain | right | 2021-01-07 | 3 | 3B 1.0 | -4150 | 1700 | -5000 | 12 | 270 | 0 | 2 |
| SC049 | Striatum | left | 2021-01-07 | 4 | 3B 1.0 | -200 | -3550 | -4300 | 6 | 225 | 120 | 3 |
| SC049 | Cortex | left | 2021-01-08 | 1 | 3B 1.0 | -1000 | -4300 | -4300 | 6 | 225 | 120 | 0 |
| SC049 | Striatum | right | 2021-01-08 | 2 | 3B 1.0 | -300 | 3500 | -4450 | 6 | 45 | -45 | 1 |
| SC049 | Midbrain | right | 2021-01-08 | 3 | 3B 1.0 | -4100 | 1600 | -5100 | 10 | 260 | 0 | 2 |

| | | | | | | | | | | | | |
|---|---|---|---|---|---|---|---|---|---|---|---|---|
| SC049 | Striatum | left | 2021-01-08 | 4 | 3B 1.0 | -200 | -3600 | -4300 | 6 | 225 | 120 | 3 |
| SC049 | ALM | left | 2021-01-10 | 1 | 3B 2.0 MS | 2500 | -1500 | -1200 | 12 | 135 | -135 | 0 |
| SC049 | ALM | right | 2021-01-10 | 2 | 3B 2.0 MS | 2500 | 1500 | -1200 | 12 | 45 | -45 | 1 |
| SC049 | BLA | left | 2021-01-10 | 3 | 3B 2.0 MS | -500 | -4100 | -4800 | 8 | 180 | 180 | 2 |
| SC049 | BLA | right | 2021-01-10 | 4 | 3B 2.0 MS | -500 | 4100 | -4800 | 8 | 0 | 0 | 3 |
| SC049 | ALM | left | 2021-01-11 | 1 | 3B 2.0 MS | 2500 | -1500 | -1200 | 12 | 135 | -135 | 0 |
| SC049 | ALM | right | 2021-01-11 | 2 | 3B 2.0 MS | 2500 | 1500 | -1200 | 12 | 45 | -45 | 1 |
| SC049 | BLA | left | 2021-01-11 | 3 | 3B 2.0 MS | -500 | -3900 | -4800 | 8 | 180 | 180 | 2 |
| SC049 | BLA | right | 2021-01-11 | 4 | 3B 2.0 MS | -500 | 3900 | -4800 | 8 | 0 | 0 | 3 |
| SC049 | ALM | right | 2021-01-12 | 1 | 3B 1.0 | 2500 | 1500 | -3300 | 12 | 45 | -45 | 0 |
| SC049 | Thalamus | left | 2021-01-12 | 2 | 3B 1.0 | -2000 | -1900 | -5200 | 8 | 180 | 180 | 1 |
| SC049 | Thalamus | right | 2021-01-12 | 3 | 3B 1.0 | -1100 | 1900 | -5200 | 8 | 0 | 0 | 2 |
| SC049 | Thalamus | left | 2021-01-12 | 4 | 3B 1.0 | -2000 | -600 | -5200 | 8 | 180 | 180 | 3 |
| SC049 | ALM | left | 2021-01-13 | 1 | 3B 1.0 | 2500 | -1500 | -3300 | 14 | 135 | -135 | 0 |
| SC049 | ALM | right | 2021-01-13 | 2 | 3B 1.0 | 2500 | 1500 | -3300 | 14 | 45 | -45 | 1 |
| SC049 | Thalamus | left | 2021-01-13 | 3 | 3B 1.0 | -1800 | -1900 | -5000 | 8 | 180 | 180 | 2 |
| SC049 | Thalamus | right | 2021-01-13 | 4 | 3B 1.0 | -1300 | 1900 | -5000 | 8 | 0 | 0 | 3 |
| SC049 | Thalamus | left | 2021-01-13 | 5 | 3B 1.0 | -2000 | -800 | -5000 | 8 | 180 | 180 | 4 |
| SC052 | Striatum | left | 2021-01-21 | 1 | 3B 1.0 | 0 | -3500 | -4600 | 12 | 135 | -135 | 0 |
| SC052 | Striatum | right | 2021-01-21 | 2 | 3B 1.0 | 0 | 3400 | -4800 | 12 | 45 | -45 | 1 |
| SC052 | Midbrain | left | 2021-01-21 | 3 | 3B 1.0 | -3900 | -1800 | -5100 | 8 | 260 | 180 | 2 |
| SC052 | Midbrain | right | 2021-01-21 | 4 | 3B 1.0 | -3900 | 1800 | -5100 | 8 | 280 | 0 | 3 |
| SC052 | Striatum | left | 2021-01-22 | 1 | 3B 1.0 | 0 | -3400 | -4700 | 12 | 135 | -135 | 0 |
| SC052 | Striatum | right | 2021-01-22 | 2 | 3B 1.0 | 0 | 3500 | -4700 | 12 | 45 | -45 | 1 |

| | | | | | | | | | | | | |
|---|---|---|---|---|---|---|---|---|---|---|---|---|
| SC052 | Midbrain | left | 2021-01-22 | 3 | 3B 1.0 | -3900 | -1700 | -5150 | 8 | 260 | 180 | 2 |
| SC052 | Midbrain | right | 2021-01-22 | 4 | 3B 1.0 | -3900 | 1700 | -5150 | 8 | 280 | 0 | 3 |
| SC052 | Striatum | left | 2021-01-23 | 1 | 3B 1.0 | -300 | -3450 | -4750 | 12 | 135 | -135 | 0 |
| SC052 | Striatum | right | 2021-01-23 | 2 | 3B 1.0 | -300 | 3450 | -4750 | 12 | 45 | -45 | 1 |
| SC052 | Midbrain | left | 2021-01-23 | 3 | 3B 1.0 | -4000 | -1750 | -5200 | 8 | 260 | 180 | 2 |
| SC052 | Midbrain | right | 2021-01-23 | 4 | 3B 1.0 | -4000 | 1750 | -5200 | 8 | 280 | 0 | 3 |
| SC052 | ALM | left | 2021-01-25 | 1 | 3B 1.0 | 2500 | -1500 | -3400 | 12 | 135 | -135 | 0 |
| SC052 | ALM | right | 2021-01-25 | 2 | 3B 1.0 | 2500 | 1500 | -3400 | 12 | 45 | -45 | 1 |
| SC052 | Thalamus | left | 2021-01-25 | 3 | 3B 1.0 | -2000 | -1800 | -4800 | 10 | 220 | 160 | 2 |
| SC052 | Thalamus | right | 2021-01-25 | 4 | 3B 1.0 | -1900 | 1800 | -5000 | 12 | 340 | 10 | 3 |
| SC052 | Thalamus | left | 2021-01-25 | 5 | 3B 1.0 | -1200 | -800 | -4800 | 10 | 220 | 160 | 4 |
| SC052 | ALM | left | 2021-01-26 | 1 | 3B 1.0 | 2500 | -1600 | -3500 | 12 | 135 | -135 | 0 |
| SC052 | Thalamus | left | 2021-01-26 | 2 | 3B 1.0 | -1800 | -1800 | -5000 | 10 | 220 | 160 | 1 |
| SC052 | Thalamus | right | 2021-01-26 | 3 | 3B 1.0 | -2000 | 1700 | -5200 | 10 | 340 | 10 | 2 |
| SC052 | Thalamus | right | 2021-01-26 | 4 | 3B 1.0 | -1400 | 800 | -5000 | 10 | 340 | 10 | 3 |
| SC052 | ALM | left | 2021-01-27 | 1 | 3B 1.0 | 2500 | -1700 | -3500 | 12 | 135 | -135 | 0 |
| SC052 | ALM | right | 2021-01-27 | 2 | 3B 1.0 | 2500 | 1400 | -3500 | 12 | 45 | -45 | 1 |
| SC052 | Thalamus | left | 2021-01-27 | 3 | 3B 1.0 | -1850 | -1750 | -4900 | 10 | 220 | 160 | 2 |
| SC052 | Thalamus | right | 2021-01-27 | 4 | 3B 1.0 | -1900 | 1700 | -4900 | 10 | 340 | 10 | 3 |
| SC052 | Thalamus | right | 2021-01-27 | 5 | 3B 1.0 | -1300 | 800 | -4900 | 10 | 340 | 10 | 4 |
| SC052 | ALM | left | 2021-01-28 | 1 | 3B 1.0 | 2400 | -1500 | -3500 | 12 | 135 | -135 | 0 |
| SC052 | ALM | right | 2021-01-28 | 2 | 3B 1.0 | 2400 | 1400 | -3500 | 12 | 45 | -45 | 1 |
| SC052 | Thalamus | left | 2021-01-28 | 3 | 3B 1.0 | -1300 | -1750 | -4800 | 10 | 220 | 160 | 2 |
| SC052 | Thalamus | right | 2021-01-28 | 4 | 3B 1.0 | -1200 | 1800 | -4800 | 10 | 340 | -45 | 3 |

| | | | | | | | | | | | | |
|---|---|---|---|---|---|---|---|---|---|---|---|---|
| SC052 | Thalamus | right | 2021-01-28 | 5 | 3B 1.0 | -1900 | 800 | -4800 | 10 | 340 | -45 | 4 |
| SC052 | ALM | left | 2021-01-29 | 1 | 3B 1.0 | 2400 | -1600 | -3500 | 12 | 135 | -135 | 0 |
| SC052 | ALM | right | 2021-01-29 | 2 | 3B 1.0 | 2400 | 1500 | -3500 | 12 | 45 | -45 | 1 |
| SC052 | Thalamus | left | 2021-01-29 | 3 | 3B 1.0 | -1200 | -1800 | -4800 | 10 | 220 | 160 | 2 |
| SC052 | Thalamus | right | 2021-01-29 | 4 | 3B 1.0 | -1300 | 1800 | -4800 | 10 | 340 | 10 | 3 |
| SC052 | Thalamus | left | 2021-01-29 | 5 | 3B 1.0 | -1900 | -800 | -4800 | 10 | 220 | 160 | 4 |
| SC052 | ALM | left | 2021-01-30 | 1 | 3B 1.0 | 2400 | -1400 | -3600 | 12 | 135 | -135 | 0 |
| SC052 | ALM | right | 2021-01-30 | 2 | 3B 1.0 | 2400 | 1600 | -3700 | 12 | 45 | -45 | 1 |
| SC052 | BLA | left | 2021-01-30 | 3 | 3B 2.0 MS | -1500 | -4250 | -5100 | 8 | 180 | 180 | 2 |
| SC052 | BLA | right | 2021-01-30 | 4 | 3B 2.0 MS | -1500 | 4250 | -5150 | 8 | 0 | 0 | 3 |
| SC053 | ALM | left | 2021-02-18 | 1 | 3B 1.0 | 2500 | -1500 | -3300 | 15 | 120 | -135 | 0 |
| SC053 | Thalamus | left | 2021-02-18 | 2 | 3B 1.0 | -1200 | -600 | -4400 | 9 | 45 | -45 | 1 |
| SC053 | Striatum | left | 2021-02-18 | 3 | 3B 1.0 | 0 | -3300 | -4400 | 6 | 225 | -120 | 2 |
| SC053 | Midbrain | left | 2021-02-18 | 4 | 3B 1.0 | -4000 | -1900 | -4500 | 8 | 280 | 90 | 3 |
| SC053 | ALM | left | 2021-02-19 | 1 | 3B 2.0 SS | 2500 | -1600 | -2600 | 15 | 120 | -135 | 0 |
| SC053 | Thalamus | left | 2021-02-19 | 2 | 3B 1.0 | -1200 | -800 | -4800 | 10 | 0 | 0 | 1 |
| SC053 | Cortex | left | 2021-02-19 | 3 | 3B 1.0 | -1200 | -4400 | -3600 | 5 | 225 | -120 | 2 |
| SC053 | ALM | left | 2021-02-20 | 1 | 3B 2.0 SS | 2400 | -1600 | -2600 | 15 | 120 | -135 | 0 |
| SC053 | Thalamus | left | 2021-02-20 | 2 | 3B 1.0 | -1800 | -800 | -4800 | 10 | 315 | 0 | 1 |
| SC053 | Cortex | left | 2021-02-20 | 3 | 3B 1.0 | -1300 | -4400 | -4100 | 5 | 225 | -120 | 2 |
| SC053 | Midbrain | left | 2021-02-20 | 4 | 3B 1.0 | -4050 | -1800 | -4800 | 8 | 280 | 90 | 3 |
| SC053 | ALM | left | 2021-02-21 | 1 | 3B 2.0 SS | 2300 | -1700 | -2650 | 17.5 | 135 | -135 | 0 |
| SC053 | Thalamus | left | 2021-02-21 | 2 | 3B 1.0 | -1400 | -800 | -4800 | 10 | 0 | 0 | 1 |
| SC053 | Cortex | left | 2021-02-21 | 3 | 3B 1.0 | -1200 | -4300 | -4000 | 5 | 225 | -120 | 2 |

| | | | | | | | | | | | | |
|---|---|---|---|---|---|---|---|---|---|---|---|---|
| SC053 | Midbrain | left | 2021-02-21 | 4 | 3B 1.0 | -4000 | -1750 | -4600 | 10 | 280 | 90 | 3 |
| SC053 | ALM | left | 2021-02-22 | 1 | 3B 2.0 SS | 2400 | -1700 | -2600 | 17.5 | 135 | -135 | 0 |
| SC053 | BLA | left | 2021-02-22 | 2 | 3B 1.0 | -1300 | -4300 | -4800 | 15 | 225 | -120 | 1 |
| SC053 | Midbrain | left | 2021-02-22 | 3 | 3B 1.0 | -4000 | -1800 | -4850 | 10 | 315 | 0 | 2 |
| SC053 | Thalamus | right | 2021-02-23 | 1 | 3B 1.0 | -1200 | 1000 | -5100 | 10 | 170 | -135 | 0 |
| SC053 | ALM | right | 2021-02-23 | 2 | 3B 2.0 SS | 2500 | 1400 | -2650 | 15 | 45 | -45 | 1 |
| SC053 | Midbrain | right | 2021-02-23 | 3 | 3B 1.0 | -3900 | 1900 | -4700 | 10 | 260 | 90 | 2 |
| SC053 | Thalamus | right | 2021-02-24 | 1 | 3B 1.0 | -1200 | 1300 | -5200 | 10 | 190 | 135 | 0 |
| SC053 | ALM | right | 2021-02-24 | 2 | 3B 1.0 | 2500 | 1500 | -3500 | 16 | 45 | -45 | 1 |
| SC053 | Midbrain | right | 2021-02-24 | 3 | 3B 1.0 | -3800 | 1900 | -5000 | 10 | 260 | 90 | 2 |
| SC053 | Striatum | right | 2021-02-24 | 4 | 3B 2.0 SS | 0 | 3800 | -3400 | 6 | 340 | 0 | 3 |
| SC053 | Thalamus | right | 2021-02-25 | 1 | 3B 1.0 | -1400 | 1300 | -5300 | 9 | 190 | 135 | 0 |
| SC053 | ALM | right | 2021-02-25 | 2 | 3B 1.0 | 2400 | 1600 | -3500 | 15 | 45 | -45 | 1 |
| SC053 | Midbrain | right | 2021-02-25 | 3 | 3B 1.0 | -3800 | 1750 | -5000 | 10.5 | 260 | 90 | 2 |
| SC053 | Striatum | right | 2021-02-25 | 4 | 3B 2.0 SS | 200 | 3600 | -4000 | 12 | 0 | 0 | 3 |
| SC053 | Thalamus | right | 2021-02-26 | 1 | 3B 1.0 | -1400 | 1000 | -4450 | 10 | 190 | 135 | 0 |
| SC053 | ALM | right | 2021-02-26 | 2 | 3B 1.0 | 2500 | 1650 | -3200 | 15 | 45 | -45 | 1 |
| SC053 | Midbrain | right | 2021-02-26 | 3 | 3B 1.0 | -3900 | 1750 | -5050 | 11 | 260 | 90 | 2 |
| SC053 | Striatum | right | 2021-02-26 | 4 | 3B 2.0 SS | 400 | 3700 | -3500 | 12 | 0 | 0 | 3 |
| SC053 | Striatum | right | 2021-02-28 | 1 | 3B 2.0 SS | 0 | 3550 | -3900 | 15 | 80 | -45 | 0 |
| SC053 | Midbrain | left | 2021-02-28 | 2 | 3B 2.0 MS | -4000 | -2500 | -3200 | 20 | 200 | 180 | 1 |
| SC053 | Striatum | left | 2021-02-28 | 3 | 3B 2.0 MS | 0 | -3000 | -4200 | 20 | 350 | 90 | 2 |
| SC050 | ALM | left | 2021-02-25 | 1 | 3B 1.0 | 2500 | -1500 | -3200 | 15 | 135 | -135 | 0 |
| SC050 | ALM | right | 2021-02-25 | 2 | 3B 1.0 | 2500 | 1500 | -3200 | 15 | 45 | -45 | 1 |

| | | | | | | | | | | | | |
|---|---|---|---|---|---|---|---|---|---|---|---|---|
| SC050 | Thalamus | left | 2021-02-25 | 3 | 3B 2.0 MS | -1200 | -1600 | -4900 | 15 | 200 | 80 | 2 |
| SC050 | Thalamus | right | 2021-02-25 | 4 | 3B 2.0 MS | -1200 | 1500 | -4700 | 15 | 340 | 100 | 3 |
| SC050 | ALM | left | 2021-02-26 | 1 | 3B 1.0 | 2500 | -1600 | -3300 | 15 | 135 | -135 | 0 |
| SC050 | ALM | right | 2021-02-26 | 2 | 3B 1.0 | 2500 | 1600 | -3300 | 15 | 45 | -45 | 1 |
| SC050 | Thalamus | left | 2021-02-26 | 3 | 3B 2.0 MS | -1400 | -1600 | -4500 | 15 | 200 | 80 | 2 |
| SC050 | Thalamus | right | 2021-02-26 | 4 | 3B 2.0 MS | -1400 | 1500 | -4500 | 15 | 340 | 100 | 3 |
| SC050 | ALM | left | 2021-02-27 | 1 | 3B 1.0 | 2400 | -1700 | -3300 | 15 | 135 | -135 | 0 |
| SC050 | ALM | right | 2021-02-27 | 2 | 3B 1.0 | 2400 | 1700 | -3300 | 15 | 45 | -45 | 1 |
| SC050 | Thalamus | left | 2021-02-27 | 3 | 3B 2.0 MS | -1700 | -1600 | -4450 | 15 | 200 | 80 | 2 |
| SC050 | Thalamus | right | 2021-02-27 | 4 | 3B 2.0 MS | -1800 | 1500 | -4400 | 15 | 340 | 100 | 3 |
| SC050 | ALM | left | 2021-03-01 | 1 | 3B 1.0 | 2400 | -1500 | -3200 | 15 | 135 | -135 | 0 |
| SC050 | ALM | right | 2021-03-01 | 2 | 3B 1.0 | 2400 | 1500 | -3200 | 15 | 45 | -45 | 1 |
| SC050 | Midbrain | left | 2021-03-01 | 3 | 3B 2.0 MS | -3800 | -2750 | -3200 | 20 | 200 | 80 | 2 |
| SC050 | Striatum | left | 2021-03-01 | 4 | 3B 2.0 MS | 0 | -2100 | -4400 | 20 | 340 | 90 | 3 |
| SC050 | ALM | left | 2021-03-02 | 1 | 3B 1.0 | 2450 | -1400 | -3300 | 15 | 135 | -135 | 0 |
| SC050 | ALM | right | 2021-03-02 | 2 | 3B 1.0 | 2400 | 1400 | -3300 | 15 | 45 | -45 | 1 |
| SC050 | Midbrain | left | 2021-03-02 | 3 | 3B 2.0 MS | -3900 | -2750 | -3000 | 20 | 200 | 90 | 2 |
| SC050 | Striatum | left | 2021-03-02 | 4 | 3B 2.0 MS | -200 | -2100 | -4300 | 20 | 340 | 110 | 3 |
| SC050 | ALM | left | 2021-03-03 | 1 | 3B 1.0 | 2300 | -1500 | -3300 | 17.5 | 135 | -135 | 0 |
| SC050 | ALM | right | 2021-03-03 | 2 | 3B 1.0 | 2300 | 1500 | -3300 | 17.5 | 45 | -45 | 1 |
| SC050 | Striatum | right | 2021-03-03 | 3 | 3B 2.0 MS | 0 | 2100 | -4200 | 20 | 200 | 90 | 2 |
| SC050 | Midbrain | right | 2021-03-03 | 4 | 3B 2.0 MS | -3800 | 2750 | -3100 | 15 | 340 | 90 | 3 |
| SC050 | Striatum | right | 2021-03-04 | 1 | 3B 1.0 | -50 | 2400 | -4500 | 17.5 | 45 | -45 | 0 |
| SC050 | Striatum | right | 2021-03-04 | 2 | 3B 2.0 MS | -200 | 2100 | -4200 | 20 | 200 | 90 | 1 |

| | | | | | | | | | | | | |
|---|---|---|---|---|---|---|---|---|---|---|---|---|
| SC050 | Midbrain | right | 2021-03-04 | 3 | 3B 2.0 MS | -3900 | 2750 | -3300 | 15 | 340 | 90 | 2 |
| SC060 | Cortex | left | 2021-03-18 | 1 | 3B 2.0 SS | -2200 | -4500 | -3600 | 10 | 135 | 160 | 0 |
| SC060 | ALM | left | 2021-03-18 | 2 | 3B 2.0 SS | 2500 | -1500 | -2650 | 20 | 80 | 80 | 1 |
| SC060 | Midbrain | left | 2021-03-18 | 3 | 3B 2.0 MS | -3600 | -2500 | -3300 | 20 | 200 | 85 | 2 |
| SC060 | Striatum | left | 2021-03-18 | 4 | 3B 2.0 MS | 0 | -1800 | -4350 | 20 | 350 | 90 | 3 |
| SC060 | Cortex | left | 2021-03-19 | 1 | 3B 2.0 SS | -2250 | -4550 | -3600 | 10 | 135 | 160 | 0 |
| SC060 | ALM | left | 2021-03-19 | 2 | 3B 2.0 SS | 2450 | -1550 | -2650 | 20 | 80 | 80 | 1 |
| SC060 | Midbrain | left | 2021-03-19 | 3 | 3B 2.0 MS | -3800 | -2450 | -3350 | 20 | 200 | 85 | 2 |
| SC060 | Striatum | left | 2021-03-19 | 4 | 3B 2.0 MS | 300 | -1800 | -4250 | 20 | 350 | 90 | 3 |
| SC060 | Cortex | left | 2021-03-20 | 1 | 3B 2.0 SS | -2300 | -4500 | -3600 | 10 | 135 | 160 | 0 |
| SC060 | ALM | left | 2021-03-20 | 2 | 3B 2.0 SS | 2400 | -1500 | -2650 | 20 | 80 | 80 | 1 |
| SC060 | Midbrain | left | 2021-03-20 | 3 | 3B 2.0 MS | -4000 | -2450 | -3350 | 20 | 200 | 85 | 2 |
| SC060 | Thalamus | left | 2021-03-20 | 4 | 3B 2.0 MS | -1800 | -800 | -4700 | 8 | 0 | -90 | 3 |
| SC060 | ALM | right | 2021-03-22 | 1 | 3B 2.0 SS | 2500 | 1500 | -2600 | 20 | 100 | -100 | 0 |
| SC060 | Cortex | right | 2021-03-22 | 2 | 3B 2.0 SS | -2400 | 4550 | -3600 | 10 | 60 | -20 | 1 |
| SC060 | Striatum | right | 2021-03-22 | 3 | 3B 2.0 MS | 0 | 1800 | -4400 | 20 | 200 | 90 | 2 |
| SC060 | Midbrain | right | 2021-03-22 | 4 | 3B 2.0 MS | -3700 | 1600 | -3200 | 20 | 350 | 90 | 3 |
| SC060 | ALM | right | 2021-03-23 | 1 | 3B 2.0 SS | 2450 | 1550 | -2700 | 20 | 100 | -100 | 0 |
| SC060 | Cortex | right | 2021-03-23 | 2 | 3B 2.0 SS | -2450 | 4550 | -3550 | 10 | 60 | -20 | 1 |
| SC060 | Striatum | right | 2021-03-23 | 3 | 3B 2.0 MS | 200 | 1800 | -4300 | 20 | 200 | 90 | 2 |
| SC060 | Midbrain | right | 2021-03-23 | 4 | 3B 2.0 MS | -4000 | 1600 | -3200 | 20 | 350 | 90 | 3 |
| SC060 | ALM | left | 2021-03-24 | 1 | 3B 2.0 SS | 2400 | -1650 | -2750 | 15 | 135 | -135 | 0 |
| SC060 | ALM | right | 2021-03-24 | 2 | 3B 2.0 SS | 2400 | 1600 | -2700 | 15 | 45 | -45 | 1 |
| SC060 | Thalamus | left | 2021-03-24 | 3 | 3B 2.0 MS | -1700 | -900 | -4350 | 10 | 190 | 90 | 2 |

| | | | | | | | | | | | | |
|---|---|---|---|---|---|---|---|---|---|---|---|---|
| SC061 | Cortex | left | 2021-03-18 | 1 | 3B 2.0 SS | -2250 | -4500 | -3600 | 10 | 135 | 160 | 0 |
| SC061 | ALM | left | 2021-03-18 | 2 | 3B 2.0 SS | 2500 | -1700 | -2650 | 20 | 80 | 80 | 1 |
| SC061 | Midbrain | left | 2021-03-18 | 3 | 3B 2.0 MS | -3900 | -2500 | -3300 | 20 | 200 | 85 | 2 |
| SC061 | Striatum | left | 2021-03-18 | 4 | 3B 2.0 MS | 0 | -1800 | -4300 | 20 | 350 | 90 | 3 |
| SC061 | Cortex | left | 2021-03-19 | 1 | 3B 1.0 | -2350 | -4500 | -3600 | 10 | 120 | 160 | 0 |
| SC061 | Midbrain | left | 2021-03-19 | 2 | 3B 1.0 | -4000 | -1700 | -4300 | 20 | 200 | 100 | 1 |
| SC061 | Striatum | left | 2021-03-19 | 3 | 3B 1.0 | 400 | -2300 | -4700 | 20 | 0 | 0 | 2 |
| SC061 | Striatum | left | 2021-03-19 | 4 | 3B 1.0 | 400 | -1500 | -4700 | 20 | 0 | 0 | 3 |
| SC061 | ALM | right | 2021-03-21 | 1 | 3B 2.0 SS | 2500 | 1500 | -2600 | 20 | 100 | -100 | 0 |
| SC061 | Cortex | right | 2021-03-21 | 2 | 3B 2.0 SS | -2300 | 4500 | -3600 | 10 | 50 | -20 | 1 |
| SC061 | Striatum | right | 2021-03-21 | 3 | 3B 2.0 MS | 0 | 1800 | -4200 | 20 | 200 | 90 | 2 |
| SC061 | Midbrain | right | 2021-03-21 | 4 | 3B 2.0 MS | -3950 | 1700 | -3250 | 20 | 350 | 90 | 3 |
| SC061 | ALM | right | 2021-03-22 | 1 | 3B 2.0 SS | 2450 | 1550 | -2700 | 20 | 100 | -100 | 0 |
| SC061 | Cortex | right | 2021-03-22 | 2 | 3B 2.0 SS | -2350 | 4500 | -3500 | 10 | 60 | -20 | 1 |
| SC061 | Striatum | right | 2021-03-22 | 3 | 3B 2.0 MS | 200 | 1800 | -4300 | 20 | 200 | 90 | 2 |
| SC061 | Midbrain | right | 2021-03-22 | 4 | 3B 2.0 MS | -4050 | 1650 | -3200 | 20 | 350 | 90 | 3 |
| SC061 | Striatum | left | 2021-03-23 | 1 | 3B 2.0 SS | 0 | -3500 | -4400 | 15 | 135 | -160 | 0 |
| SC061 | ALM | right | 2021-03-23 | 2 | 3B 2.0 SS | 2400 | 1600 | -2650 | 15 | 45 | -45 | 1 |
| SC061 | Thalamus | left | 2021-03-23 | 3 | 3B 2.0 MS | -2200 | -1800 | -4350 | 20 | 190 | 90 | 2 |
| SC061 | Thalamus | right | 2021-03-23 | 4 | 3B 2.0 MS | -1800 | 2200 | -4500 | 20 | 350 | 90 | 3 |
| SC061 | ALM | right | 2021-03-24 | 1 | 3B 2.0 SS | 2350 | 1450 | -2600 | 15 | 45 | -45 | 0 |
| SC061 | Thalamus | left | 2021-03-24 | 2 | 3B 2.0 MS | -2300 | -1850 | -4300 | 20 | 190 | 90 | 1 |
| SC061 | Thalamus | right | 2021-03-24 | 3 | 3B 2.0 MS | -1900 | 2200 | -4400 | 20 | 350 | 90 | 2 |
| SC066 | ALM | left | 2021-04-13 | 1 | 3B 2.0 SS | 2500 | -1600 | -2500 | 15 | 135 | -135 | 0 |

| | | | | | | | | | | | | |
|---|---|---|---|---|---|---|---|---|---|---|---|---|
| SC066 | Striatum | left | 2021-04-13 | 2 | 3B 2.0 MS | -200 | -1800 | -4050 | 18 | 5 | 90 | 1 |
| SC066 | Midbrain | left | 2021-04-13 | 3 | 3B 1.0 | -4000 | -1800 | -4700 | 10 | 260 | 180 | 2 |
| SC066 | Medulla | right | 2021-04-13 | 4 | 3B 1.0 | -6500 | 1000 | -4800 | 8 | 280 | 0 | 3 |
| SC066 | ALM | left | 2021-04-14 | 1 | 3B 2.0 SS | 2500 | -1500 | -2600 | 16 | 135 | -135 | 0 |
| SC066 | Striatum | left | 2021-04-14 | 2 | 3B 2.0 MS | -400 | -1800 | -4000 | 18 | 5 | 90 | 1 |
| SC066 | Midbrain | left | 2021-04-14 | 3 | 3B 1.0 | -4000 | -1700 | -4700 | 12 | 260 | 180 | 2 |
| SC066 | Medulla | right | 2021-04-14 | 4 | 3B 1.0 | -6500 | 1800 | -4800 | 10 | 280 | 0 | 3 |
| SC066 | ALM | left | 2021-04-15 | 1 | 3B 2.0 SS | 2400 | -1500 | -2600 | 16 | 135 | -135 | 0 |
| SC066 | Striatum | left | 2021-04-15 | 2 | 3B 2.0 SS | -50 | -2000 | -4500 | 18 | 5 | 90 | 1 |
| SC066 | Midbrain | left | 2021-04-15 | 3 | 3B 1.0 | -4000 | -1600 | -4900 | 12 | 260 | 180 | 2 |
| SC066 | Medulla | right | 2021-04-15 | 4 | 3B 1.0 | -6500 | 1900 | -4800 | 10 | 280 | 0 | 3 |
| SC066 | ALM | left | 2021-04-16 | 1 | 3B 2.0 SS | 2400 | -1600 | -2700 | 16 | 135 | -135 | 0 |
| SC066 | Striatum | left | 2021-04-16 | 2 | 3B 2.0 SS | -50 | -1900 | -4600 | 18 | 5 | 90 | 1 |
| SC066 | Midbrain | left | 2021-04-16 | 3 | 3B 1.0 | -4050 | -1800 | -4700 | 12 | 260 | 180 | 2 |
| SC066 | Medulla | right | 2021-04-16 | 4 | 3B 1.0 | -6500 | 2000 | -4700 | 10 | 280 | 0 | 3 |
| SC066 | Striatum | right | 2021-04-19 | 1 | 3B 2.0 SS | 50 | 1800 | -4200 | 18 | 180 | 90 | 0 |
| SC066 | ALM | right | 2021-04-19 | 2 | 3B 2.0 SS | 2500 | 1500 | -2750 | 18 | 45 | -45 | 1 |
| SC066 | Medulla | left | 2021-04-19 | 3 | 3B 1.0 | -6600 | -1100 | -4800 | 12 | 260 | 180 | 2 |
| SC066 | Midbrain | right | 2021-04-19 | 4 | 3B 1.0 | -4050 | 1700 | -4900 | 14 | 280 | 0 | 3 |
| SC066 | Medulla | left | 2021-04-19 | 5 | 3B 1.0 | -6600 | -2200 | -4800 | 12 | 260 | 180 | 4 |
| SC066 | Striatum | right | 2021-04-20 | 1 | 3B 2.0 SS | 50 | 1600 | -4350 | 18 | 180 | 90 | 0 |
| SC066 | ALM | right | 2021-04-20 | 2 | 3B 2.0 SS | 2500 | 1500 | -2700 | 18 | 45 | -45 | 1 |
| SC066 | Medulla | left | 2021-04-20 | 3 | 3B 1.0 | -6550 | -1100 | -4700 | 12 | 260 | 180 | 2 |
| SC066 | Midbrain | right | 2021-04-20 | 4 | 3B 1.0 | -4000 | 1700 | -4800 | 14 | 280 | 0 | 3 |

| | | | | | | | | | | | | |
|---|---|---|---|---|---|---|---|---|---|---|---|---|
| SC066 | Medulla | left | 2021-04-20 | 5 | 3B 1.0 | -6550 | -2200 | -4700 | 12 | 260 | 180 | 4 |
| SC066 | Striatum | right | 2021-04-21 | 1 | 3B 2.0 SS | 50 | 1700 | -4400 | 18 | 180 | 90 | 0 |
| SC066 | ALM | right | 2021-04-21 | 2 | 3B 2.0 SS | 2400 | 1500 | -2700 | 18 | 45 | -45 | 1 |
| SC066 | Medulla | left | 2021-04-21 | 3 | 3B 1.0 | -6550 | -2000 | -4700 | 12 | 260 | 180 | 2 |
| SC066 | Midbrain | right | 2021-04-21 | 4 | 3B 1.0 | -4000 | 1600 | -5000 | 14 | 280 | 0 | 3 |
| SC067 | ALM | left | 2021-04-13 | 1 | 3B 2.0 SS | 2450 | -1600 | -2550 | 15 | 135 | -135 | 0 |
| SC067 | Striatum | left | 2021-04-13 | 2 | 3B 2.0 MS | -200 | -1800 | -4000 | 18 | 5 | 90 | 1 |
| SC067 | Midbrain | left | 2021-04-13 | 3 | 3B 1.0 | -4000 | -1800 | -4700 | 10 | 260 | 180 | 2 |
| SC067 | Medulla | right | 2021-04-13 | 4 | 3B 1.0 | -6500 | 1000 | -4800 | 8 | 280 | 0 | 3 |
| SC067 | Medulla | right | 2021-04-13 | 5 | 3B 1.0 | -6500 | 2200 | -4800 | 8 | 280 | 0 | 4 |
| SC067 | ALM | left | 2021-04-14 | 1 | 3B 2.0 SS | 2400 | -1600 | -2500 | 16 | 135 | -135 | 0 |
| SC067 | Striatum | left | 2021-04-14 | 2 | 3B 2.0 MS | -200 | -1800 | -4000 | 18 | 5 | 90 | 1 |
| SC067 | Midbrain | left | 2021-04-14 | 3 | 3B 1.0 | -4050 | -1700 | -5000 | 12 | 260 | 180 | 2 |
| SC067 | Medulla | right | 2021-04-14 | 4 | 3B 1.0 | -6550 | 1000 | -4850 | 10 | 280 | 0 | 3 |
| SC067 | Medulla | right | 2021-04-14 | 5 | 3B 1.0 | -6550 | 2200 | -4850 | 10 | 280 | 0 | 4 |
| SC067 | ALM | left | 2021-04-15 | 1 | 3B 2.0 SS | 2400 | -1500 | -2700 | 16 | 135 | -135 | 0 |
| SC067 | Striatum | left | 2021-04-15 | 2 | 3B 2.0 SS | -150 | -2000 | -4550 | 18 | 5 | 90 | 1 |
| SC067 | Midbrain | left | 2021-04-15 | 3 | 3B 1.0 | -4050 | -1600 | -4900 | 12 | 260 | 180 | 2 |
| SC067 | Medulla | right | 2021-04-15 | 4 | 3B 1.0 | -6550 | 1800 | -4800 | 10 | 280 | 0 | 3 |
| SC067 | Striatum | right | 2021-04-18 | 1 | 3B 2.0 SS | 0 | 1800 | -4550 | 18 | 180 | 90 | 0 |
| SC067 | ALM | right | 2021-04-18 | 2 | 3B 2.0 SS | 2500 | 1500 | -2700 | 18 | 45 | -45 | 1 |
| SC067 | Medulla | left | 2021-04-18 | 3 | 3B 1.0 | -6600 | -1100 | -4550 | 12 | 260 | 180 | 2 |
| SC067 | Midbrain | right | 2021-04-18 | 4 | 3B 1.0 | -4050 | 1700 | -4850 | 12 | 280 | 0 | 3 |
| SC067 | Medulla | left | 2021-04-18 | 5 | 3B 1.0 | -6600 | -2200 | -4550 | 12 | 260 | 180 | 4 |

| | | | | | | | | | | | | |
|---|---|---|---|---|---|---|---|---|---|---|---|---|
| SC067 | Striatum | right | 2021-04-19 | 1 | 3B 2.0 SS | 50 | 1800 | -4350 | 18 | 180 | 90 | 0 |
| SC067 | ALM | right | 2021-04-19 | 2 | 3B 2.0 SS | 2500 | 1500 | -2700 | 18 | 45 | -45 | 1 |
| SC067 | Medulla | left | 2021-04-19 | 3 | 3B 1.0 | -6600 | -1100 | -4650 | 12 | 260 | 180 | 2 |
| SC067 | Midbrain | right | 2021-04-19 | 4 | 3B 1.0 | -4050 | 1700 | -4900 | 14 | 280 | 0 | 3 |
| SC067 | Medulla | left | 2021-04-19 | 5 | 3B 1.0 | -6600 | -2200 | -4650 | 12 | 260 | 180 | 4 |
| SC067 | Striatum | right | 2021-04-20 | 1 | 3B 2.0 SS | 50 | 1500 | -4500 | 18 | 180 | 90 | 0 |
| SC067 | ALM | right | 2021-04-20 | 2 | 3B 2.0 SS | 2400 | 1600 | -2700 | 18 | 45 | -45 | 1 |
| SC067 | Medulla | left | 2021-04-20 | 3 | 3B 1.0 | -6600 | -1100 | -4700 | 12 | 260 | 180 | 2 |
| SC067 | Midbrain | right | 2021-04-20 | 4 | 3B 1.0 | -3950 | 1750 | -5100 | 14 | 280 | 0 | 3 |
| SC067 | Medulla | left | 2021-04-20 | 5 | 3B 1.0 | -6600 | -2200 | -4700 | 12 | 260 | 180 | 4 |
| SC064 | ALM | left | 2021-04-27 | 1 | 3B 2.0 SS | 2550 | -1500 | -2600 | 18 | 135 | -135 | 0 |
| SC064 | Thalamus | left | 2021-04-27 | 2 | 3B 1.0 | -1200 | -700 | -4200 | 6 | 0 | 90 | 1 |
| SC064 | Cortex | left | 2021-04-27 | 3 | 3B 2.0 MS | -1800 | -4200 | -3300 | 6 | 180 | 180 | 2 |
| SC064 | Midbrain | left | 2021-04-27 | 4 | 3B 1.0 | -3800 | -2000 | -4500 | 10 | 270 | 90 | 3 |
| SC064 | Thalamus | left | 2021-04-27 | 5 | 3B 1.0 | -2000 | -700 | -4200 | 6 | 0 | 90 | 4 |
| SC064 | ALM | left | 2021-04-28 | 1 | 3B 2.0 SS | 2500 | -1450 | -2600 | 18 | 135 | -135 | 0 |
| SC064 | Thalamus | left | 2021-04-28 | 2 | 3B 2.0 MS | -1200 | -850 | -4100 | 6 | 0 | 90 | 1 |
| SC064 | Cortex | left | 2021-04-28 | 3 | 3B 2.0 SS | -1900 | -4150 | -3450 | 6 | 180 | 180 | 2 |
| SC064 | Midbrain | left | 2021-04-28 | 4 | 3B 1.0 | -3800 | -1900 | -4550 | 12 | 270 | 90 | 3 |
| SC064 | ALM | left | 2021-04-29 | 1 | 3B 2.0 SS | 2500 | -1550 | -2600 | 18 | 135 | -135 | 0 |
| SC064 | Thalamus | left | 2021-04-29 | 2 | 3B 2.0 SS | -1600 | -700 | -4300 | 6 | 0 | 90 | 1 |
| SC064 | Cortex | left | 2021-04-29 | 3 | 3B 2.0 SS | -1900 | -4250 | -3500 | 6 | 180 | 180 | 2 |
| SC064 | Midbrain | left | 2021-04-29 | 4 | 3B 1.0 | -3800 | -1850 | -4700 | 14 | 270 | 90 | 3 |
| SC064 | ALM | left | 2021-04-30 | 1 | 3B 2.0 SS | 2400 | -1600 | -2600 | 18 | 135 | -135 | 0 |

| | | | | | | | | | | | | |
|---|---|---|---|---|---|---|---|---|---|---|---|---|
| SC064 | Thalamus | left | 2021-04-30 | 2 | 3B 1.0 | -1200 | -1200 | -4700 | 6 | 0 | 90 | 1 |
| SC064 | BLA | left | 2021-04-30 | 3 | 3B 2.0 SS | -1700 | -4150 | -4850 | 12 | 180 | 180 | 2 |
| SC064 | Midbrain | left | 2021-04-30 | 4 | 3B 1.0 | -3750 | -1850 | -4900 | 15 | 270 | 90 | 3 |
| SC064 | Thalamus | left | 2021-04-30 | 5 | 3B 1.0 | -2000 | -1200 | -4700 | 6 | 0 | 90 | 4 |
| SC064 | Striatum | right | 2021-05-04 | 1 | 3B 2.0 SS | 0 | 1700 | -4400 | 18 | 180 | 90 | 0 |
| SC064 | ALM | right | 2021-05-04 | 2 | 3B 2.0 SS | 2500 | 1500 | -2650 | 16 | 45 | -45 | 1 |
| SC064 | Midbrain | right | 2021-05-04 | 3 | 3B 1.0 | -3800 | 1700 | -5250 | 13 | 270 | 90 | 2 |
| SC064 | Cortex | right | 2021-05-04 | 4 | 3B 2.0 SS | -1200 | 4200 | -3700 | 8 | 0 | 0 | 3 |
| SC064 | Striatum | right | 2021-05-05 | 1 | 3B 2.0 SS | 0 | 1800 | -4500 | 18 | 180 | 90 | 0 |
| SC064 | ALM | right | 2021-05-05 | 2 | 3B 2.0 SS | 2450 | 1450 | -2700 | 16 | 45 | -45 | 1 |
| SC064 | Midbrain | right | 2021-05-05 | 3 | 3B 1.0 | -3800 | 1800 | -4900 | 13 | 270 | 90 | 2 |
| SC064 | Cortex | right | 2021-05-05 | 4 | 3B 2.0 SS | -1250 | 4200 | -3600 | 8 | 0 | 0 | 3 |
| SC064 | Striatum | right | 2021-05-06 | 1 | 3B 2.0 MS | -100 | 1750 | -4200 | 18 | 180 | 90 | 0 |
| SC064 | ALM | right | 2021-05-06 | 2 | 3B 2.0 SS | 2450 | 1550 | -2700 | 16 | 45 | -45 | 1 |
| SC064 | Midbrain | right | 2021-05-06 | 3 | 3B 1.0 | -3850 | 1750 | -5300 | 14 | 270 | 90 | 2 |
| SC064 | Cortex | right | 2021-05-06 | 4 | 3B 2.0 SS | -1300 | 4200 | -3600 | 6 | 0 | 0 | 3 |
| SC064 | Striatum | left | 2021-05-07 | 1 | 3B 2.0 MS | -50 | -3050 | -4000 | 6 | 180 | 180 | 0 |
| SC064 | ALM | right | 2021-05-07 | 2 | 3B 2.0 SS | 2400 | 1450 | -2700 | 16 | 45 | -45 | 1 |
| SC064 | Thalamus | right | 2021-05-07 | 3 | 3B 1.0 | -2200 | 700 | -4400 | 15 | 270 | 90 | 2 |
| SC064 | BLA | right | 2021-05-07 | 4 | 3B 2.0 SS | -1200 | 4100 | -4000 | 10 | 0 | 0 | 3 |
| SC064 | Thalamus | right | 2021-05-07 | 5 | 3B 1.0 | -3000 | 700 | -4400 | 15 | 270 | 90 | 4 |
| SC064 | Striatum | left | 2021-05-08 | 1 | 3B 2.0 MS | 500 | -3000 | -4300 | 8 | 180 | 180 | 0 |
| SC064 | ALM | right | 2021-05-08 | 2 | 3B 2.0 SS | 2400 | 1550 | -2700 | 16 | 45 | -45 | 1 |
| SC064 | Thalamus | right | 2021-05-08 | 3 | 3B 1.0 | -2500 | 800 | -4500 | 15 | 270 | 90 | 2 |

| | | | | | | | | | | | | |
|---|---|---|---|---|---|---|---|---|---|---|---|---|
| SC064 | BLA | right | 2021-05-08 | 4 | 3B 2.0 SS | -1300 | 4100 | -4400 | 10 | 0 | 0 | 3 |
| SC065 | Striatum | right | 2021-05-04 | 1 | 3B 2.0 SS | 0 | 2000 | -4400 | 18 | 180 | 90 | 0 |
| SC065 | ALM | right | 2021-05-04 | 2 | 3B 2.0 SS | 2500 | 1500 | -2700 | 16 | 60 | -45 | 1 |
| SC065 | Midbrain | right | 2021-05-04 | 3 | 3B 1.0 | -3750 | 1800 | -5250 | 13 | 270 | 90 | 2 |
| SC065 | Cortex | right | 2021-05-04 | 4 | 3B 2.0 SS | -1200 | 4300 | -3600 | 6 | 0 | 0 | 3 |
| SC065 | Striatum | right | 2021-05-05 | 1 | 3B 2.0 SS | 0 | 2100 | -4700 | 18 | 180 | 90 | 0 |
| SC065 | ALM | right | 2021-05-05 | 2 | 3B 2.0 SS | 2450 | 1550 | -2700 | 16 | 60 | -45 | 1 |
| SC065 | Midbrain | right | 2021-05-05 | 3 | 3B 1.0 | -3750 | 1700 | -5000 | 13 | 270 | 90 | 2 |
| SC065 | Cortex | right | 2021-05-05 | 4 | 3B 2.0 SS | -1250 | 4250 | -3500 | 6 | 0 | 0 | 3 |
| SC065 | Striatum | right | 2021-05-06 | 1 | 3B 2.0 SS | -100 | 1900 | -4300 | 18 | 180 | 90 | 0 |
| SC065 | ALM | right | 2021-05-06 | 2 | 3B 2.0 SS | 2450 | 1450 | -2700 | 16 | 60 | -45 | 1 |
| SC065 | Midbrain | right | 2021-05-06 | 3 | 3B 1.0 | -3800 | 1800 | -5200 | 14 | 270 | 90 | 2 |
| SC065 | Cortex | right | 2021-05-06 | 4 | 3B 2.0 SS | -1300 | 4300 | -3600 | 6 | 0 | 0 | 3 |
| SC065 | Striatum | right | 2021-05-07 | 1 | 3B 2.0 SS | -100 | 2100 | -4200 | 16 | 180 | 90 | 0 |
| SC065 | Thalamus | right | 2021-05-07 | 2 | 3B 1.0 | -2000 | 600 | -4400 | 15 | 270 | 90 | 1 |
| SC065 | Striatum | right | 2021-05-09 | 1 | 3B 2.0 SS | -100 | 2200 | -4100 | 16 | 180 | 90 | 0 |
| SC065 | ALM | right | 2021-05-09 | 2 | 3B 2.0 SS | 2450 | 1600 | -2700 | 16 | 60 | -45 | 1 |
| SC065 | Thalamus | right | 2021-05-09 | 3 | 3B 1.0 | -3200 | 600 | -4700 | 21 | 270 | 90 | 2 |
| SC065 | BLA | right | 2021-05-09 | 4 | 3B 1.0 | -1300 | 4200 | -4500 | 10 | 0 | 0 | 3 |
| SC065 | ALM | left | 2021-05-10 | 1 | 3B 2.0 SS | 2500 | -1500 | -2600 | 16 | 125 | -135 | 0 |
| SC065 | Striatum | left | 2021-05-10 | 2 | 3B 2.0 MS | 0 | -2000 | -4200 | 17 | 0 | 90 | 1 |
| SC065 | Cortex | left | 2021-05-10 | 3 | 3B 2.0 SS | -2200 | -4300 | -3600 | 8 | 180 | 180 | 2 |
| SC065 | Midbrain | left | 2021-05-10 | 4 | 3B 1.0 | -3800 | -1800 | -5000 | 12 | 270 | 90 | 3 |
| SC065 | ALM | left | 2021-05-11 | 1 | 3B 2.0 SS | 2400 | -1400 | -2650 | 16 | 125 | -135 | 0 |

| | | | | | | | | | | | | |
|---|---|---|---|---|---|---|---|---|---|---|---|---|
| SC065 | Striatum | left | 2021-05-11 | 2 | 3B 2.0 MS | -100 | -2000 | -4000 | 17 | 0 | 90 | 1 |
| SC065 | Cortex | left | 2021-05-11 | 3 | 3B 2.0 SS | -2250 | -4300 | -3600 | 8 | 180 | 180 | 2 |
| SC065 | Midbrain | left | 2021-05-11 | 4 | 3B 1.0 | -3800 | -1700 | -5050 | 12 | 270 | 90 | 3 |
| SC065 | ALM | left | 2021-05-13 | 1 | 3B 2.0 SS | 2300 | -1600 | -2700 | 16 | 125 | -135 | 0 |
| SC065 | Thalamus | left | 2021-05-13 | 2 | 3B 2.0 MS | -1500 | -1200 | -4200 | 6 | 0 | -45 | 1 |
| SC065 | BLA | left | 2021-05-13 | 3 | 3B 2.0 SS | -2200 | -4200 | -4050 | 12 | 180 | 180 | 2 |
| SC065 | Midbrain | left | 2021-05-13 | 4 | 3B 1.0 | -3850 | -1750 | -4900 | 14 | 270 | 90 | 3 |
| SC065 | ALM | left | 2021-05-14 | 1 | 3B 2.0 SS | 2400 | -1350 | -2700 | 16 | 125 | -135 | 0 |
| SC065 | Thalamus | left | 2021-05-14 | 2 | 3B 1.0 | -1400 | -1200 | -4900 | 6 | 0 | 90 | 1 |
| SC065 | BLA | left | 2021-05-14 | 3 | 3B 2.0 SS | -2250 | -4200 | -5000 | 12 | 180 | 180 | 2 |
| SC065 | Midbrain | left | 2021-05-14 | 4 | 3B 1.0 | -3750 | -1700 | -5000 | 14 | 270 | 90 | 3 |

**Table 3 - Reconstructed neurons used for defining ALM projection zones**

Reconstructed neurons (in ALM and nearby areas) used for defining ALM projection zones. Neurons can be viewed at (https://ml-neuronbrowser.janelia.org/).

| ID | DOI |
|---|---|
| **AA0006** | 10.25378/janelia.5520223 |
| **AA0010** | 10.25378/janelia.5521600 |
| **AA0011** | 10.25378/janelia.5521615 |
| **AA0012** | 10.25378/janelia.5521618 |
| **AA0013** | 10.25378/janelia.5521621 |
| **AA0014** | 10.25378/janelia.5521624 |
| **AA0059** | 10.25378/janelia.5521780 |
| **AA0060** | 10.25378/janelia.5521783 |
| **AA0062** | 10.25378/janelia.5521789 |
| **AA0065** | 10.25378/janelia.5521801 |
| **AA0108** | 10.25378/janelia.5526703 |
| **AA0114** | 10.25378/janelia.5526721 |
| **AA0115** | 10.25378/janelia.5526724 |
| **AA0116** | 10.25378/janelia.5526727 |
| **AA0118** | 10.25378/janelia.5526733 |
| **AA0122** | 10.25378/janelia.5527240 |
| **AA0133** | 10.25378/janelia.5527273 |
| **AA0134** | 10.25378/janelia.5527276 |
| **AA0179** | 10.25378/janelia.5527438 |
| **AA0180** | 10.25378/janelia.5527441 |
| **AA0181** | 10.25378/janelia.5527444 |
| **AA0183** | 10.25378/janelia.5527450 |
| **AA0185** | 10.25378/janelia.5527456 |
| **AA0187** | 10.25378/janelia.5527465 |

| | |
|---|---|
| **AA0188** | 10.25378/janelia.5527468 |
| **AA0190** | 10.25378/janelia.5527474 |
| **AA0245** | 10.25378/janelia.5527657 |
| **AA0250** | 10.25378/janelia.5527678 |
| **AA0265** | 10.25378/janelia.5527738 |
| **AA0267** | 10.25378/janelia.5527747 |
| **AA0269** | 10.25378/janelia.5527753 |
| **AA0271** | 10.25378/janelia.5527762 |
| **AA0274** | 10.25378/janelia.5527774 |
| **AA0276** | 10.25378/janelia.5527780 |
| **AA0279** | 10.25378/janelia.5527792 |
| **AA0281** | 10.25378/janelia.5527798 |
| **AA0284** | 10.25378/janelia.5527807 |
| **AA0285** | 10.25378/janelia.5527810 |
| **AA0286** | 10.25378/janelia.5527813 |
| **AA0287** | 10.25378/janelia.5527816 |
| **AA0288** | n/a |
| **AA0291** | 10.25378/janelia.5527828 |
| **AA0324** | 10.25378/janelia.7613486 |
| **AA0327** | 10.25378/janelia.7613498 |
| **AA0328** | 10.25378/janelia.7613507 |
| **AA0329** | 10.25378/janelia.7613525 |
| **AA0332** | 10.25378/janelia.7613684 |
| **AA0333** | 10.25378/janelia.7613693 |
| **AA0390** | 10.25378/janelia.7614077 |
| **AA0394** | 10.25378/janelia.7614101 |
| **AA0395** | 10.25378/janelia.7614104 |
| **AA0396** | 10.25378/janelia.7614107 |

| | |
|---|---|
| **AA0397** | 10.25378/janelia.7614113 |
| **AA0398** | 10.25378/janelia.7614116 |
| **AA0400** | 10.25378/janelia.7614122 |
| **AA0401** | 10.25378/janelia.7614125 |
| **AA0402** | 10.25378/janelia.7614128 |
| **AA0407** | 10.25378/janelia.7614158 |
| **AA0408** | 10.25378/janelia.7614173 |
| **AA0409** | 10.25378/janelia.7614182 |
| **AA0410** | 10.25378/janelia.7614191 |
| **AA0411** | 10.25378/janelia.7614194 |
| **AA0412** | 10.25378/janelia.7614200 |
| **AA0413** | 10.25378/janelia.7614203 |
| **AA0415** | 10.25378/janelia.7614212 |
| **AA0416** | 10.25378/janelia.7614215 |
| **AA0418** | 10.25378/janelia.7614221 |
| **AA0419** | 10.25378/janelia.7614227 |
| **AA0421** | 10.25378/janelia.7614233 |
| **AA0422** | 10.25378/janelia.7614236 |
| **AA0424** | 10.25378/janelia.7614242 |
| **AA0426** | 10.25378/janelia.7614251 |
| **AA0439** | 10.25378/janelia.7614329 |
| **AA0440** | 10.25378/janelia.7614341 |
| **AA0441** | 10.25378/janelia.7614356 |
| **AA0442** | 10.25378/janelia.7614368 |
| **AA0445** | 10.25378/janelia.7614596 |
| **AA0446** | 10.25378/janelia.7614707 |
| **AA0450** | 10.25378/janelia.7614965 |
| **AA0452** | 10.25378/janelia.7615274 |

| | |
|---|---|
| **AA0460** | 10.25378/janelia.7615835 |
| **AA0461** | 10.25378/janelia.7615838 |
| **AA0462** | 10.25378/janelia.7615862 |
| **AA0463** | 10.25378/janelia.7615868 |
| **AA0464** | 10.25378/janelia.7615880 |
| **AA0465** | 10.25378/janelia.7615889 |
| **AA0466** | 10.25378/janelia.7615895 |
| **AA0467** | 10.25378/janelia.7615901 |
| **AA0469** | 10.25378/janelia.7615934 |
| **AA0470** | 10.25378/janelia.7615940 |
| **AA0471** | 10.25378/janelia.7615952 |
| **AA0472** | 10.25378/janelia.7615958 |
| **AA0473** | 10.25378/janelia.7615961 |
| **AA0474** | 10.25378/janelia.7615964 |
| **AA0475** | 10.25378/janelia.7615967 |
| **AA0481** | 10.25378/janelia.7615991 |
| **AA0534** | 10.25378/janelia.7640063 |
| **AA0541** | 10.25378/janelia.7640162 |
| **AA0543** | 10.25378/janelia.7640168 |
| **AA0544** | 10.25378/janelia.7640171 |
| **AA0545** | 10.25378/janelia.7640174 |
| **AA0548** | 10.25378/janelia.7640198 |
| **AA0549** | 10.25378/janelia.7640201 |
| **AA0553** | 10.25378/janelia.7640216 |
| **AA0554** | 10.25378/janelia.7640219 |
| **AA0556** | 10.25378/janelia.7640234 |
| **AA0576** | 10.25378/janelia.7649849 |
| **AA0577** | 10.25378/janelia.7649852 |

| | |
|---|---|
| **AA0578** | 10.25378/janelia.7649858 |
| **AA0582** | 10.25378/janelia.7649873 |
| **AA0596** | 10.25378/janelia.7649984 |
| **AA0599** | 10.25378/janelia.7650020 |
| **AA0600** | 10.25378/janelia.7650029 |
| **AA0602** | 10.25378/janelia.7650038 |
| **AA0624** | 10.25378/janelia.7655762 |
| **AA0625** | 10.25378/janelia.7655765 |
| **AA0626** | 10.25378/janelia.7655768 |
| **AA0627** | 10.25378/janelia.7655771 |
| **AA0628** | 10.25378/janelia.7655780 |
| **AA0632** | 10.25378/janelia.7658054 |
| **AA0633** | 10.25378/janelia.7658057 |
| **AA0637** | 10.25378/janelia.7658075 |
| **AA0639** | 10.25378/janelia.7658081 |
| **AA0641** | 10.25378/janelia.7658087 |
| **AA0646** | 10.25378/janelia.7658126 |
| **AA0650** | 10.25378/janelia.7658144 |
| **AA0653** | 10.25378/janelia.7658153 |
| **AA0654** | 10.25378/janelia.7658156 |
| **AA0656** | 10.25378/janelia.7658219 |
| **AA0659** | 10.25378/janelia.7658231 |
| **AA0663** | 10.25378/janelia.7658246 |
| **AA0666** | 10.25378/janelia.7658255 |
| **AA0668** | 10.25378/janelia.7658261 |
| **AA0669** | 10.25378/janelia.7658264 |
| **AA0670** | 10.25378/janelia.7704197 |
| **AA0671** | 10.25378/janelia.7704200 |

| | |
|---|---|
| **AA0672** | 10.25378/janelia.7704203 |
| **AA0726** | 10.25378/janelia.7707194 |
| **AA0727** | 10.25378/janelia.7707206 |
| **AA0733** | 10.25378/janelia.7707284 |
| **AA0734** | 10.25378/janelia.7707296 |
| **AA0735** | 10.25378/janelia.7707302 |
| **AA0738** | 10.25378/janelia.7707323 |
| **AA0739** | 10.25378/janelia.7707326 |
| **AA0741** | 10.25378/janelia.7707338 |
| **AA0744** | 10.25378/janelia.7707356 |
| **AA0745** | 10.25378/janelia.7707359 |
| **AA0746** | 10.25378/janelia.7707365 |
| **AA0747** | 10.25378/janelia.7707368 |
| **AA0748** | 10.25378/janelia.7707371 |
| **AA0749** | 10.25378/janelia.7707374 |
| **AA0767** | 10.25378/janelia.7710077 |
| **AA0772** | 10.25378/janelia.7710107 |
| **AA0773** | 10.25378/janelia.7710113 |
| **AA0774** | 10.25378/janelia.7710116 |
| **AA0780** | 10.25378/janelia.7739285 |
| **AA0781** | 10.25378/janelia.7739288 |
| **AA0782** | 10.25378/janelia.7739303 |
| **AA0783** | 10.25378/janelia.7739312 |
| **AA0784** | 10.25378/janelia.7739321 |
| **AA0785** | 10.25378/janelia.7739324 |
| **AA0786** | 10.25378/janelia.7739330 |
| **AA0787** | 10.25378/janelia.7739366 |
| **AA0788** | 10.25378/janelia.7739369 |

| | |
|---|---|
| **AA0789** | 10.25378/janelia.7739375 |
| **AA0802** | 10.25378/janelia.7739555 |
| **AA0803** | 10.25378/janelia.7739558 |
| **AA0817** | 10.25378/janelia.7739717 |
| **AA0836** | 10.25378/janelia.7739879 |
| **AA0837** | 10.25378/janelia.7739888 |
| **AA0840** | 10.25378/janelia.7739903 |
| **AA0853** | 10.25378/janelia.7739954 |
| **AA0854** | 10.25378/janelia.7739957 |
| **AA0859** | 10.25378/janelia.7740050 |
| **AA0865** | 10.25378/janelia.7740089 |
| **AA0866** | 10.25378/janelia.7740092 |
| **AA0880** | 10.25378/janelia.7742777 |
| **AA0882** | 10.25378/janelia.7742783 |
| **AA0884** | 10.25378/janelia.7742789 |
| **AA0887** | 10.25378/janelia.7742807 |
| **AA0888** | 10.25378/janelia.7742810 |
| **AA0889** | 10.25378/janelia.7742816 |
| **AA0897** | 10.25378/janelia.7780811 |
| **AA0898** | 10.25378/janelia.7780814 |
| **AA0899** | 10.25378/janelia.7780820 |
| **AA0900** | 10.25378/janelia.7780826 |
| **AA0905** | 10.25378/janelia.7780856 |
| **AA0907** | 10.25378/janelia.7780862 |
| **AA0908** | 10.25378/janelia.7780865 |
| **AA0911** | 10.25378/janelia.7780874 |
| **AA0914** | 10.25378/janelia.7780883 |
| **AA0916** | 10.25378/janelia.7780898 |

| | |
|---|---|
| **AA1096** | n/a |
| **AA1105** | n/a |
| **AA1111** | n/a |
| **AA1121** | n/a |
| **AA1124** | n/a |
| **AA1131** | n/a |
| **AA1144** | n/a |
| **AA1145** | n/a |
| **AA1156** | n/a |
| **AA1167** | n/a |
| **AA1169** | n/a |
| **AA1190** | n/a |
| **AA1240** | n/a |
| **AA1476** | n/a |
| **AA1481** | n/a |
| **AA1494** | n/a |
| **AA1537** | n/a |
| **AA1540** | n/a |
| **AA1541** | n/a |
| **AA1543** | n/a |
